# Phylotranscriptomics Allows Distinguishing Major Gene Flow Events from Incomplete Lineage Sorting in Rapidly Diversifying Mimetic Orchids (Genus *Ophrys*)

**DOI:** 10.64898/2026.01.07.698074

**Authors:** Lucas Vandenabeele, Anaïs Gibert, Pascaline Salvado, Roselyne Buscail, Christel Llauro, Michèle Laudie, Hervé Philippe, Joris A. M. Bertrand

## Abstract

*Ophrys* orchids (or bee orchids) provide an outstanding example of a plant adaptive radiation. Over the last five million years, this genus has diversified into hundreds of taxa as a result of its unconventional pollination strategy, known as ‘sexual swindling’. However, the rapid and substantial diversification of this genus, combined with its capacity for hybridisation and large genome size, poses significant challenges in addressing its systematics. We used phylotranscriptomics as a genome complexity reduction technique to infer the phylogenetic relationships among *Ophrys* main lineages. More than seven thousand gene trees enabled us to determine the relative contributions of gene flow and incomplete lineage sorting (ILS) in *Ophrys* evolution. First, we propose a new phylogenetic hypothesis for the genus with an unprecedented resolution that largely confirms the relationships between the main *Ophrys* lineages, but also provides new insights within each sub-genera. By combining phylogenetic network inference with introgression analyses based on gene tree topologies and branch lengths, we then show that the numerous phylogenetic incongruences among gene tree topologies result from a pervasive background of ILS, over which stand out several well-supported, ancient and potentially adaptive gene flow events between lineages. These major gene flow events provide a new perspective on the evolution of the *Ophrys* genus and its pollination, questioning previous hypotheses inferred without considering its reticulate evolution, and providing a better understanding of discrepancies observed among previous phylogenetic studies of the genus.

## Introduction

The systematics and taxonomy of groups that exhibit high diversification rates are challenging. These groups not only present a scarce phylogenetic signal, but are also particularly prone to incomplete lineage sorting (ILS), whereby ancestral genetic variation has not been sorted into distinct lineages, and reticulate evolution, whereby gene flow can occur between emerging lineages. Although gene flow has long been considered an obstacle to diversification (Mayr, 1942), empirical studies have repeatedly demonstrated its importance in the speciation and diversification of various groups of organisms (see Suarez-Gonzalez et al., 2018, for several examples). Various hypotheses have been put forward to explain how hybridisation promotes diversification during evolutionary radiations (see Combrink et al., 2025). Both ILS and gene flow can affect the recovery of the tree-like (or vertical) phylogenetic signal, and these phenomena are not mutually exclusive (Degnan and Rosenberg, 2009). In groups undergoing radiation, where the vertical (speciation) phylogenetic signal is already weak, the combined presence of ILS and horizontal signal from gene flow can further obscure the true species relationships (or species tree), resulting in topological incongruences between individual gene trees and the species tree. In extreme cases, the horizontal signal may even dominate genome-wide patterns (Fontaine et al., 2015). Thus, it has become evident that an increased number of genetic markers is necessary to accurately infer the evolutionary history of investigated groups (Maddison and Knowles, 2006; Brito and Edwards, 2009). Phylogenetic hypotheses derived from a single *locus*, or a modest number of *loci*, often lack sufficient information. In plants, for example, the conventional genetic markers (*e.g.* nrITS, matK or rbcl), which have been frequently employed to reconstruct molecular phylogenies, are inadequate for estimating the effects of ILS and gene flow.

The advent of high-throughput sequencing technologies has enabled the use of hundreds to thousands of *loci*, or even entire genomes, to gain resolution and compare a substantial number of individual gene trees. In some groups however, genome size continues to constrain the (re)sequencing of a large number of individuals. In such cases, several complexity reduction techniques exist that facilitate sub-sampling of the genome. This is the case of phylotranscriptomics (Cheon et al., 2021), which focuses on capturing genes expressed in specific tissues by sequencing the transcriptome (here, mRNAs). The sequences obtained include the exons of coding genes, as well as untranslated regions (5’ and 3’UTR). Other alternative methods exist, such as RAD seq-like approaches (*e.g.* Baird et al., 2008; Peterson et al., 2012), whereby orthologous genomic regions in close proximity to restriction enzyme cut sites are consistently sequenced in a set of individuals. Another example is the use of bait approaches, which are specifically designed to target hundreds of *loci*. In plants, for example, recent publications have introduced ‘universal’ kits, such as the Angiosperms353 kit (Johnson et al., 2019), as well as more specialised sets of probes that hybridise to specific genomic regions of particular families, such as the Orchidaceae963 (Eserman et al., 2021) or Orchidinae-205 kit (Veltman et al., 2024) for orchids. Compared to these two approaches, phylotranscriptomics generally gives a greater amount of information, although it focuses on transcribed regions and does not provide information on highly variable intronic regions. The transcriptome provides access to the thousands of genomic sequences transcribed in the studied tissue (typically more information than with bait kits), which are longer (once assembled) than with the two aforementioned approaches. Furthermore, these sequences are likely to contain some genes functionally relevant to studying the link between genetic and phenotypic variation. Like the other alternatives, phylotranscriptomics is also not devoid of difficulty to infer orthologous relationships (Yang and Smith, 2013) due to the presence of multiple transcriptional isoforms (resulting from alternative splicing) and assembly artefacts. Nevertheless, phylotranscriptomics has already been successfully employed to resolve the phylogeny of several groups (Cheon et al., 2020) and to study reticulate evolution (Rancilhac et al., 2021).

Phylotranscriptomics may thus be an effective method for investigating the systematics and taxonomy of a particularly challenging group: the *Ophrys* genus, a representative of the highly diverse orchid family with a large genome (haploid genome size of 5 to 7 Gb, see Bou Dagher-Kharrat et al., 2013; Abreu et al., 2017; Gibert et al., 2025; Salvado et al., 2025). All *Ophrys* species employ an unconventional pollination strategy called ‘sexual swindling’ to ensure their reproduction (reviewed in Baguette et al., 2020). The flowers mimic the appearance and the scent (through the emission of pseudo-pheromones) of female insects, particularly hymenopterans, and deceive conspecific males attempting to copulate with the flower, potentially enabling pollen transfer between plants. This pollination strategy is specific, with each *Ophrys* species being pollinated by only a few or even a single species of insect (Joffard et al., 2019). It has been proposed that the high degree of specificity induced by the selective pressures exerted by male insect pollinators on flower phenotypes is the driver of *Ophrys* evolutionary diversification (Baguette et al., 2020). The *Ophrys* genus indeed provides an excellent illustration of adaptive radiation in plants. This genus emerged less than five million years ago, and has since diversified into tens to hundreds of taxa (Breitkopf et al., 2015). In the absence of clear evidence of post-zygotic barriers between the majority of its representatives, it can be assumed that pre-zygotic isolation induced by the pollinators is primarily responsible for the formation and the persistence of *Ophrys* lineages. It should be noted that the majority of these lineages remain interfertile and occasionally form hybrids in the wild (Scopece et al., 2007). The *Ophrys* genus is thus particularly susceptible to reticulate evolution, which may explain the lack of consensus concerning its systematics and taxonomy.

Depending on the authors and criteria, the *Ophrys* genus comprises between 9 and ca. 350 species (Bateman et al., 2018; Delforge, 2016). Most molecular phylogenetic hypotheses published to date generally agree on the existence of three major subgenera, which are themselves subdivided into 9 or 10 macro- or flagship species. Studies based on traditional genetic markers, such as nrITS, sometimes coupled with plastid *loci* (Soliva et al., 2001; Devey et al., 2008; Tyteca and Baguette, 2017), a handful of low-copy nuclear genes (Breitkopf et al., 2015), RADseq (Bateman et al., 2018) or whole-plastid genomes (Bateman and Rudall, 2023) have failed to delineate lineages at lower taxonomic levels (but see Sedeek et al., 2014; Gibert et al., 2025; Salvado et al., 2025, for counter-examples using RADseq-like methods). Moreover, these studies sometimes report discordant tree topologies. For example, several studies have first inferred a basal position for the *O. insectifera* clade (Devey et al., 2008; Breitkopf et al., 2015), a placement not further supported by different types of ‘-omic’ data: namely RAD-seq (Bateman et al., 2018), transcriptomes (Piñeiro Fernández et al., 2019) and whole-plastid genomes (Bertrand et al., 2021b).

In a nutshell, *Ophrys* consists of 9 lineages: *insectifera* (A), *tenthredinifera* (B), *speculum* (C), *bombyliflora* (D), *fusca* (E), *apifera* (F), *sphegodes* (G), the *fuciflora* complex (H’, which corresponds to the fusion of the former *scolopax* (I) and *fuciflora* (H) lineages) and *umbilicata* (J) (also referred as clade A to J by Devey et al., 2008; Bateman et al., 2018), which are well characterized by both morphological and genetic data but whose phylogenetic relationships remain unclear and have never been explored based on genomic data, out of the notable exception of Bateman et al. (2018), who rely on ca. 4000 SNPs derived from a RAD-seq approach. In this context, phylotranscriptomics could clarify *Ophrys* systematics based on an unprecedented number of *loci*, and establish whether, and if so to which extent, the evolution of the genus *Ophrys* is reticulated. In this study, we considered floral transcriptomes of 8 out these (9-10) lineages to infer the first near exhaustive phylotranscriptomic hypothesis for the *Ophrys* genus. Beyond systematics and taxonomic considerations, our main objective is to investigate the contribution of gene flow to *Ophrys* evolution. By comparing multiple gene tree topologies and branch lengths with different methods, ranging from ABBA-BABA tests to approaches that explicitly model reticulate evolution (such as phylogenetic networks), we first aim to test for the existence of ‘major’ events of gene flow that could have shaped the history of *Ophrys* diversification. In doing so, our objective is not to explain all topological incongruences, but to determine whether and which fraction of them results from gene flow rather than from incomplete lineage sorting (ILS), the latter representing a systematic source of conflict considered here as a confounding factor. We then aim at further exploring the genomic patterns associated with gene flow, namely whether genes associated with gene flow events are scattered across the genomes or form more localised clusters. Finally, an investigation of candidate genes known to be associated with *Ophrys* flower phenotypes aims at testing the potential adaptive role, or at least the phenotypic relevance, of inferred gene flow in *Ophrys* evolution.

## Materials and Methods

### Taxon Sampling

The floral transcriptomes of one species from each of the main *Ophrys* lineages (Table 1), with the exception of *umbilicata* (J) lineage, and two species from the closely related genus *Himantoglossum* (*H. robertianum* and *H. hircinum*), were newly sequenced. Unpollinated flowers from these species were sampled in the wild in southern France, and then immersed in an RNA-stabilising buffer (RNAlater) and stored at -20°C until RNA extraction. *Ophrys* RNA extraction was performed using Monarch brand kits according to the manufacturer’s protocol. Libraries were prepared and sequenced on a NextSeq500 sequencer (Illumina) in paired-end mode (2 x 150 bp) at the BioEnvironnement platform of the University of Perpignan Via Domitia. The two *Himantoglossum* were sequenced in a subsequent run with similar settings. A permit (n°2018-s-20) was obtained from the DREAL of Occitanie (Direction Régionale de l’Environnement et de l’Aménagement du Territoire) to collect samples in France, where several *Ophrys* species are legally protected.

**Table 1.** Orchid species sampled for floral transcriptome sequencing.

| Species | Lineage | Locality | Accession number |
| --- | --- | --- | --- |
| *Ophrys insectifera* | *insectifera* (A) | Versols-Et-Lapeyre, France | ERS28372667 |
| *Ophrys tenthredinifera* | *tenthredinifera* (B) | Saint-Paul de Fenouillet, France | ERS28372672 |
| *Ophrys bombyliflora* | *bombyliflora* (C) | Narbonne, France | ERS28372664 |
| *Ophrys speculum* | *speculum* (D) | Rivesaltes, France | ERS28372669 |
| *Ophrys lutea* | *fusca* (E) | Rivesaltes, France | ERS28372668 |
| *Ophrys apifera* | *apifera* (F) | Villeneuve de la Raho, France | ERS28372673 |
| *Ophrys exaltata* | *sphagodes* (G) | Torreilles, France | ERS28372666 |
| *Ophrys scolopax* | *fuciflora* complex (H') | Rivesaltes, France | ERS28372671 |
| *Himantoglossum hircinum* | outgroup | Versols-Et-Lapeyre, France | ERS28372676 |
| *Himantoglossum robertianum* | outgroup | Torreilles, France | ERS28372675 |

Collector: Joris A. M. BERTRAND and R. BUSCAIL with contribution of A. GIBERT

### Transcriptome Assembly

The quality of Illumina reads was initially assessed with FastQC v0.11.9 (https://www.bioinformatics.babraham.ac.uk/projects/fastqc). Adapter and low-quality reads were filtered out using Trimmomatic v0.39 (Bolger et al., 2014). Filtered reads were then assembled *de novo* using three distinct software: Trinity v2.13.2 (Grabherr et al., 2011), Trans-Abyss v2.0.1 (Robertson et al., 2010), and rnaSPAdes v3.15.4 (Bushmanova et al., 2019), with 5 different k-mer sizes for Trans-Abyss and rnaSPAdes (21, 39, 59, 79, and 99). The resulting assemblies were then merged using Trans-Abyss-Merge v2.0.1. Redundancy was evaluated with EvigeneR (Gilbert, 2013), and only the isoforms with the longest CDS were retained. The completeness of the transcriptome assemblies was then assessed using BUSCO v5.6.1 (Manni et al., 2021) against the Liliopsida_odb10 database.

### Nuclear Orthologous Genes Identification

Single-copy orthologous genes were inferred from *de novo* assembled transcriptomes using a pipeline based on both sequence similarity and phylogenetic tree topology (summarised in Supplementary Fig. S1). Homologous gene clusters were inferred using OrthoFinder v2.5.4 (Emms and Kelly, 2019). Only gene clusters containing sequences for all 8 *Ophrys* species and at least one of the two outgroups were retained. Sequences whose length was less than 300 bp were removed. For each gene cluster, nucleotide sequences were aligned using MAFFT v7.490 (Katoh and Standley, 2013) with the L-INS-i option, and low-similarity fragments were removed using Hmmcleaner v0.180750 (Di Franco et al., 2019). Gene alignments were filtered to retain only single-copy orthologous genes. Distinct ancient paralogous groups were divided into two different clusters. Chimeric sequences were constructed from inparalogs using SCaFoS v1.25 (Roure et al., 2007).

A first concatenated sequence was generated from all orthologous gene alignments using SCaFoS. Alignments containing sequences excessively divergent compared to expectations based on this concatenation were removed using the Branch Length Comparison method (Simion et al., 2020) (see section ‘Orthologous gene alignments filtering details’ in Supplementary Methods for details). To exclude putative mitochondrial and plastid orthologs, BLASTx searches (Camacho et al., 2009) were performed on our gene alignments using the available *Ophrys* chloroplast genomes (Roma et al., 2018; Bertrand et al., 2021a) and *Dendrobium nobile* mitochondrial genomes, as no mitochondrial genome is available for *Ophrys* or closely related genera. For alignments containing sequences from both outgroup species (genus *Himantoglossum*), a chimeric sequence was constructed from the two sequences using SCaFoS (*Himantoglossum* species were monophyletic in all genes where both sequences were present). This chimeric sequence was then used as outgroup for subsequent analyses.

### Plastid Genome Reconstruction

For each individual, plastid genome sequence was reconstructed by first mapping the filtered reads onto the *Ophrys lutea* plastid genome (GenBank accession: NC_058525.1) (Bertrand et al., 2021a) using STAR v2.7.10 (Dobin et al., 2013). Variant calling was then performed with bcftools v1.22 (Li, 2011) in haploid mode, and SNPs were filtered based on quality (QUAL ≥ 30) and read depth (DP ≥ 10). For each individual, a consensus sequence was reconstructed by incorporating the filtered SNPs into the reference genome, with positions lacking coverage replaced by missing data (‘N’). As all the consensus sequences had the same size (the one of the *O. lutea* plastid genome), no further alignment steps were required. Positions consisting of only missing data were discarded. Regions corresponding to ndh subunits (ndhA to ndhK) were also removed from the alignment, as these genes are likely to be pseudogenised in *Ophrys* plastomes because their functional copies have been transferred to the nuclear genome (Bertrand et al., 2021a), as in many other orchids (Lin et al., 2015). The plastid phylogeny was then inferred from this plastid transcriptome alignment using IQ-TREE 2 v2.3.6 (Minh et al., 2020), with the ‘model selection’ option and 500 standard nonparametric bootstrap replicates.

### Gene and Species Trees Reconstruction

A gene tree was inferred for each single-copy orthologous gene using IQ-TREE 2 with the ‘model selection’ option and 500 standard non-parametric bootstrap replicates. Two methods were employed to infer species trees: a coalescent-based method using ASTRAL-III v5.7.8 (Zhang et al., 2018) based on these single-copy orthologous gene trees; and a concatenation method using IQ-TREE 2 with the aforementioned options, based on the concatenated sequence of the single-copy orthologous genes obtained with SCaFoS. Gene trees contain several bipartitions (*i.e.* nodes). For each bipartition, the frequency of gene trees supporting the given partition was calculated using Phyparts (Smith et al., 2015). The resulting frequencies were visualised as pie charts in the two species trees using the Python script ‘phypartspiecharts.py’ (github.com/mossmatters/MJPythonNotebooks). We then inferred consensus split networks (Holland et al., 2004) using SplitsTree v6.4.12 (Huson and Bryant, 2006) and our single-copy orthologous gene trees. Consensus split networks display discrepancies and ambiguous signals in a set of trees by showing all the ‘splits’ (*i.e.* all the bipartitions of the taxa) that are present in at least a specified proportion of the input tree set. We used two different proportions: 15% and 20%. Each ‘split’ was here weighted by the number of trees supporting it (‘count’ weighting method).

### Gene Flow Inference

To detect interspecific gene flow, we employed three different phylogenetic approaches, relying on single-copy orthologous gene trees.

First, we inferred phylogenetic networks using PhyloNet (Than et al., 2008) following the maximum pseudo-likelihood algorithm (Yu and Nakhleh, 2015). This approach relies exclusively on gene tree topologies and enables the simultaneous estimation of both the species tree and potential reticulation events. Each reticulation is associated with an inheritance probability, representing the proportion of genetic material inherited from each parental lineage. PhyloNet was used with an increasing number of reticulations ‘*h*’, ranging from 0 to 6, with 3 replicates of 100 independent iterations per *h* value. The branch lengths and inheritance probabilities of each proposed phylogenetic network were then optimised to compute its pseudo-likelihood score. To account for phylogenetic uncertainty, PhyloNet was also run after collapsing nodes with *<* 70% bootstrap support in the gene trees. An alternative tool to infer phylogenetic networks by maximum pseudo-likelihood is SNaQ (Solís-Lemus and Ané, 2016; Solís-Lemus et al., 2017). Unlike PhyloNet, SNaQ does not use gene trees directly, but instead calculates quartet concordance factors (CFs) from gene trees and uses these CFs to infer phylogenetic networks. It should be noted that SNaQ is limited to level 1 networks and is therefore unable to infer more than one reticulation event involving a given taxon, unlike PhyloNet. SNaQ was run with the coalescent-based tree topology as starting tree, and with an increasing number of reticulations ‘*h*’ ranging from 0 to 6, with 100 iterations per *h* value.

The slope heuristic approach described in Solís-Lemus and Ané (2016) was used to identify the ‘best’ number of reticulations for both tools. For each number of reticulations, the pseudo-likelihood score (or negative log-pseudo-likelihood for SNaQ) of the five best networks was plotted as a function of the number of reticulations. The last number of reticulations inducing a sharp increase of the network pseudo-likelihood score was determined to be the ‘best fitting number of reticulations’. To assess reticulation support in SNaQ best networks, we used bootSNaQ from the PhyloNetworks Julia package (Solís-Lemus et al., 2017) with 100 bootstrap replicates. Each of these bootstrap replicates recalculates the CFs using IQ-TREE’s bootstrap gene trees. A network is then inferred from these CFs, with the ‘best’ number of reticulations and 10 iterations. The best bootstrapped networks were then visualised using PhyloPlots from PhyloNetworks. For the PhyloNet results, the 15 best networks from the 3 replicates with the ‘best’ number of reticulations were visualised using Dendroscope 3 (Huson and Scornavacca, 2012). Reticulations and backbone tree of these networks were then compared using SummarizeNetworks from PhyloNet. A backbone tree is a tree obtained by removing all the minor branches (*i.e.* branches with inheritance probabilities of less than 0.5 in a reticulation) of each reticulation in a network.

We then employed the Δ test (Huson et al., 2005), which uses the same logic as the widely used Patterson’s D statistic (also known as the ‘ABBA/BABA’ test (Green et al., 2010; Durand et al., 2011; Patterson et al., 2012)), but applied to gene tree topologies rather than single-nucleotide polymorphisms (SNPs). This method uses rooted quartets, a subset of four taxa consisting of an inner triplet (*P*_1_, *P*_2_, *P*_3_) and an outgroup (*O*), which have three possible topologies: a ‘matching topology’, corresponding to the species tree topology ((*P*_1_,*P*_2_),*P*_3_), which is expected to be in a major proportion, and two discordant topologies, ((*P*_1_,*P*_3_),*P*_2_) and ((*P*_2_,*P*_3_),*P*_1_). This test compares the proportions of *loci* that support each of the two discordant topologies, and assumes a null hypothesis of ILS only, with the expectation that the two discordant topologies will be in equal proportion. A significant deviation from this expected equality is interpreted as a sign of gene flow. This test was conducted using a R function developed by Rancilhac et al. (2021), here with *Himantoglossum* used as the outgroup and all possible combinations of three *Ophrys* taxa as the inner triplet. The function assessed the significance of the departure from expectation under ILS using 1000 bootstrap replicates. The *p*-values were corrected for multiple testing using the Benjamini–Hochberg method. For each significant Δ statistic (*p*-value *<* 0.01), the proportion of introgressed genes *γ* was calculated as outlined in Uckele et al. (2024). This metric is an adaptation of the f4-ratio (Patterson et al., 2012), initially developed for SNPs data, to gene trees. To better interpret this result, we used Dsuite v0.5r50 (Malinsky et al., 2021) to calculate and visualise the ‘f-branch’ statistic (Malinsky et al., 2018) from the introgressed gene proportion *γ*. This ‘f-branch’ statistic detects introgression events between tips and internal branches in a species tree. The Δ statistic was also calculated after collapsing nodes with *<* 70% bootstrap support in the gene trees, in order to account for phylogenetic uncertainty.

In addition to all previous methods that rely solely on tree topologies to infer introgression events, we also conducted introgression tests based on gene tree branch lengths as a proxy for coalescence times, in order to detect introgression and to estimate the amount of ILS. We used Aphid v0.11 Galtier (2024), an approximate maximum likelihood method that quantifies sources of phylogenetic conflict by analysing the topology and branch lengths of rooted quartets. For each gene tree, Aphid calculates the probability that it has been affected by ILS, gene flow, or neither of these events. In a triplet, branch lengths separating the two sister taxa are expected to be shorter than average if gene flow has occurred, (as gene flow takes place after speciation), and longer than average if ILS has occurred (as ILS results from a failure of coalescence in the ancestor of the two sister taxa). Aphid was applied to each rooted quartet used for the Δ test (formed by an outgroup and an inner triplet). Before triplet analysis, Aphid performs a preprocessing step that filters gene trees: genes with mutation rate estimates outside the default thresholds implemented in Aphid are excluded. As suggested in Galtier (2024), the significance of gene flow between non-sister taxa was assessed by recalculating the likelihood for each quartet under the constraint that the probability of gene flow equals zero. The two resulting likelihoods, with and without gene flow, were then compared using a likelihood ratio test. Aphid also estimates the ILS imbalance (*I*_ILS_), which is the difference in the amount of ILS between the two discordant topologies. We expect the ILS amount to be balanced, with an equal amount of ILS in both discordant topologies (*I*_ILS_ = 0.5). An *I*_ILS_ value significantly different from 0.5 suggests that Aphid may have misidentified the origin of some genes Galtier (2024), indicating either an Aphid failure or the existence of ghost gene flow (*i.e.* gene flow from an unsampled or extinct taxon). For each triplet, we then tested whether the *I*_ILS_ value significantly departed from 0.5 using 1000 bootstrap replicates. The *p*-values were corrected for multiple testing using the Benjamini–Hochberg method. For each quartet, Aphid estimates the timing of gene flow relative to the speciation of the inner triplet’s sister taxa using the statistic *p_a_* (probability of ancient gene flow). By default, Aphid considers two possible timings: one coinciding with the speciation event, and one that is approximately twice as recent. The parameter *p_a_* corresponds to the probability that gene flow occurred close to the time of speciation. To refine the gene flow dating, Aphid was rerun with ten possible timings instead of two, ranging from the time of speciation to ten times more recently.

To facilitate the estimation of species divergence times, we first used Sortadate (Smith et al., 2018) to identify genes following a clock-like evolution pattern (or ‘clock-like genes’) within the dataset. Selection was based on three criteria, in order of importance: bipartition support, root-to-tip variance, and tree length. Bipartition support was assessed by comparing the topology of the gene trees with the topology of a reference species tree. This enabled the selection of 1,000 clock-like genes from the dataset. These genes were then used to perform a species divergence time estimation using the Bayesian tool MCMCTree from the Paml package v4.10.6 (Yang, 2007). We calibrated the root of our species trees using divergence time estimates between *Himantoglossum* and *Ophrys* found by Inda et al. (2012) (9.1–16.5 million years ago (Ma)).

### Gene Annotation and Genomic Position

Orthologous gene alignments were mapped to the first and so far, only available reference genome for *Ophrys* (the one of *Ophrys sphegodes*, Russo et al. (2024)) using Minimap2 v2.24 (Li, 2018, 2021) to determine their genomic positions and annotations. Genes that could not be mapped with Minimap2 were subsequently searched against the reference genome using a combination of Exonerate v2.2.0 (Slater and Birney, 2005) and BLASTX v2.13.0 (Camacho et al., 2009). The 396 genes showing non-overlapping or conflicting mapping results between Exonerate and BLASTX were excluded from this analysis. Based on this gene set, a relative gene density for each chromosome was then evaluated by counting the number of genes mapped in consecutive 1-Mb windows. For the three major gene flow events, we identified putatively introgressed genes using gene tree topology. Gene trees in which two non-sister species, with evidence of gene flow, formed a monophyletic group were considered as potentially subject to introgression.

We then used twisst (Martin and Van Belleghem, 2017) to visualise the position of these candidate genes along *Ophrys* reference genome chromosomes. We also decided to represent the distribution of topologies along ‘pseudochromosomes’ (*i.e.* here, only consisting of the concatenation of the genes considered in this study), as our genes did not cover the entire chromosomes, resulting in a substantial amount of missing data. Within these pseudochromosomes, the relative position of our genes is conserved, but all regions of the chromosomes with missing data were removed and the relative size of the genes was determined by the length of their alignments.

To identify chromosomes with a significantly different proportion of putativly introgressed genes compared to the others, we performed a *χ*^2^ test on each chromosome against all the others. The *p*-values were corrected for multiple testing using the Benjamini–Hochberg method.

Finally, to identify and evaluate the status of potentially phenotypically relevant genes in the *Ophrys* adaptive radiation in our orthologous gene alignments, we used a list of 296 *Ophrys* flower phenotype key genes (odour production, anthocyanin or carotenoid biosynthesis, fatty acid, wax and hydrocarbon biosynthesis, flower development …) localised on the *Ophrys* reference genome by Russo et al. (2024), Gibert et al. (2025) and Salvado et al. (2025). We then used our previous mapping onto the reference genome to determine whether any of our gene alignments corresponded to these key genes. Next, we evaluated if these genes exhibited gene flow evidence using both gene tree topology (as described in the previous paragraph) and Aphid results.

## Results

### Phylotranscriptomics Analyses

The final transcriptome assemblies contained over 80% of the expected BUSCO genes (Supplementary Table S1). From these assemblies, we identified 7,821 single-copy nuclear orthologous genes, with alignments ranging from 341 to 13,403 bp, totaling 14,979,757 bp and including 1,578,109 informative sites (1,281,756 without *Himantoglossum*) in the concatenated alignment.

Both the concatenation-based (Fig. 1) and coalescent-based (Supplementary Fig. S2) topologies were well supported, with all branches presenting bootstrap support of 100% and local posterior probabilities of 1, respectively. The two trees supported the existence of two main *Ophrys* clades, which we will refer to as clade 1 and clade 2 respectively. Clade 1 comprises the taxa *O. insectifera*, *O. apifera*, *O. scolopax* (*fuciflora* complex) and *O. exaltata* (*sphegodes*), and clade 2 the taxa *O. tenthredinifera*, *O. bombyliflora*, *O. lutea* (*fusca*) and *O. speculum*. However, the concatenation-based topology positioned *O. tenthredinifera* and *O. bombyliflora* as sister lineages, whereas the coalescent-based approach positioned *O. tenthredinifera* and *O. lutea* (*fusca*) as sister lineages. Phyparts (Fig. 1 and Supplementary Fig. S2) revealed substantial gene tree conflicts among clade 2, with only around a quarter of gene trees supporting the concordant topology. This result showed the presence of two alternative bipartitions in this clade, each of which being present in around 20% of the gene trees: one that groups *O. lutea* (*fusca*) and *O. tenthredinifera*, and the other that groups *O. bombyliflora* and *O. tenthredinifera*. This substantial amount of conflict, especially within clade 2, was corroborated by the split network (Supplementary Fig. S3).

**Figure 1.**
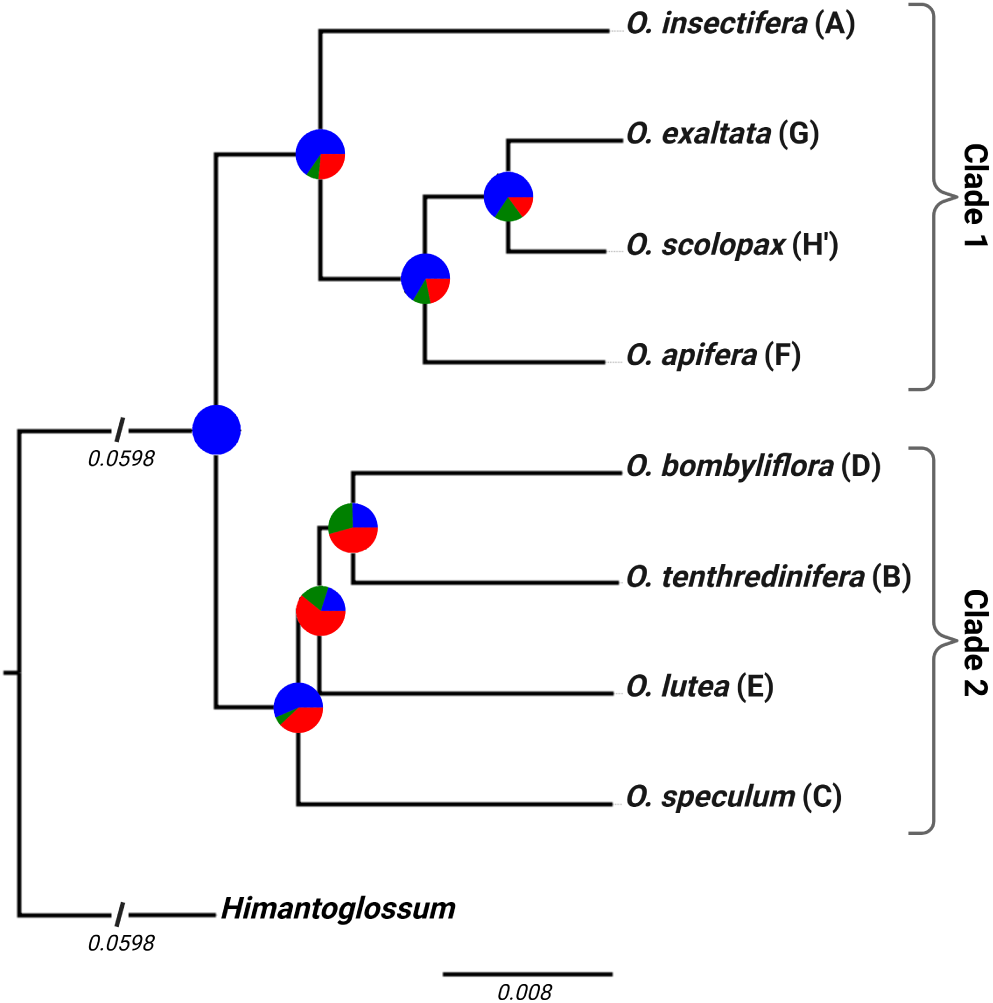
Maximum likelihood concatenation-based tree based on 7,821 orthologous gene alignments. All nodes have 100% bootstrap support. At each node, a pie chart depicts the proportion of gene trees supporting the concatenation-based tree node (blue), the main discordant alternative node (green), or other discordant alternative nodes (red). Tree was drawn with FigTree (http://tree.bio.ed.ac.uk/software/figtree/), and pie charts were added from ‘phypartspiecharts.py’ results with Biorender (https://BioRender.com/7gqnhty).

### Phylogenetic Networks Inference

Using the slope heuristic approach, we identified 4 as the best-fitting number of reticulations with PhyloNet (Fig. 2a). Among the 15 reconstructed networks with 4 reticulations, 14 had similar pseudo-likelihood scores, with a difference of less than 100 points between them (see these 14 networks on Supplementary Fig. S4). The best-scoring network (Fig. 2c) had four reticulations which were also present in most of the other 14 high-scoring networks. Alternate reticulations were also present, but in a minority of networks. When accounting for phylogenetic uncertainty by keeping only nodes with ≥ 70% bootstrap support, the best-fitting number of reticulations was also identified as 4 with PhyloNet (Fig. 2b), with 12 of the 15 best networks showing a similar pseudo-likelihood score (see these 12 networks on Supplementary Fig. S5). The three major reticulations of Fig. 2c were also found in the best-scoring network (Fig. 2d) and in the majority of the other 12 high-scoring networks, with similar inheritance probabilities. These three reticulation events consisted of:

- an important gene flow from *O. bombyliflora* and *O. lutea* (*fusca*) to *O. tenthredinifera*, with an average inheritance probability of 0.50 from *O. bombyliflora* and 0.50 from *O. lutea* (*fusca*) (0.42 and 0.58, respectively, when accounting for phylogenetic uncertainty).
- a gene flow from *O. apifera* to *O. scolopax* (*fuciflora* complex), with an average inheritance probability of 0.21 (0.28 when accounting for phylogenetic uncertainty);
- a gene flow from *O. tenthredinifera* to *O. apifera*, with an average inheritance probability of 0.07 (same value when accounting for phylogenetic uncertainty);

**Figure 2.**
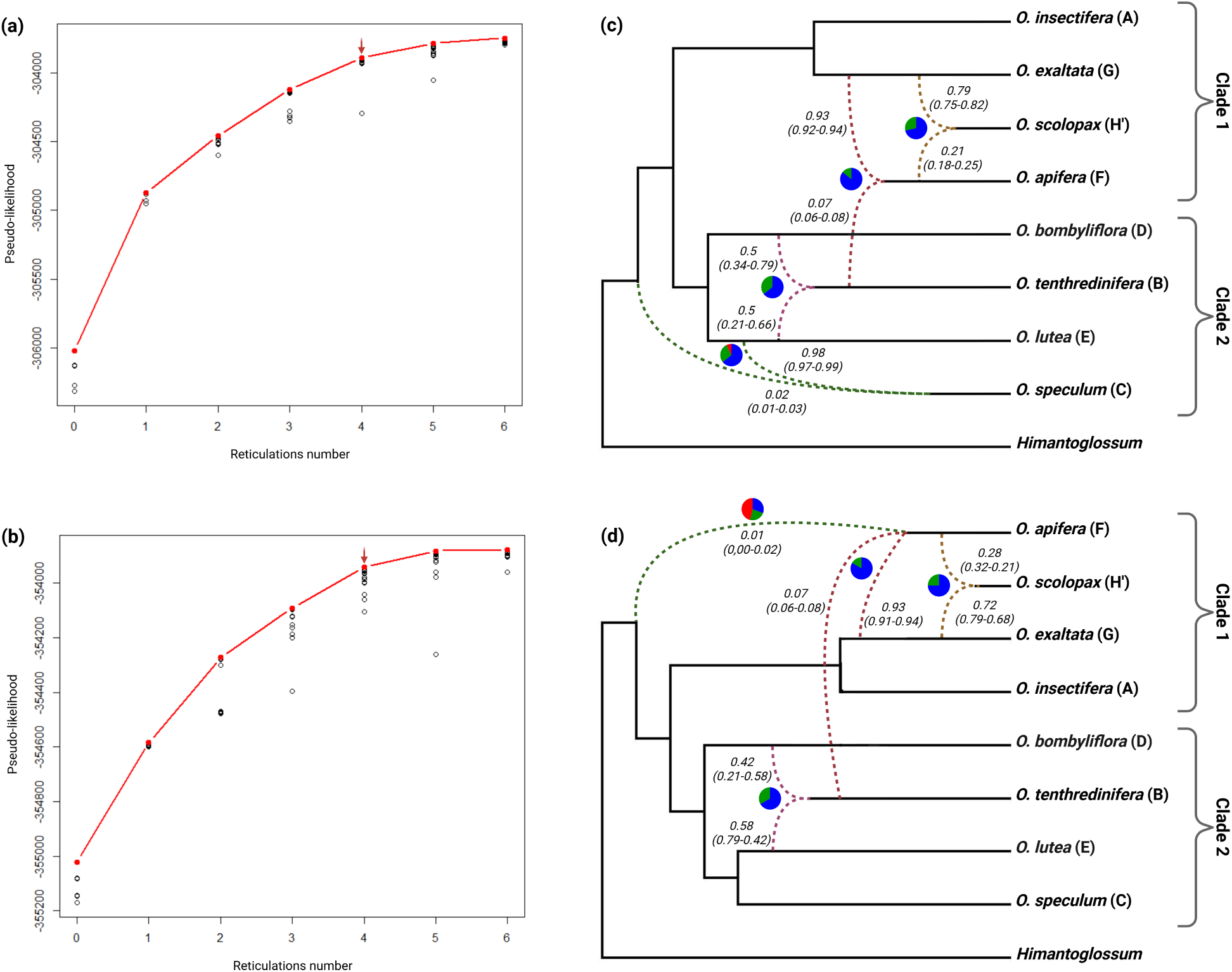
(a-b) Pseudo-likelihood scores of the 105 networks inferred with PhyloNet as a function of their number of reticulations, without accounting for phylogenetic uncertainty (a) and when keeping only nodes *≥* 70% bootstrap support (b). The red dot depicts the "best" network for each number of reticulations, and the red arrow indicates the ‘best’ number of reticulations according to the heuristic slope. (c-d) Average best PhyloNet networks with 4 reticulations according to pseudo-likelihood scores without accounting for phylogenetic uncertainty (c) and after keeping only nodes *≥* 70% bootstrap support (d). Dashed lines represent the reticulation events, and the associated numbers indicate the average inheritance probability rounded to the nearest hundredth of each reticulation (with min and max values associated). For each reticulation event, a pie chart represents the proportion of networks (among the 14 best networks (c) or 12 best networks (d)) supporting the reticulation (blue), the main alternative reticulation (green), and other alternatives (red). Networks were drawn with Dendroscope 3 (Huson and Scornavacca, 2012), and pie charts were added with Biorender (https://BioRender.com/rw1upon). Branch lengths are arbitrary.

A potential ghost introgression from the base of the *Ophrys* phylogeny was inferred in every best network, with an average inheritance probability of 0.02 (0.01 when accounting for phylogenetic uncertainty). In most phylogenetic networks inferred without accounting for phylogenetic uncertainty, the recipient of this gene flow was *O. speculum*. However, when accounting for phylogenetic uncertainty, it became unclear which species was the recipient of this gene flow (*O. insectifera*, *O. speculum*, *Himantoglossum*, *O. apifera*…, see Supplementary Fig. S5).

In brief, PhyloNet supported the presence of 3 consecutive reticulation events during *Ophrys* evolution, plus a potential ghost introgression. PhyloNet analyses also supported two alternative backbone trees, both of which differed from concatenation- and coalescent-based topologies :

- A ‘majority’ backbone tree (Supplementary Fig. S6a), supported by most of the best networks (9/14 without accounting for phylogenetic uncertainty, and 9/12 when accounting for it). In this tree, *O. bombyliflora* had a basal position in the clade 2 instead of *O. speculum*, and *O. tenthredinifera* was grouped with *O. lutea* (*fusca*).
- A ‘minority’ backbone tree (Supplementary Fig. S6b), supported by the remaining best networks (5/14 without accounting for phylogenetic uncertainty, and 3/12 when accounting for it). Clade 2 in this tree contained two subgroups: one comprising *O. tenthredinifera*and *O. bombyliflora* and the other comprising *O. speculum* and *O. lutea* (*fusca*).

The best-fitting number of reticulations was two based on the SNaQ analysis (Fig. 3a). The best-scoring network presented a backbone tree identical to the PhyloNet minority backbone tree. The two reticulations (Fig. 3b) were well supported by the bootstrap analysis, confirming two of the reticulations inferred by PhyloNet. As in PhyloNet, the first reticulation was an important gene flow from *O. lutea* (*fusca*) and/or *O. bombyliflora* to *O. tenthredinifera*, with an average inheritance probability of 0.50 and 91% bootstrap support. The second reticulation corresponded to a gene flow from the ancestral clade 2 lineage to *apifera* lineage, with an average inheritance probability of 0.08 and 100% bootstrap support. However, this result is likely an artefact, caused by SNaQ’s limitation to level-1 networks.

**Figure 3.**
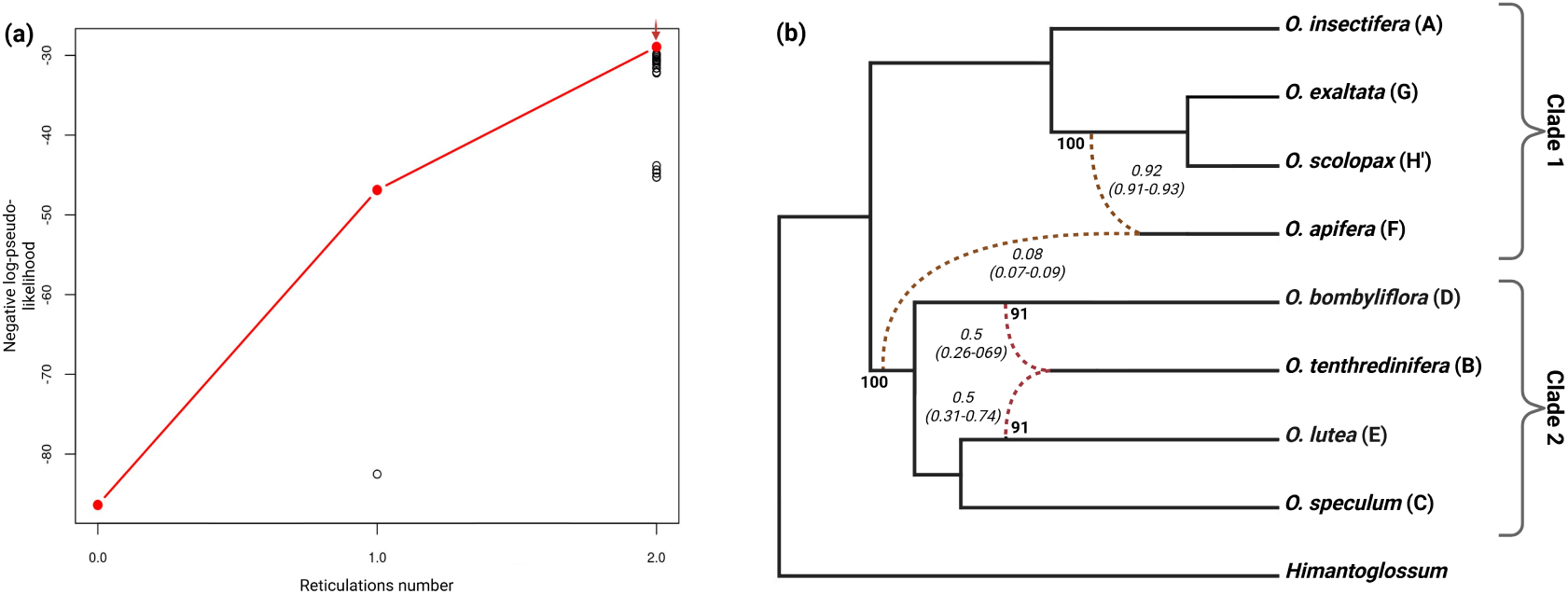
(a) Pseudo-likelihood scores of the best networks inferred with SNaQ as a function of their number of reticulations. The red dot depicts the "best" network for each number of reticulations, and the red arrow indicates the ‘best’ number of reticulations. (c) Best SNaQ network with 2 reticulations according to pseudo-likelihood scores. Dashed lines represent the reticulation events, and the associated numbers indicate the average inheritance probability rounded to the nearest hundredth of each reticulation (with min and max values associated) across the 100 bootstrap replicates. Numbers at nodes between reticulations and branches represent the bootstrap support for each reticulation. Networks were drawn with Dendroscope 3 (Huson and Scornavacca, 2012), and bootstrap support were added with Biorender (https://BioRender.com/mel8oii). Branch lengths are arbitrary.

In the best SNaQ two-reticulation network, *O. tenthredinifera*, a member of clade 2, is already involved in another reticulation event, preventing SNaQ from inferring the *O. tenthredinifera* to *O. apifera* reticulation event, which was recovered by PhyloNet. Consistently, the best SNaQ network with 1 reticulation (Supplementary Fig. S7) did recover this *O. tenthredinifera* to *O. apifera* reticulation, confirming that the clade 2 to *O. apifera* reticulation likely results from SNaQ’s level-1 network limitation. Therefore, SNaQ seems less suitable than PhyloNet for our dataset, as it cannot infer consecutive reticulate events.

The majority backbone tree topology (Supplementary Fig. S6a) and minority backbone tree topology (Supplementary Fig. S6b) were used as references to set up two distinct datasets of 1,000 clock-like genes using SortaDate, which were used to perform species divergence time estimation (presented respectively in Fig. 4 and Supplementary Fig. S8). All Clade 2 taxa had very similar divergence times, with three consecutive speciation events occurring around 3 Ma. However, the taxa in clade 1 diverged much more recently, in particular with *O. exaltata* and *O. scolopax* having diverged from each other less than 1 Ma.

**Figure 4.**
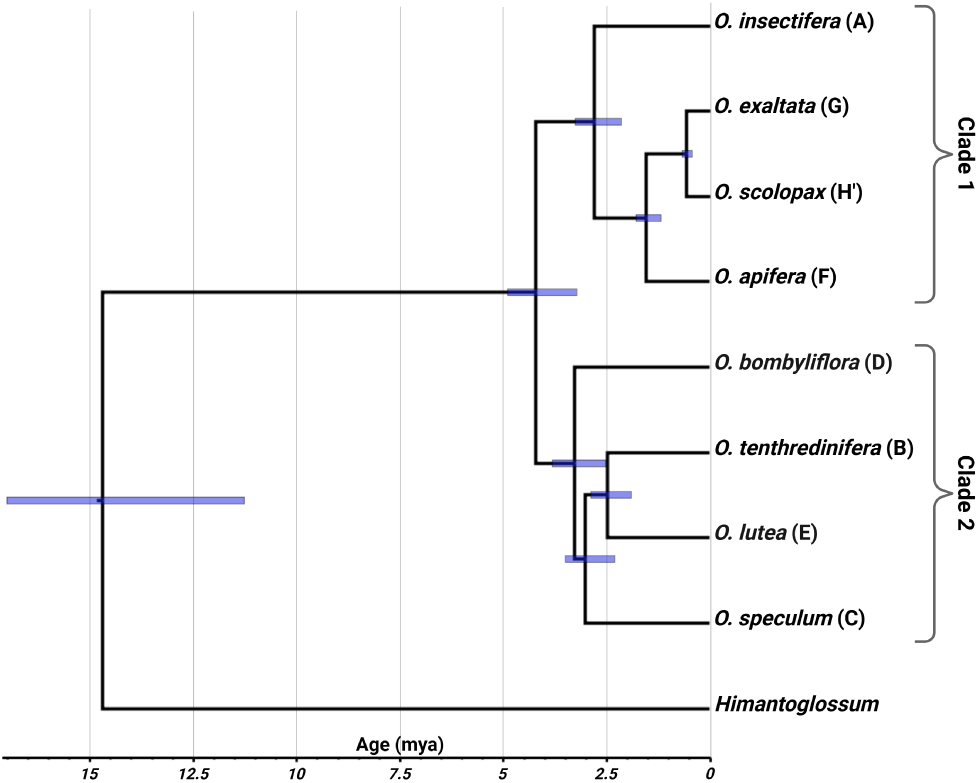
Dated majority backbone tree inferred with mcmctree, based on 1,000 clock-like genes selected using SortaDate. The scale represents the absolute age in million years. Blue bars at nodes represent 95% highest posterior density intervals. Tree was drawn with FigTree (http://tree.bio.ed.ac.uk/software/figtree/).

### Introgression Tests

Both the Δ test and Aphid require a reference topology to infer gene flow events. Their results can vary greatly depending on the topology used. In this context, we decided to use the majority backbone tree topology from network analyses as a reference. Unlike concatenation and coalescent-based phylogenetic reconstructions, this topology was inferred by explicitly taking gene flow events into account, which may provide a more realistic framework for inferring *Ophrys* evolution. However, unlike network inference, the Δ test and Aphid infer gene flow independently for each triplet. This could lead to many false positives (Hibbins and Hahn, 2022), particularly when gene flow occurs between very distinct lineages, in cases of ghost gene flow, or when multiple consecutive gene flow events occur. Given our previous results, interpreting these two tests would be particularly challenging here. In this context, they were mainly used to verify the three major reticulations of network inference.

The Δ test was conducted on 56 different triplets (Supplementary Table S2). The results were similar whether or not phylogenetic uncertainty was accounted for; only lowly significant gene flow events were different (Fig. 5). The three major gene flow events previously inferred by network inference were also supported by the Δ test, with estimates of introgressed gene proportions consistent with network inheritance probabilities:

- 16% (or 22% when accounting for phylogenetic uncertainty) of introgressed genes for the gene flow event from *O. bombyliflora* to *O. tenthredinifera*;
- 11% (or 12% ) of introgressed genes for the gene flow event from *O. apifera* to *O. scolopax* (*fuciflora* complex);
- 5% (or 4%) of introgressed genes for the gene flow event from *O. tenthredinifera* to *O. apifera*;

**Figure 5.**
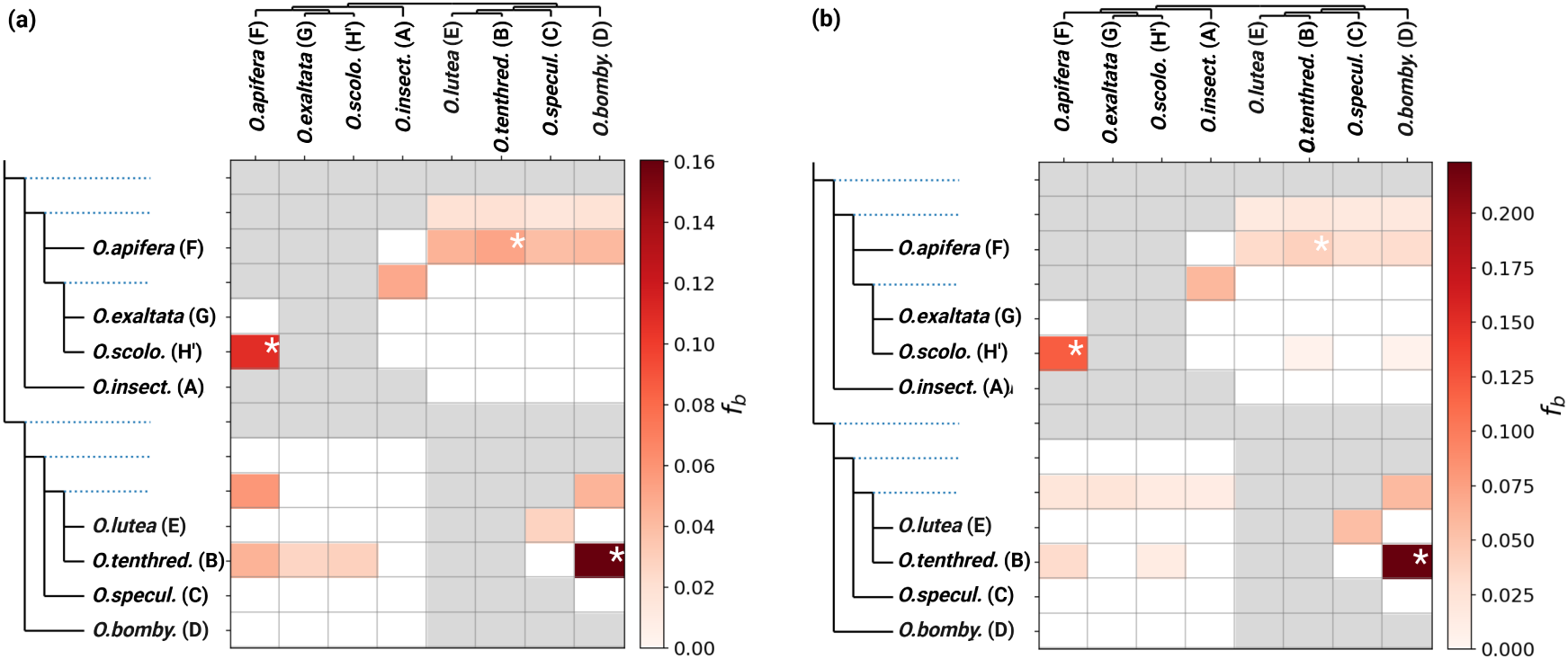
‘f branch’ (‘*f_b_*’) heatmap inferred without accounting for phylogenetic uncertainty (a) and after keeping only nodes with *≥* 70% bootstrap support (b). Color shading represents the proportion of introgressed genes calculated for each triplet presenting a significant Δ (*p*-value *<* 0.01). White asterisks represent the three gene flow inferred by PhyloNet.

The Δ test also inferred several additional gene flow events, many of which are likely artefacts due to Δ test limitations. These additional gene flow events are discussed in detail in section ‘Δ Test’ in Supplementary results.

As for the Δ test, Aphid was applied to 56 triplets (Supplementary Table S3a). 14 of these triplets presented a significant difference in the amount of topologies arising from ILS between the two discordant topologies (*I*_ILS_ significantly departed from 0.5, *p*-value *<* 0.01) and were excluded from subsequent analysis.

The likelihood ratio test did not provide a meaningful way to discriminate between gene flow events, as all the remaining triplets indicated highly significant gene flow between each pair of non-sister taxa (*p*-value *<* 0.001). We therefore relied directly on log-likelihood difference (Δ ln *L*) between models with and without this type of gene flow (Supplementary Fig. S9). Ten triplets present high Δ ln *L* values (Δ ln *L >* 3000), supporting 8 different gene flow events between non-sister taxa (presented in Fig. 6). These corresponded to the three major gene flow events previously inferred by network inference, as well as:

- gene flow between all the different clade 2 taxa (*O. bombyliflora* and *O. lutea*, *O. bombyliflora* and *O. speculum*, *O. speculum* and *O. tenthredinifera*, *O. speculum* and *O. lutea*, in addition to the major gene flow from *O. bombyliflora* to *O. tenthredinifera*). The *O. bombyliflora* to *O. tenthredinifera* gene flow remains the most supported among them (Δ ln *L >* 6000);
- a gene flow from *O. bombyliflora* to *O. apifera*. This gene flow was potentially an artefact due to the combination of the gene flows from *O. bombyliflora* to *O. tenthredinifera*, and from *O. tenthredinifera* to *O. apifera*. Indeed, in the triplet *O. tenthredinifera*, *O. bombyliflora* and *O. apifera*, the gene flow from *O. tenthredinifera* to *O. apifera* was still supported (Δ ln *L* = 3324), but this was no longer the case for the gene flow from *O. bombyliflora* to *O. apifera* (Δ ln *L* = 2203);

**Figure 6.**
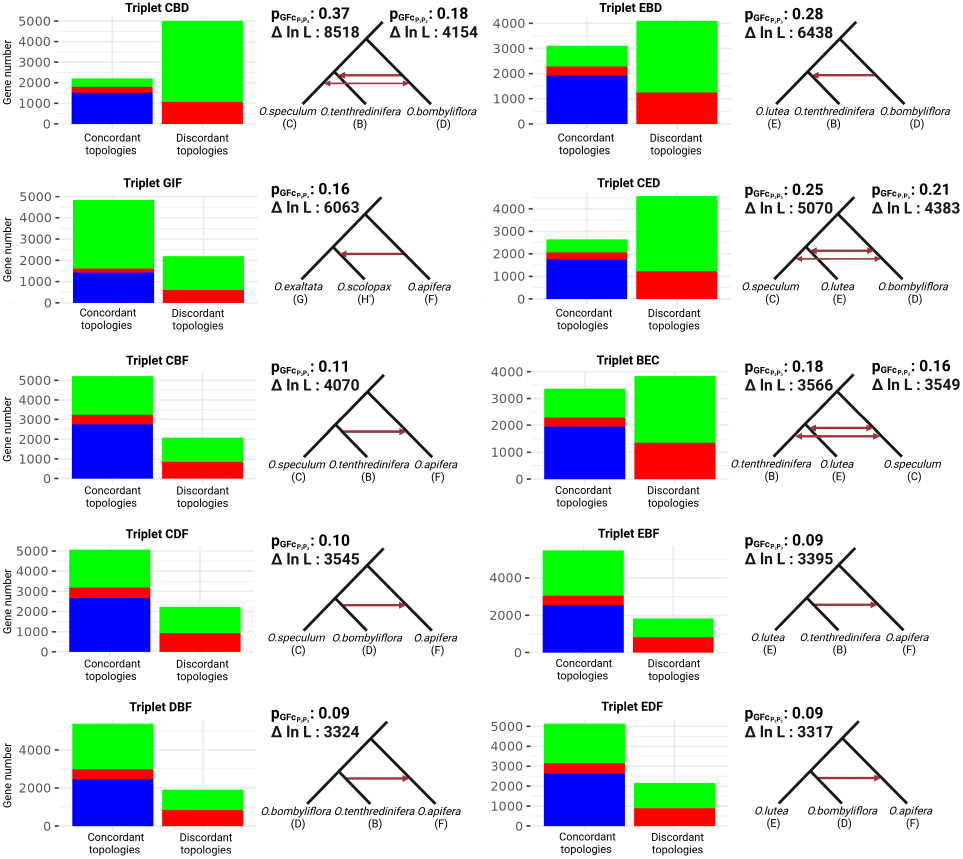
Triplets ((*P*_1_,*P*_2_),*P*_3_) with the highest Δln *L* values (Δln *L >* 3000) between models with and without gene flow between non sister taxa. Bar plots show the proportions of concordant and discordant topologies explained by incomplete lineage sorting (red), gene flow (green), or speciation (blue). Schematic trees depict the inferred gene flow events. Statistics include *p_GF c_* (proportion of introgressed genes between non sister taxa) for *P*_2_-*P*_3_ and *P*_1_-*P*_3_ gene flow, and Δln *L* associated (likelihood difference between models with and without gene flow).

Other less significant gene flow events (Δ ln *L <* 3000) are discussed in detail in the Supplementary Data. Aphid results supported the presence of an ILS background in our dataset, with an average of 1,285 genes identified as affected by ILS (against 3,863 genes identified as affected by gene flow in average) in each triplet (Supplementary Fig. S10).

Since the three major gene flow events supported by network inference were confirmed by the Δ test and by Aphid, we used Aphid to characterise them in greater detail by estimating their relative timing (Supplementary Table S4). For this purpose, Aphid estimates the parameter *p_a_*, which corresponds to the probability that gene flow occurred close to the time of speciation. When ten different timings of gene flow were considered (from close to speciation to ten times more recently), Aphid indicated that the *O. apifera* to *O. scolopax* (*fuciflora* complex) gene flow event was a mixture of an important ancient gene flow (*p_a_* = 0.56), followed by weaker and continuous gene flows extending up to the present day. The important gene flow event from *O. bombyliflora* to *O. tenthredinifera* seems to have occurred at around the same time as *O. tenthredinifera* speciation with *O. lutea* (*fusca*) (*p_a_* = 0.831). This result is consistent with a potential hybrid origin of *O. tenthredinifera*. This gene flow also occurred at around the same time as speciation between the common ancestor of *O. tenthredinifera* and *O. lutea* (*fusca*) with *O. speculum* (*p_a_* = 0.784). Finally, the gene flow from *O. tenthredinifera* to *O. apifera* seems to have occurred at around the same time as *O. tenthredinifera* speciation (*p_a_* ≈ 0.79) with its closest relatives (clade 2 taxa).

### Gene Annotation and Genome Position

Of our 7,821 orthologous genes, 7,425 were successfully mapped onto the *Ophrys* reference genome, with around half of them (3,521) mapping to one of the 18 chromosomes and the other half (3,904) mapping to unanchored scaffolds. We explored the position of putative introgressed genes for the three major gene flow events:

- from *O. apifera* to *O. scolopax* (*fuciflora* complex) (Supplementary Fig. S11, S12, S13);
- from *O. tenthredinifera* to *O. apifera* (Supplementary Fig. S14, S15, S16);
- from *O. lutea* (*fusca*) and *O. bombyliflora* to *O. tenthredinifera* (Supplementary Fig. S17, S18, S19);

Putatively introgressed genes associated with the gene flow events from *O. apifera* to *O. scolopax* and from *O. tenthredinifera* to *O. apifera* were distributed in a mosaic-like pattern across the genome (Supplementary Fig. S13, S16) . They appeared to be partially concentrated towards the ends of chromosomes, particularly for the *O. apifera* to *O. scolopax* (*fuciflora* complex) gene flow (Supplementary Fig. S12), although this may reflect a higher gene density in these regions (Supplementary Fig. S20). Notably, chromosome 1 presented a significantly higher number of introgressed genes for the *O. apifera* to *O. scolopax* gene flow event (*p*-value *<* 0.001, Supplementary Fig. S11). As the origin of the major gene flow towards *O. tenthredinifera* was uncertain (*O. bombyliflora* and/or *O. lutea* (*fusca*)), we focused on both genes that grouped *O. tenthredinifera* with *O. bombyliflora* and genes that grouped *O. tenthredinifera* with *O. lutea* (*fusca*). Across most of the genome, these two topologies were distributed in a mosaic-like pattern, although some chromosomes contained large blocks of one of the two topologies, or even blocks of other alternative topologies (Supplementary Fig. S19). These blocks were associated with an over-representation of the associated topology within the chromosomes (Supplementary Fig. S17). The most significant examples (*p*-value *<* 0.001) were chromosome 12 which presented an over-representation of topologies that grouped *O. tenthredinifera* with *O. lutea* (*fusca*), chromosome 17 with an over-representation of topologies that grouped *O. tenthredinifera* with *O. bombyliflora*, and chromosome 9 with an over-representation of other topologies.

58 *Ophrys* flower phenotype key genes out of 296 were identified in our dataset (see Supplementary Table S5). All three gene flow major events were found in these key genes:

- 2 key genes for the *O. tenthredinifera* to *O. apifera* gene flow event.
- 7 key genes for the *O. apifera* to *O. scolopax* (*fuciflora* complex) gene flow event, most of which (6/7) were confirmed as potentially introgressed by Aphid.
- 35 key genes for the *O. lutea* (*fusca*) and *O. bombyliflora* to *O. tenthredinifera* gene flow event. Aphid identified these key genes as mainly resulting from speciation or gene flow (31/35). A comparable number of key genes supported the two alternative topologies: 18 grouped *O. tenthredinifera* with *O. bombyliflora* and 17 grouped *O. tenthredinifera* with *O. lutea* (*fusca*).

## Discussion

### Phylogenetics Relationships in *Ophrys*

Based on 7,821 gene trees and 1,578,109 informative positions, the phylotranscriptomic approach that we implemented yielded an unparalleled dataset for reconstructing the *Ophrys* phylogeny. This approach unveiled for the first time the presence of three major gene flow events that stand out from an ILS background and possibly additional minor gene flow events. The different species trees that were reconstructed - whether they are coalescent-based, concatenation-based or backbone network tree-based - were, in general, consistent with the handful of ‘-omic’ studies that had been published to date for *Ophrys* (Bateman et al., 2018; Piñeiro Fernández et al., 2019; Bertrand et al., 2021a; Anthoons et al., 2025). Our results corroborate recent findings according to which the genus *Ophrys* is subdivided into two major clades (Fig. 1). The first clade is formed by the lineages *insectifera* (A), *apifera* (F), *sphegodes* (with *O. exaltata*, G) and the *fuciflora* complex (with *O. scolopax*, H’). The second one comprises the lineages *tenthredinifera* (B), *speculum* (C), *bombyliflora* (D) and *fusca* (with *O. lutea*, E). Contrary to previous Sanger-based phylogenetic hypotheses (Devey et al., 2008; Breitkopf et al., 2015), our species trees thus support the position of the *insectifera* lineage as the sister lineage to the clade formed by *apifera*, *sphegodes* and the *fuciflora* complex (as well as lineage *umbilicata* (J) not sampled in this study), rather than as sister lineage to all the other *Ophrys* species.

Additionally, our study provides new insights into the systematic relationships within clade 2. Former studies had supported a basal position for *speculum* within this clade (Breitkopf et al., 2015; Bateman et al., 2018), which is also the case in the concatenation and coalescent-based trees in our study (Fig. 1, Supplementary Fig. S2). However, this position appeared as uncertain in our split networks (Supplementary Fig. S3) as it was also the case in Bateman et al. (2018). When gene flow is taken into account, *bombyliflora* instead occupies a basal position within this clade (Supplementary Fig. S6). To test whether this incongruence might be explained by the gene flow from the *bombyliflora* and *fusca* lineages to *tenthredinifera*, we reran the analyses without *tenthredinifera*. The resulting concatenation and coalescent-based trees (Supplementary Fig. S21, S22) support a basal position of *bombyliflora* rather than *speculum* within clade 2, in agreement with phylogenetic network inference. This gene flow involving *tenthredinifera* also renders its phylogenetic position difficult to infer. *Tenthredinifera* is either grouped with *fusca*, as found in former studies (Breitkopf et al., 2015; Bateman et al., 2018), or with *bombyliflora*. Overall, it confirms that phylogenetic networks are more appropriate than phylogenetic tree reconstruction methods to infer the reticulate evolution of genera such as *Ophrys*.

Our species divergence time estimation (Fig. 4, Supplementary Fig. S8) broadly coincides with previously published studies (Inda et al., 2012; Breitkopf et al., 2015; Bateman et al., 2018). The most recent common ancestor of our *Ophrys* lineages originated around 4.5 ± 1 Ma, at the beginning of the Pliocene epoch, *i.e.* almost 10 Ma after diverging from the nearest related genus used, here *Himantoglossum*. The genus *Ophrys* would have diversified around 3 Ma, with all the studied lineages diverging around this period except *apifera*, *sphegodes* and the *fuciflora* complex that may have formed even more recently: 1.5 ± 0.5 Ma for *apifera* and less than 1 Ma for *sphegodes* / *fuciflora* complex.

The slow evolutionary rate of the plastid genome yielded a limited amount of informative positions (490 when the outgroup was discarded) despite its large size (135,806 bp). Nevertheless, it provides a well-resolved phylogenetic tree (Supplementary Fig. S23). In this tree, the *tenthredinifera* lineage groups with *bombyliflora* at the end of a long branch, suggesting that *tenthredinifera* may have acquired its plastid genome from *bombyliflora*. The basal position of *speculum* in clade 2 of the plastid tree is inconsistent with network-based inferences and potentially reflects ghost gene flow, as inferred by network inference and the Δ test on the nuclear dataset. Finally, the group formed by the lineages *apifera*, *sphegodes* and the *fuciflora* complex presents a very short internal branch compared to the nuclear phylogenies, with few differences in their plastid sequences (explaining the relatively poor bootstrap support of 84). Thus, plastid exchange potentially occurred during a gene flow event between *apifera* and the *fuciflora* complex. Overall, the plastid genome reveals a complex history potentially shaped by different gene flow events. Due to its mutation rate being different from that of the nuclear genome, this *locus* cannot be incorporated into the nuclear Aphid analysis to confirm these gene flow events.

### Orthologous Inference with Phylotranscriptomic

Accurate inference of orthologous groups from RNA-seq data remains one of the main challenges of this approach, particularly in non-model organisms lacking a high-quality reference genome. De novo transcriptome assembly is prone to incomplete assemblies, mis-assemblies, and redundancy, which can compromise ortholog identification, alignment quality, and downstream phylogenetic inference (Yang and Smith, 2013). Assembly performance is strongly influenced by both the choice of assembler and k-mer size, and no single method or parameter set consistently performs best across datasets (Zhao et al., 2011; Lu et al., 2013; Sahraeian et al., 2017; Hölzer and Marz, 2019). Small k-mer sizes tend to better recover lowly expressed transcripts but increase assembly errors, whereas large k-mer sizes miss these lowly expressed transcripts but are prone to fewer errors (Bushmanova et al., 2019). In this context, combining different assembly tools with different k-mer sizes enables a more complete and accurate assembly. To maximize transcriptome completeness while limiting assembly artefacts, we combined three high-performing assemblers (Hölzer and Marz, 2019): Trinity, Trans-Abyss and rnaSPAdes, using k-mer sizes ranging from 21 to 99. All *Ophrys* transcriptomes were sequenced and assembled in a strand-specific manner to reduce trans-chimeric assemblies (Yang and Smith, 2013). This strategy produced highly complete assemblies, with very few missing or fragmented BUSCO genes (Supplementary Table S1), but resulted in substantial redundancy, with an average of 342,175 transcripts per assembly after perfect fragment removal. Redundancy was efficiently removed using EvigeneR (Gilbert, 2013), which clusters similar sequences and selects the transcript encoding the longest protein as representative. Compared to simpler strategies (*i.e.* retaining the longest transcript), this approach minimizes erroneous sequence selection (Gilbert, 2019). As a result, the average number of transcripts per assembly was reduced from 342,175 to 28,156, while maintaining a comprehensive dataset (see Supplementary Table S1).

Despite these improvements, some homologous gene alignments still exhibited assembly-related artefacts, including fragmented sequences, occasional chimeras, and poorly aligned or non-homologous regions, likely arising from residual assembly errors or isoform-specific exons. To further improve alignment quality, we applied two additional filters: (i) removal of short sequences (*<* 300 bp), assembled with only a few reads, which predominantly corresponded to fragmented or low support sequences; and (ii) iterative removal of low-similarity alignment regions using Hmmcleaner. Filtering was repeated until no further sequences or regions were removed. Finally, we applied the Branch Length Comparison method (Simion et al., 2020), which excluded 332 alignments, primarily containing fragmented sequences, distinct paralogous genes, or insufficient phylogenetic information. This filtering step did not appear to have reduced gene flow signals in our dataset, as indicated by Aphid analyses on excluded alignments (see ‘Orthologous gene alignments filtering details’ section in Supplementary methods and Supplementary Table S6). Overall, our multi-step assembly, redundancy reduction, and filtering strategy yielded highly complete and reliable transcriptomes and a robust ortholog dataset, suitable for investigating gene flow during *Ophrys* diversification.

### Gene Flow Inference based on Gene Trees

The gene tree-based approaches employed here rely on the assumption that each gene has a single evolutionary history (*i.e.* no intra-genic recombination). Because this assumption is central to our analyses, we explicitly tested it (see ‘Evaluation of Gene-Wise Recombination’ section in Supplementary methods and Supplementary Fig. S24) and found it is generally upheld across our dataset. Branch-length-based tests such as Aphid additionally assume that branch lengths are shaped only by speciation, gene flow, or ILS. Since selection or mutation rate variation can also impact branch lengths, Aphid filters out genes with excessively high or low mutation rates estimates, leading to the exclusion of between 500 and 787 gene trees depending on the triplet.

Our results also underscore the limitations of triplet-based approaches in lineages with complex evolutionary histories. Such methods are particularly susceptible to false positives arising from ‘ghost’ gene flow (Hibbins and Hahn, 2022), since they analyse only three taxa at a time. In this study, gene flow from *tenthredinifera* to the *apifera* lineage likely produced false positive signals in the Δ test. For example, triplets including *O. insectifera*, *O. apifera*, and *O. scolopax* (or *O. exaltata*) presented significant gene flow between *O. insectifera* and *O. scolopax* (or *O. exaltata*), which is most plausibly explained by gene flow between *O. apifera* and *O. tenthredinifera*, rather than by direct gene flow between the former taxa. Triplet-based approaches also strongly depend on a reference topology to determine the origin and direction of gene flow. Both Aphid and the Δ test test supported gene flow from *O. bombyliflora* to *O. tenthredinifera* when the concatenation-based topology was used as a reference, whereas gene flow was instead inferred from *O. lutea* to *O. tenthredinifera* under the coalescent-based topology (Supplementary Fig. S25).

Phylogenetic network inference avoids these issues by using the whole dataset and estimating both species tree topology and gene flow events simultaneously. This approach is also generally more sensitive to complex patterns of gene flow than triplet-based methods (Hibbins and Hahn, 2022). Network inference, however, remains computationally demanding for large genomic datasets. As maximum likelihood and Bayesian methods were infeasible here, we used pseudo-likelihood methods and a reduced taxon set. We therefore limited our analysis to a single representative species per lineage, at the cost of restricting the interpretation of our results in an evolutionary context. Network searches are heuristic and the optimal solution cannot be guaranteed. This problem worsens with increasing reticulation levels as the search space grows exponentially and searches risk becoming trapped in local optima. This issue is more pronounced in PhyloNet than in SNaQ, since SNaQ restricts analyses to level-1 networks, reducing the search space particularly for large reticulation numbers. To minimise suboptimal solutions, multiple independent runs are recommended, especially in groups with expected multiple overlapping gene flow events (Kong et al., 2025). We therefore performed extensive replication, particularly with PhyloNet (three independent analyses of 100 replicates each), and optimised branch lengths and inheritance probabilities for all candidate networks. Overlapping gene flow cannot be inferred by approaches restricted to level 1 networks such as SNaQ, unlike higher-level networks (*i.e.* ≥ 2) methods such as PhyloNet. However, these higher-level networks face identifiability problems: distinct evolutionary scenarios may produce identical sets of expected gene tree topologies and thus identical likelihoods, leading to distinct networks with the same ‘canonical forms’ (Pardi and Scornavacca, 2015). As a consequence, the relative timing of overlapping gene flow events may not be fully resolved from network inference alone, since networks with the same reticulation but in a different order can present the same canonical forms.

### Major Reticulation Events in addition to overall Incomplete Lineage Sorting (ILS) during the *Ophrys* Diversification

Overall, our results revealed extensive gene tree discordances in *Ophrys*, driven by multiple gene flow events and, to a lesser extent, by ILS (Supplementary Fig. S10). We identified three major gene flow events during the diversification of the genus (Fig. 7). The first involves gene flow from *apifera* (F) to the *fuciflora* complex (with *O. scolopax*, H’), two lineages that are closely related but not sister groups. Our analyses suggest a combination of several introgression events, including an early event shortly after the divergence between the *sphegodes* (with *O. exaltata*, G) lineage and the *fuciflora* complex, followed by continuous gene flow persisting up to the present (Supplementary Table S4). The recent gene flow signal may be specific to the species or the populations sampled. Naturally occurring hybrids between *O. apifera* and *O. scolopax* can be observed (*Ophrys* x *minuticauda*), despite differences in flowering times. Some described taxa, such as *O. corbariensis* (Samuel and Lewin, 2002), have even been proposed to have a hybrid origin between these two lineages, based on floral and phenological similarities, although genetic confirmation is still lacking.

**Figure 7.**
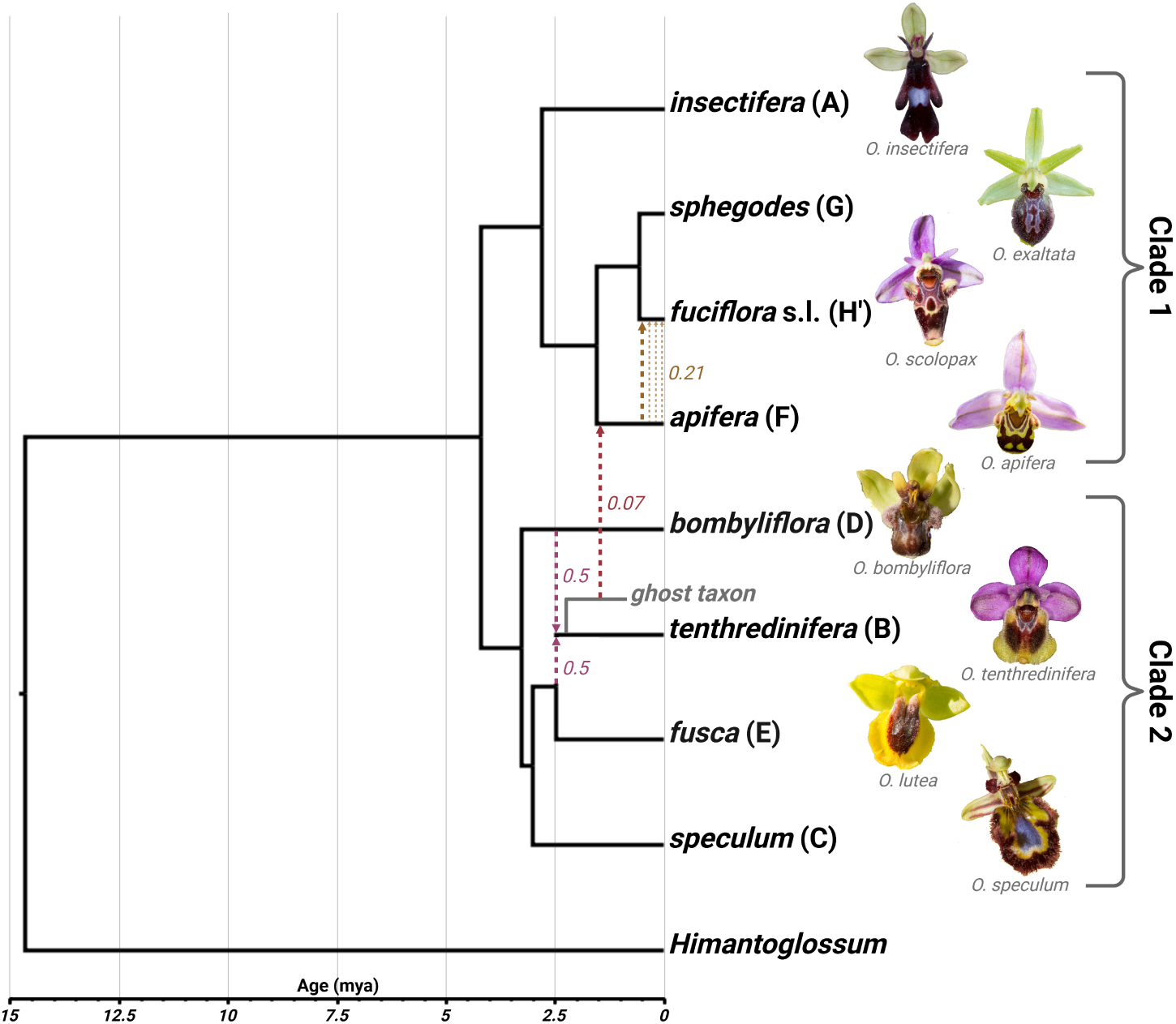
Simplified *Ophrys* evolution scenario, based on the dated network majority backbone tree (Fig. 4, 95% highest posterior density intervals of each node are not represented). Dashed lines represent the three most supported reticulation events, with associated inheritance probability (average inheritance probability from Fig. 2c). Reticulations are replaced on the backbone tree relatively to speciation events, based on Aphid results (Supplementary Table S4). *Fuciflora* sensu lato (*fuciflora* s.l.) corresponds to the *fuciflora* complex. Tree was drawn with FigTree and modified with Biorender (https://BioRender.com/590w2hx).

The second gene flow event involves the *bombyliflora* (D), *tenthredinifera* (B) and *fusca* (with *O. lutea*, E) lineages. This event is substantial in magnitude (inheritance probabilities ∼ 0.5) and appears to reflect gene flow either from *bombyliflora* to *tenthredinifera* or from *fusca* to *tenthredinifera*. Aphid results show that genes grouping *tenthredinifera* with either of these lineages present similar branch lengths, suggesting that the *tenthredinifera* lineage may have originated through homoploid hybrid speciation between ancestors of the *bombyliflora* and *fusca* lineages. The inferred timing of this admixture, approximately 300,000–500,000 years after the divergence between *speculum* and *fusca* as suggested by Aphid (Supplementary Table S4) and species divergence time estimations (Fig. 4, Supplementary Fig. S8), is consistent with such a scenario but does not allow a definitive conclusion. Distinguishing hybrid speciation from introgression remains challenging (Hibbins and Hahn, 2022). Evidence of strong admixture represents only one of the three required criteria (Long and Rieseberg, 2025), and can also be produced by extensive continuous gene flow (Kong et al., 2025). The other two, reproductive isolation from parental species and hybrid-derived reproductive barriers, require further investigation. Assessing these criteria in *Ophrys* is difficult because reproductive isolation is mainly pre-zygotic (Scopece et al., 2007). Although the *tenthredinifera*, *bombyliflora* and *fusca* lineages remain interfertile, natural hybrids are relatively rare in the wild due to pronounced differences in floral traits and pollinator specificity.

The third major event involves gene flow from the *tenthredinifera* lineage to *apifera*. Several genes support both this event and the one between *apifera* and the *fuciflora* complex. In such cases, *apifera* and the *fuciflora* complex form a sister clade to *tenthredinifera*, indicating that the *tenthredinifera* to *apifera* gene flow event predates the *apifera* to the *fuciflora* complex one. Interestingly, 12 of the 14 triplets rejected by Aphid *I*_ILS_ filter (Supplementary Table S3a) presented an excess of discordant topologies arising from ILS grouping *apifera* with *tenthredinifera* or its close relatives, a pattern expected in the presence of a gene flow from a ‘ghost’ taxon closely related to *tenthredinifera* rather than from *tenthredinifera* itself (see section ‘Ghost gene flow detection with Aphid’ in Supplementary results for details). In these rejected triplets, the timing of gene flow inferred by Aphid likely corresponds to the divergence between *tenthredinifera* and this unsampled taxon, which is therefore close to *tenthredinifera* divergence with its closest relatives. Because current species within the *tenthredinifera* lineage diverged only recently (Devey et al., 2008; Bateman et al., 2018), and all extant closely related lineages of *tenthredinifera* were included in our dataset, the ghost donor is most likely an extinct *Ophrys* lineage. This ghost gene flow cannot have been detected using topology-based methods alone, such as pseudo-likelihood network inference or the Δ test, underscoring the value of combining branch-length-based approaches like Aphid with topology-based tools.

The mosaic distribution of potentially introgressed genes between *apifera* and the *fuciflora* complex (Supplementary Fig. S12), or between *tenthredinifera* and *apifera* (Supplementary Fig. S15) is consistent with ancient introgression, as recombination progressively breaks long blocks into smaller fragments over time (Martin and Jiggins, 2017). In contrast, the putative hybrid origin of *tenthredinifera* shows a more complex pattern: although a genome-wide mosaic is present (Supplementary Fig. S18), some chromosomes (*e.g.* 9, 12, and 17) present large homogeneous blocks of genes supporting the same topology (*tenthredinifera* with *bombyliflora*; *tenthredinifera* with *fusca* or other alternatives). The persistence of such blocks may reflect secondary gene flow, selection maintaining intact haplotypes, or reduced recombination rates associated with structural genomic variants such as inversions or supergenes (Edelman et al., 2019). The identification of genomic islands of divergence at the end of chromosome 2 in the *sphegodes* lineage (in *Ophrys aveyronensis* (Gibert et al., 2025) and other species (Russo et al., 2024)) supports the existence of such structural variation in *Ophrys*. Notably, this genomic region of around 7-22 Mb contains a cluster of potentially introgressed genes from *apifera* to the *fuciflora* complex, including the key gene VPS45 likely to be involved in *Ophrys* odour variation.

### Ecological Consideration of Gene Flow Events

The three major gene flow events inferred from our sequence data are also relevant from phenotypic and ecological perspectives. Both the *apifera* (F) lineage and the vast majority of species in the *fuciflora* complex (H’) are pollinated by *Eucera* bees and share similar floral traits in terms of size, shape and colouration. The *tenthredinifera* (B) lineage, also predominantly *Eucera*-pollinated, exhibits an outstanding flower morphology within clade 2, similar to that of several clade 1 lineages: *apifera*, *fuciflora* complex, *sphegodes* (G) and *umbilicata* (J), despite their phylogenetic distance. This similarity likely reflects evolutionary convergence due to *Eucera* pollination adaptation. Although the *tenthredinifera*, *bombyliflora* (D) and *fusca* (E) lineages differ in floral phenotype, especially the *fusca* lineage which exhibits unique flower morphology within the genus, the putative hybrid origin of the *tenthredinifera* lineage is ecologically plausible. The *bombyliflora* lineage is pollinated by *Eucera* bees (Joffard et al., 2019) and shares unique morphological traits with *tenthredinifera*. These two lineages were historically grouped in morphological classifications (Devillers and Devillers-Terschuren, 1994; Delforge, 2006). While no extant species in the *fusca* lineage are pollinated by *Eucera* bees, the lineage was likely ancestrally *Eucera*-pollinated with a relatively recent shift, mainly to *Andrena* and other bee genera (*Antophora*, *Colletes*, *Megachile* and *Osmia*), occurring around 1 Ma (Breitkopf et al., 2015). The extinct ghost lineage, identified as the source of the gene flow to *apifera*, was closely related to *tenthredinifera* and therefore likely also *Eucera*-pollinated.

Overall, our major gene flow events correspond to ancient exchanges among lineages that are mainly or ancestrally pollinated by *Eucera* bees. Their detection in several key floral-trait genes (Supplementary Table S5) suggests that these events may have been adaptive, potentially involving the transfer of genes associated with *Eucera* pollination. This pattern supports an initial diversification of *Ophrys* linked to *Eucera* pollination 2-3 million years ago, during which interspersed gene flow may have facilitated lineage radiation. Later, independent shifts toward *Andrena* and other pollinators (mainly bees), particularly in the *fusca* (E) and *sphegodes* (G) groups, led to their more recent and extensive diversification (Breitkopf et al., 2015), but did not produce significant inter-lineage gene flow between them. This may reflect the higher specificity of *Andrena* pollination compared to the more colour-based mate-location behaviour of *Eucera* bees (Spaethe et al., 2010). The exceptional pollinator diversity in the Mediterranean especially among *Eucera* and *Andrena* bees whose diversification predates *Ophrys* (Cardinal and Danforth, 2013), likely enabled the genus’s remarkable radiation. The greater species richness and specificity of *Andrena* bees may explain why *Andrena*-pollinated *Ophrys* lineages are more diverse overall. Small changes in floral odour can produce shifts in pollinator attraction and generate reproductive isolation (Sedeek et al., 2014; Xu et al., 2011). Although artificial crosses frequently yield viable hybrids, suggesting generally weak, though not completely absent, post-zygotic barriers in *Ophrys* (Scopece et al., 2007; Cortis et al., 2009), strong pre-zygotic isolation due to pollinator specificity maintains species boundaries in most situations (Baguette et al., 2020). Our detection of only a few ancient gene flow events, rather than pervasive introgression, aligns with this view. Together, these findings highlight the predominance of pre-zygotic reproductive isolation in *Ophrys* and illustrate the inherent difficulty of defining species limits in a group where gene flow is episodic yet possible over evolutionary timescales.

### Possibility of Ghost Introgression from a Basal Extinct Lineage

Introgression tests based on gene-tree topologies indicate a possible ghost introgression event from an extinct taxon or unsampled outgroup at the base of the *Ophrys* genus, primarily towards the *speculum* (C) and *insectifera* (A) lineages (Supplementary Fig. S4, S5, S25). However, this pattern may reflect long-branch attraction (Philippe et al., 2005). The long branch separating the outgroup *Himantoglossum* (see Fig. 1) may artificially ‘attract’ long-branched *Ophrys* lineages, such as *insectifera* and *speculum*, toward the base of the trees. Topology-based methods (*e.g.* the Δ test or network inference) may then misinterpret this as ghost introgression. Although *Himantoglossum* genus is quite divergent, it remains one of the closest living relatives of *Ophrys* (Inda et al., 2012). The only closer, *Steveniella*, occurs outside the sampling area and is also separated from *Ophrys* by a long branch (Bateman et al., 2018). Given the substantial phylogenetic distance between *Ophrys* and its nearest extant relatives, the existence of extinct sister genera or basal extinct *Ophrys* lineages is plausible.

Interestingly, *speculum* and *insectifera* lineages are among the most phenotypically divergent lineages within *Ophrys*. They include species with typical wasp pollination and a distinctive floral morphology (*e.g.* elongated labellum). This wasp pollination has been interpreted as ancestral for *Ophrys*, due to the inferred basal placement of these lineages. Yet, the basal position of the *insectifera* lineage is no longer supported by -omic data, and only one of its three described species (*O. insectifera* s.s) is wasp-pollinated (*O. aymoninii* and *O. subinsectifera* being pollinated by an *Andrena* bee and a sawfly respectively). Moreover, our analyses place *bombyliflora* rather than speculum in clade 2 basal position once gene flow is accounted for. Together, these results challenge the hypothesis of ancestral wasp pollination in *Ophrys*.

## Conclusions

Although reticulate evolution was assumed in the case of the genus *Ophrys*, our study provides the first evidence of three major and structuring gene flow events at an early stage of the genus diversification (between 3 and 1 Ma). All *Ophrys* individuals sequenced in this study display levels of heterozygosity (∼ 10%; Supplementary Table S7) consistent with values previously found within this genus (Russo et al., 2024), suggesting they are not of recent hybrid origin. The global mosaic distribution of potentially introgressed genes along the chromosomes, rather than in large intact blocks, supports ancient gene flow with subsequent recombination rather than recent hybridization.

These elements, combined with Aphid gene flow relative datation, support that these gene flow may be ancient and fixed across extant species. Of course, such pattern will have to be supported by incorporating additional taxa for each lineage, ideally from different geographical origins.

Nevertheless, these encouraging results call for a thorough investigation of the functional impact of gene flow events on *Ophrys* floral phenotype and their potential adaptive role. Although we identified in our introgressed genes some candidates likely involved in odour production consistently found as outliers in different *Ophrys* studies (*e.g.* Sedeek et al., 2014; Russo et al., 2024; Gibert et al., 2025), upcoming studies should integrate additional data, such as the expression levels of phenotypically relevant genes (*e.g.* flower morphology, colour and odour). Analysing phenotypic traits in a comparative context will also allow to better assess whether they are consistent with the phylogenetic signal, or better explained by other phenomena such as convergence and/or adaptive gene flow. In this way, going from phylotranscriptomics towards whole genome resequencing data will be crucial to better understand the genomic architecture of adaptive introgression, the emergence of novel phenotypic trait combinations, and their actual role in *Ophrys* radiation.

At a more micro-evolutionary scale, recent research on the *insectifera* lineage suggests that additional gene flow may have influenced the diversification of certain clades (Salvado et al., 2025). Overall, *Ophrys* appears to be a promising model system to investigate the contribution of both early, highly structuring gene flow events and more recent, perhaps pervasive, gene flow in a fascinating example of plant adaptive radiation. Investigating recent gene flow events among closely related *Ophrys* species within a single lineage, rather than ancient exchanges between lineages, as documented here, would help to determine whether gene flow in *Ophrys* is relatively frequent over short timescales but rarely persistent, or whether it is rare but tends to leave long-lasting signatures.

## Acknowledgements and Funding

We thank Bertrand Schatz and Michel Baguette for seminal ideas and fruitful exchanges at early stages of this work. We also thank Nicolas Galtier for his constructive feedback and insightful discussions. This work was primarily supported by a ‘Bonus Qualité Recherche’ grant from the Université Perpignan Via Domitia and an Agence Nationale de la Recherche Jeune Chercheur Jeune Chercheuse (ANR JCJC) grant to J.A.M.B., grant number ANR-21-CE02-0022-01 and is supported by the ‘Laboratoires d’Excellences (LABEX)’ TULIP [ANR-10-LABX-41]. Lucas Vandenabeele PhD is funded by the ‘École Universitaire de Recherche (EUR)’ TULIP-GS (ANR-18-EURE-0019).

## Supplementary Materials and Data Availability

Sequencing data have been submitted to the European Nucleotide Archive (ENA; https:/www.ebi.ac.uk/ena/browser/home) under study with primary accesion number: PRJEB105712 (and secondary accession number: ERP186863) and samples accession number: ERS28372664 to number: ERS28372676.

