## Supplementary Information for "Phylotranscriptomics Allows Distinguishing Major Gene Flow Events from Incomplete Lineage Sorting in Rapidly Diversifying Mimetic Orchids (Genus *Ophrys*)"

### Supplementary Methods

#### Orthologous Gene Alignments Filtering Details

***Paralogous gene identification.*** Inparalogs were defined as sequences from the same species that formed monophyletic groups within gene trees. Distinct ancient paralogs, on the other hand, were defined as two clusters within a gene tree comprising at least five and nine taxa, respectively, that were separated by a branch length at least ten times greater than the gene tree's average branch length.

***Branch length comparison method (Simion et al., 2020).*** To detect excessively divergent sequences in the orthologous gene alignments, a species tree was constructed using RAxML v8.2.12 (Stamatakis, 2014) under a GTR+GAMMA model from the first SCaFoS concatenated sequence. This tree was then used as a reference. For each orthologous gene, branch lengths were estimated on this fixed species tree topology using the same GTR+GAMMA model and RAxML. Orthologous genes with external branches more than three times longer than the corresponding branches in the species tree and/or with a correlation coefficient ($R^2$) between their branch lengths and those of the species tree below 0.8 were discarded.

This filter should not be too stringent in order to avoid completely eliminating gene flow signals. To evaluate this assumption, gene trees were inferred for the 206 genes with an $R^2$ value below 0.8 using IQ-TREE 2 with the 'model selection' option. These 206 gene trees were then added to our final dataset of 7,821 gene trees, and a new aphid analysis was conducted using ten different triplets (DBA, IGA, DBF, CDF, GIF, CEI, GIB, AGC, CBD and EBD). The proportion of genes removed during the aphid pre-processing steps, as well as the proportion of the remaining genes inferred to originate from gene flow, were subsequently compared between the 206 filtered genes and the 7,821 genes of the final dataset.

#### Evaluation of Gene-Wise Recombination

To verify the assumption of no recombination within each gene underlying most of our introgression tests, we assessed whether each gene exhibited a single evolutionary history. For this purpose, we split each alignment into two halves containing an equal number of informative sites. We then reconstructed a phylogenetic tree for each half using RAxML v8.2.12 under a GTR+GAMMA model, constrained to the topology of the concatenation-based tree. We then calculated the correlation coefficient ($R^2$) between the branch lengths of the two trees. A low $R^2$ indicates substantial differences in phylogenetic signal between the two alignment halves, which is consistent with potential recombination. A threshold of $R^2 \geq 0.8$ has been used to identify genes with no recombination. Indeed, we expect genes without recombination to present $R^2$ values below 1 in this test. Comparing branch lengths from gene trees inferred using only half the genes will introduce several stochastic errors. The vast majority of gene trees (88%) displayed high $R^2$ values ($\geq 0.8$), confirming that the assumption of no recombination is generally valid in our dataset (Supplementary Fig. S24).

#### Heterozygosity Rate Estimation

To ensure that the sequenced individuals did not exhibit very recent, individual-specific gene flow, we roughly estimated heterozygosity rate of each *Ophrys* and *Himantoglossum*. Individuals with recent gene flow would be expected to exhibit unusually high heterozygosity. For each individual, the filtered reads obtained after Trimmomatic were mapped to the final transcriptome assembly using Bowtie 2 (Langmead and Salzberg, 2012), using Trinity script run_bowtie2.pl. To obtain base-level information for heterozygosity estimation, we generated mpileup files from the coordinate-sorted BAM alignments using Samtools v1.13 (Danecek et al., 2021). Each position showing at least two different nucleotides was considered heterozygous. The heterozygosity rate was then calculated as the proportion of heterozygous positions over the total number of covered positions.

### Supplementary Results

#### Δ Test

In addition to the three major gene flow events supported by network inference, the **Δ** test supported several other gene flow events (Fig. 5, Supplementary Fig. S25). However, many of these are potential artefacts:

- Gene flow events between *O. apifera* and other species close to *O. tenthredinifera* (*O. bombyliflora*, *O. lutea* (*fusca*) and *O. speculum*); between *O. tenthredinifera* and other species close to *O. apifera* (*O. scolopax* (*fuciflora* complex) and *O. exaltata* (*sphegodes*)); or between *O. insectifera* and the *O. scolopax*/*O. exaltata* subgroup, could be artefacts due to the gene flow between the very distinct taxa *O. tenthredinifera* and *O. apifera*.

- Gene flow events between the taxa *O. apifera*, *O. scolopax* (*fuciflora* complex), *O. exaltata* (*sphegodes*) and clade 2 may reflect a deep introgression between these two *Ophrys* groups, or a ghost introgression, from the base of the *Ophrys* tree into *O. insectifera* as suggested by some network reconstructions. Similarly, when accounting for phylogenetic uncertainty, a gene flow with low support ($0.001 < p < 0.01$) is assessed between the taxa *O. tenthredinifera*, *O. lutea* (*fusca*), *O. bombyliflora* and clade 1, which may reflect a ghost introgression from the base of the *Ophrys* tree into *O. speculum*. This last gene flow is more noticeable when concatenation or consensus-based topologies are used as a reference (Supplementary Fig. S25);

- A gene flow event between *O. bombyliflora* and *O. lutea* (*fusca*) (in addition to the important *O. bombyliflora*/*O. tenthredinifera* gene flow) is well supported with 4% (or 6%) of introgressed genes;

- Gene flow events between *O. speculum* and *O. lutea* or between *O. scolopax* and *O. tenthredinifera*/*O. bombyliflora* are also inferred, but with low support ($0.001 < p < 0.01$). The latter likely results from a small number of genes supporting a topology in which *O. apifera* and *O. scolopax* cluster together as a sister group to *O. tenthredinifera*, and, to a lesser extent, to other closely related taxa such as *O. bombyliflora*).

#### Aphid

To determine whether gene trees originate from incomplete lineage sorting (ILS), gene flow or speciation, Aphid calculates mutation rates based on the distance between the outgroup and ingroup triplets. In our dataset, the strong divergence between the outgroup *Himantoglossum* and *Ophrys* triplets could bias this estimation. To minimise such bias, Aphid was rerun for the 8 triplets testing for gene flow within clades 1 and 2, using a taxon from the opposite clade as the outgroup (*O. speculum* for clade 1, *O. insectifera* for clade 2; results are presented in Supplementary Table S3b). Gene flow from *O. apifera* to *O. scolopax* and from *O. bombyliflora* to *O. tenthredinifera* are still associated with a high $\Delta \ln L$ value (respectively $\Delta \ln L = 3590$ and $\Delta \ln L = 3491$), but this is no longer the case for the other gene flow between clade 2 taxa (*O. bombyliflora* and *O. lutea*, *O. bombyliflora* and *O. speculum*, *O. speculum* and *O. tenthredinifera*, *O. speculum* and *O. lutea*). Triplets FIA (*O. apifera*, *O. scolopax* and *O. insectifera*) and FGA (*O. apifera*, *O. exaltata*, *O.insectifera*) are also no longer rejected by the $I_{\text{ILS}}$ filter.

Most of the 48 triplets focussing on potential gene flow between the two main clades of *Ophrys* (clades 1 and 2) present similar $\Delta \ln L$ values ($\Delta \ln L \approx 2000$), with comparable contributions from ILS and gene flow (about 1000 genes each). We therefore tested the hypothesis that Aphid may overestimate the number of gene trees attributed to gene flow due to the very long branch separating the outgroup from *Ophrys*. Indeed, given similar internal branch lengths within the triplet, a longer outgroup branch can artificially inflate mutation rate estimates, potentially causing Aphid to misclassify discordant topologies as resulting from gene flow.

To evaluate this, we selected 4 representative triplets from the 48. For each triplet, we extracted the branch length separating *Ophrys* from *Himantoglossum* across discordant topologies assigned to ILS or gene flow. The distributions of branch lengths were then compared between the two groups (ILS and gene flow) using a non-parametric Wilcoxon rank-sum test. In all tested triplets (Supplementary Fig. S26), discordant topologies attributed to gene flow exhibited significantly longer outgroup branch lengths than those attributed to ILS (*p*-value $< 0.001$), supporting the hypothesis of a potential overestimation of gene flow in these cases.

In theory, Aphid is also able to identify potential gene flow between sister species within the internal triplet. As for non-sister taxa gene flow events, the likelihood ratio test did not provide a meaningful way to discriminate between gene flow events, as all the triplets indicated highly significant gene flow between each pair of sister taxa (*p*-value $< 0.001$). We therefore relied directly on log-likelihood difference ($\Delta \ln L$) between models with and without gene flow (Supplementary Fig. S27). One gene flow events between sister taxa was identified, associated with the highest

$\Delta \ln L$ values ($\Delta \ln L > 8000$). It is an important gene flow between the two sister species, *O. scolopax* (*fuciflora* complex) and *O. exaltata* (*sphegodes*). Most of the genes grouping these two lineages were identified as originating from gene flow by Aphid, a pattern that could be explained by a continuous and extensive gene flow between them. It could also be, at least in part, an artefact due to gene flow between *O. scolopax* and *O. apifera*, which would have blurred the speciation signal between *O. scolopax* and *O. exaltata* in triplets without *O. apifera*. However, Aphid's ability to identify this kind of gene flow has not yet been validated (Galtier, 2024), so these results should be treated with caution.

#### Ghost Gene Flow Detection with Aphid

Aphid analyse the topology and branch lengths of rooted quartets, a subset of four taxa consisting of an inner triplet and an outgroup. Each triplet consists of three taxa ($P_1$, $P_2$, and $P_3$), with three possible topologies: a 'matching topology', which corresponds to the species tree topology (($P_1$,$P_2$),$P_3$); and two 'discordant topologies' (($P_1$,$P_3$),$P_2$) and (($P_2$,$P_3$),$P_1$). These topologies have three possible origins: speciation (or no-event) , which is only present in the (($P_1$, $P_2$),$P_3$) topology; incomplete lineage sorting (ILS), which is present in equal amounts in all three topologies; and gene flow (GF). Depending on the origin of the triplet, the branch lengths between the two closest taxa are expected to differ: compared to triplets originating from speciation, branch lengths are expected to be shorter if GF has occurred, and longer if ILS has occurred. Aphid also calculates different statistics:

- the ILS imbalance ($I_{\text{ILS}}$), which is the difference in the amount of ILS between the two discordant topologies. When an equal amount of ILS is present in the two discordant topologies, $I_{\text{ILS}} = 0.5$.
- the probability of ancient gene flow $p_a$, which corresponds to the probability that gene flow occurred close to the time of speciation (and not twice as recent).
- the log-likelihood difference between models with and without gene flow ($\Delta \ln L$). Here, we will focus on gene flow between taxa $P_2$ and $P_3$ (and so on $\Delta \ln L_{P_2 P_3}$).

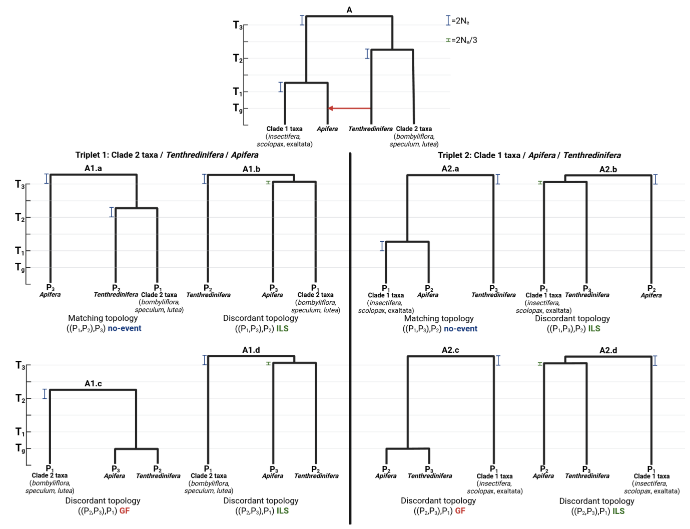


**Figure A1.** Expected gene trees (based on figure from (Galtier, 2024)) in case of gene flow from *O. tenthredinifera* to *O. apifera* (as represented in tree A), for 2 different triplets. $T_3, T_2, T_1$ correspond to divergence times and $T_g$ to gene-flow time. $N_e$ corresponds to effective population size. Red arrow reprensents gene flow event. Created in BioRender (https://BioRender.com/wuuzmj8)

Here, we focused on two different types of triplet:

- Triplet 1, with $P_1$ = clade 2 taxa (*O. bombyliflora*, *O. speculum* or *O. lutea*), $P_2$ = *O. tenthredinifera* and $P_3$ = *O. apifera*. Corresponds to triplets DBF (*O. bombyliflora*/*O. tenthredinifera*/*O. apifera*) , EBF (*O. lutea*/*O. tenthredinifera*/*O. apifera*) and CBF (*O. speculum*/*O. tenthredinifera*/*O. apifera*).
- Triplet 2, with $P_1$ = clade 1 taxa (*O. insectifera*, *O. scolopax* or *O. exaltata*), $P_2$ = *O. apifera* and $P_3$ = *O. tenthredinifera*. Corresponds to triplets AFB (*O. insectifera*/*O. apifera*/*O. tenthredinifera*), IFB (*O. scolopax*/*O. apifera*/*O. tenthredinifera*) and GFB (*O. exaltata*/*O. apifera*/*O. tenthredinifera*).

In the case of gene flow from *O. tenthredinifera* to *O. apifera* (i.e. from $P_2$ to $P_3$ in triplet 1 and from $P_3$ to $P_2$ in triplet 2), as inferred by **Δ** test (Fig. 5) and phylogenetic network inference (Fig. 2), we considered three divergence times ($T_3$, $T_2$, $T_1$) and one gene-flow time ($T_g$):

- $T_3$ the speciation time between the two *Ophrys* clades, clade 1 (*O. insectifera*, *O. apifera*, *O. scolopax* and *O. exaltata*) and clade 2 (*O. bombyliflora*, *O. tenthredinifera*, *O. speculum* and *O. lutea*).

- $T_2$ the speciation time between *O. tenthredinifera* and other taxa from clade 2.

- $T_1$ the speciation time between *O. apifera* and other taxa from clade 1. As inferred in the dated network backbone trees (Fig. 4, Supplementary Fig. S8) and concatenation-based tree (Fig. 1), $T_1$ is expected to be much lower than $T_2$ when $T_1$ represents the speciation between *O. apifera* and *O. exaltata*/*O. scolopax*, or almost equal to $T_2$ when $T_1$ represents the speciation between *O. apifera* and *O. insectifera*.

- $T_g$ the gene-flow time of the *O. tenthredinifera* to *O. apifera* gene flow.

We therefore deduce the following order : $T_3 > T_2 > T_1 > T_g$ (or $T_3 > T_2 \geq T_1 > T_g$ if *O. insectifera* is the clade 1 taxon in the triplet).

The following results are expected with aphid for the two triplets when a gene flow occurs from *O. tenthredinifera* to *O. apifera* (expected gene trees are presented in Fig. A1):

- In triplet 1, as ILS is a random event, we expect the amount of ILS to be balanced, with an equal amount in both discordant topologies ($I_{\text{ILS}} \approx 0.5$). GF from *O. tenthredinifera* to *O. apifera* is expected to be highly significant here, with high $\Delta \ln L_{P_2P_3}$ values ($\Delta \ln L_{P_2P_3} > 3000$). Finally, as the $T_1$ speciation time between *O. apifera* and *O. exaltata*/*O. scolopax* is much more recent than $T_2$ speciation time (around twice as recent), $p_a$ is expected to be close to 0 (since speciation time $T_1$ is approximately twice as recent as $T_2$, gene flow time $T_g$ is therefore at least twice as recent as $T_2$).

- In triplet 2, similar results are expected, with $I_{\text{ILS}} \approx 0.5$ and an high $\Delta \ln L_{P_2P_3}$ values ($\Delta \ln L_{P_2P_3} > 3000$). $p_a$ value will vary depending of gene flow timing compared to $T_1$ speciation time: $p_a \approx 1$ if $T_g$ is close to $T_1$; and $p_a = 0$ if $T_g$ is at least twice as recent as $T_1$.

Similar results are expected for these two triplets if there is gene flow from *O. apifera* to *O. tenthredinifera*, rather than from *O. tenthredinifera* to *O. apifera*.

However, the results obtained using our dataset do not align with these expectations (see Supplementary Table S3):

- Triplet 1 present an high $\Delta \ln L_{P_2P_3}$ value ($\Delta \ln L_{P_2P_3} > 3000$ ) and a balance ILS amount ($I_{\text{ILS}} \approx 0.5$) but are associated with high $p_a$ value ($p_a \approx 0.87$, see Supplementary Table S4). This suggests that *O. tenthredinifera* to *O. apifera* gene flow occurred at around the same time as *O. tenthredinifera* speciation with other clade 2 taxa. This contradicts the species divergence time estimation results (Fig. 4, Supplementary Fig. S8), and suggests the following order $T_3 > T_2 > T_g > T_1$, with gene flow to *O. apifera* occurring before its speciation with its closest relative.

- Triplet 2 does not present a balanced ILS amount ($I_{\text{ILS}}$ value is significantly different from 0.5), with an excess of discordant topologies arising from ILS grouping *O. apifera* with *O. tenthredinifera*. $I_{\text{ILS}} = 0.56$ for the triplet AFB (with *O. insectifera* as clade 1 taxon), and $I_{\text{ILS}} = 0.64$ for triplets IFB and GFB (with *O. scolopax* or *O. exaltata* as clade 1 taxon). Only the triplet AFB presents a high $\Delta \ln L_{P_2P_3}$ value ($\Delta \ln L_{P_2P_3} > 3000$), associated with a high $p_a$ value ($p_a = 0.83$). Gene flow from *O. tenthredinifera* to *O. apifera* is not significant in triplets IFB and GFB ($\Delta \ln L_{P_2P_3} < 3000$).

The results obtained thus contradict Aphid expectations in the case of a gene flow from *O. tenthredinifera* to *O. apifera*. However, they are entirely consistent with Aphid expectations in the case of ghost gene flow from a taxon closely related to *O. tenthredinifera*. This 'ghost taxon' would have diverged from *O. tenthredinifera* shortly after its speciation with other clade 2 taxa. Its speciation would have occurred close to the speciation of *O. insectifera* with other clade 1 taxa, but much earlier than the speciation between between *O. apifera* and *O. exaltata*/*O. scolopax*. We then considered new divergence and gene-flow times for our two triplets:

- $T_{2'}$ the ghost speciation time between *O. tenthredinifera* and the ghost taxon. $T_{2'}$ is close to $T_2$ and close to $T_1$ when $T_1$ represents the speciation between *O. apifera* and *O. insectifera*.

- $T_{g'}$ the ghost gene flow timing between *O. apifera* and the ghost taxon (which replaces time $T_g$).

We therefore have the following order: $T_3 > T_2 \geq T_{2'} > T_1 > T_{g'}$ (or $T_3 > T_2 \geq T_{2'} \geq T_1 > T_{g'}$ if *O. insectifera* is the clade 1 taxon in the triplet). As the ghost taxon is not present in our dataset, the gene trees originating from the ghost gene flow group *O. apifera* with the ghost taxon's closest relative, *O. tenthredinifera*. We therefore incorrectly

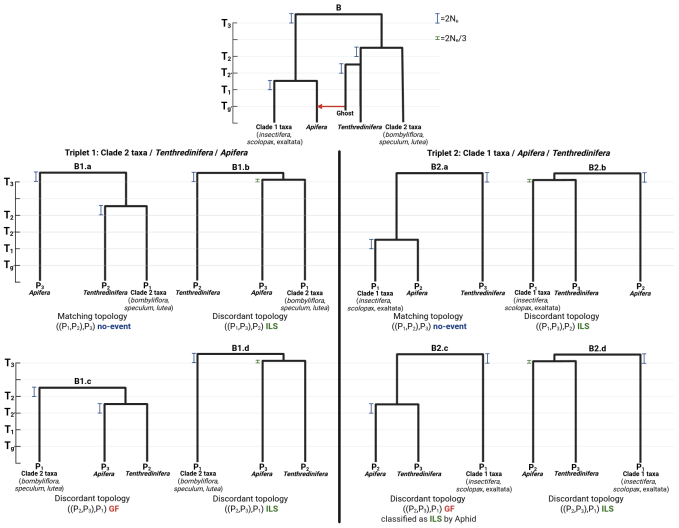


**Figure A2.** Expected gene trees in case of a ghost gene flow from a ghost taxon close to *O. tenthredinifera* to *O. apifera* (as represented in tree B), for 2 different triplets. $T_3$, $T_2$, $T_{2'}$, $T_1$ correspond to divergence times and $T_{g'}$ to gene-flow time. $N_e$ corresponds to effective population size. Red arrow reprensents gene flow event. Created in BioRender (https://BioRender.com/csvxhuj)

inferred a gene flow from *O. tenthredinifera* to *O. apifera*. The distance between *O. apifera* and *O. tenthredinifera* in gene trees originating from the ghost gene flow does not depend on the timing of the ghost gene flow $T_{g'}$, but rather on the timing of the ghost speciation $T_{2'}$. In this case, the following results are expected with aphid (expected gene trees are presented in Fig. A2):

- In triplet 1, an equal amount of ILS is expected in both discordant topologies ($I_{\text{ILS}} \approx 0.5$), with a highly significant *O. tenthredinifera* to *O. apifera* GF ($\Delta \ln L_{P_2 P_3} > 3000$). $p_a$ value is here expected to be close to 1, as $T_{2'}$ is close to $T_2$ (gene tree B1.c in Fig. A2).
- However in triplet 2, ILS is expected to be unbalanced, with an excess of ILS grouping together *O. tenthredinifera* and *O. apifera* (($I_{\text{ILS}} \neq 0.5$). Indeed, as $T_{2'} > T_1$, gene trees arising from the ghost gene flow (gene tree B2.c in Fig. A2) are wrongly assigned as ILS by aphid. As most of GF gene trees are expected to be misclassified in triplets IFB and GFB (with *O. scolopax* or *O. exaltata* as clade 1 taxon, $T_1$ is greatly lower than $T_{2'}$), we expect *O. tenthredinifera* to *O. apifera* gene flow to not be significant ($\Delta \ln L_{P_2 P_3} < 3000$) in these triplets. $I_{\text{ILS}}$ value is also expected to be more important in these triplets compared to AFB triplet. Indeed, in triplet AFB, less gene trees will be misclassified (with *O. insectifera* as clade 1 taxon, $T_1$ is close to $T_{2'}$), and *O. tenthredinifera* to *O. apifera* gene flow will possibly be significant, but associated with high $p_a$ value.
- Similar results are expected in all triplets close to triplet 2 with another clade 2 taxon than *O. tenthredinifera* (e.g. *O. bombyliflora*, *O. speculum* or *O. lutea*) as taxon $P_3$ (e.g. triplets AFD, IFD, GFD, AFE, IFE, GFE, AFC, IFC and GFC). We thus expect 12 different triplets to present an $I_{\text{ILS}}$ value significantly different from 0.5.

The observed values of $p_a$, $\Delta \ln L_{P_2 P_3}$ and $I_{\text{ILS}}$ (Supplementary Table S3, S4) for the triplets DBF, EBF, CBF, AFB, IFB, GFB, AFD, IFD, GFD, AFE, IFE, GFE, AFC, IFC and GFC in our dataset thus support a ghost gene flow from a ghost taxon closely related to *O. tenthredinifera* to *O. apifera* rather than a gene flow from *O. tenthredinifera* to *O. apifera*.

### Supplementary Figures and Tables

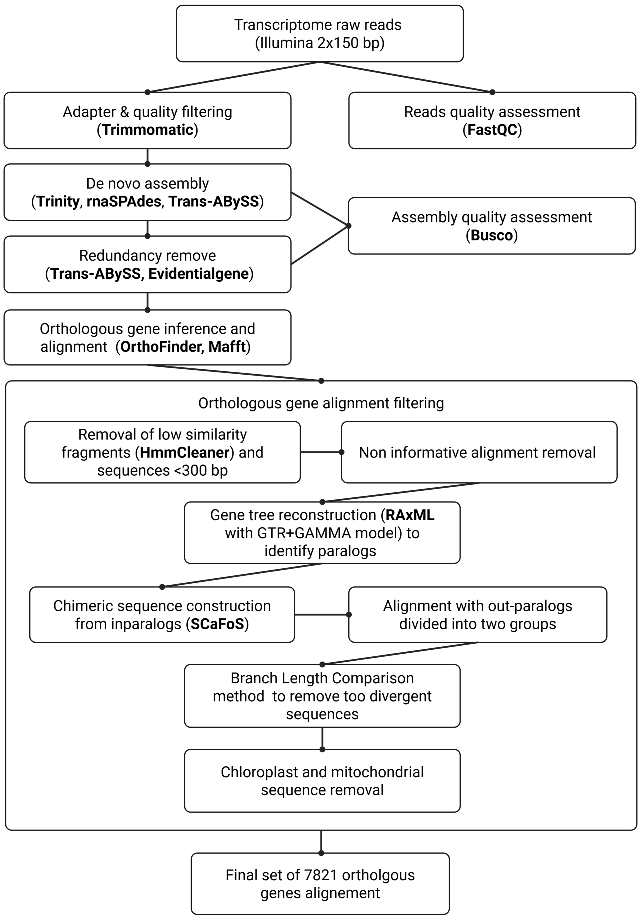


**Figure S1.** Complete pipeline to go from raw reads to orthologous gene alignments. The step 'removal of low similarity fragments (HmmCleaner) and sequences $< 300bp$' was performed until no fragments or sequences were removed. 'Informative alignments' refer to alignments with at least one sequence for each *Ophrys* and one outgroup sequence.

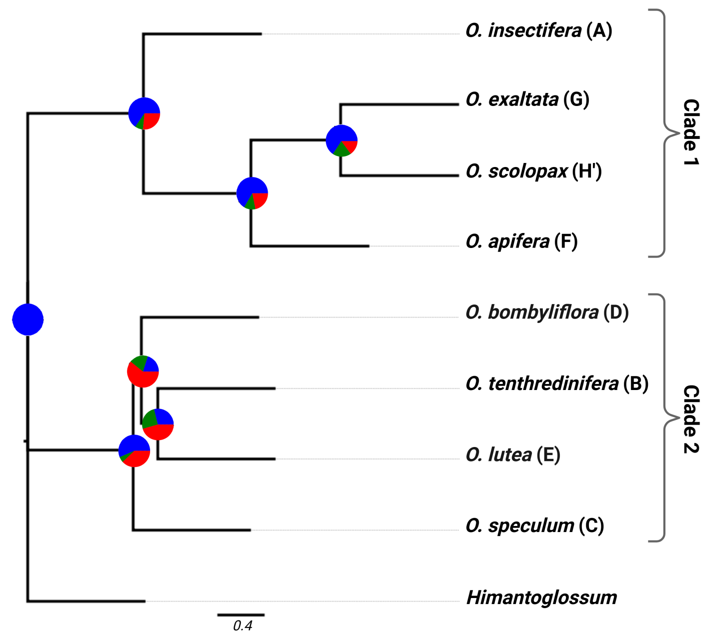


**Figure S2.** Coalescent-based tree based on 7,821 gene trees. All nodes have local posterior probabilities of 1. At each node, a pie chart depicts the proportion of gene trees supporting the species tree node (blue), the main discordant alternative node (green), or other discordant alternative nodes (red). Tree was drawn with FigTree (http://tree.bio.ed.ac.uk/software/figtree/), and pie charts were added from 'phypartspiecharts.py' results with Biorender (https://BioRender.com/8otapxc).

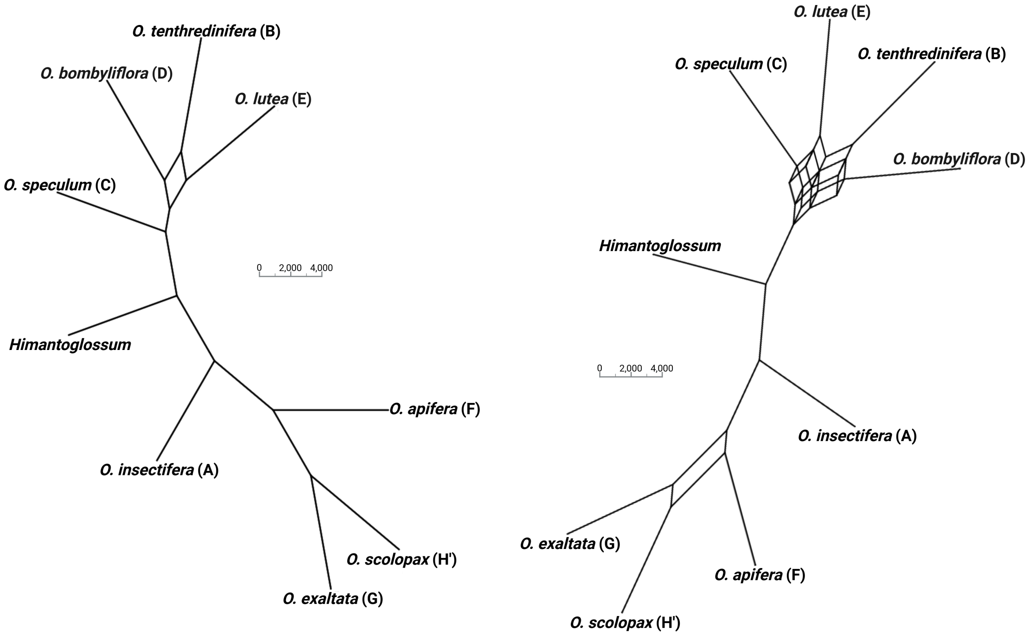


**Figure S3.** Split consensus network based on 7,821 gene trees, depicting topologies exhibited by at least 20% (on the left) or 15% (on the right) of gene trees. Branch lenghs represent the quantity of gene trees supporting the bipartitions.

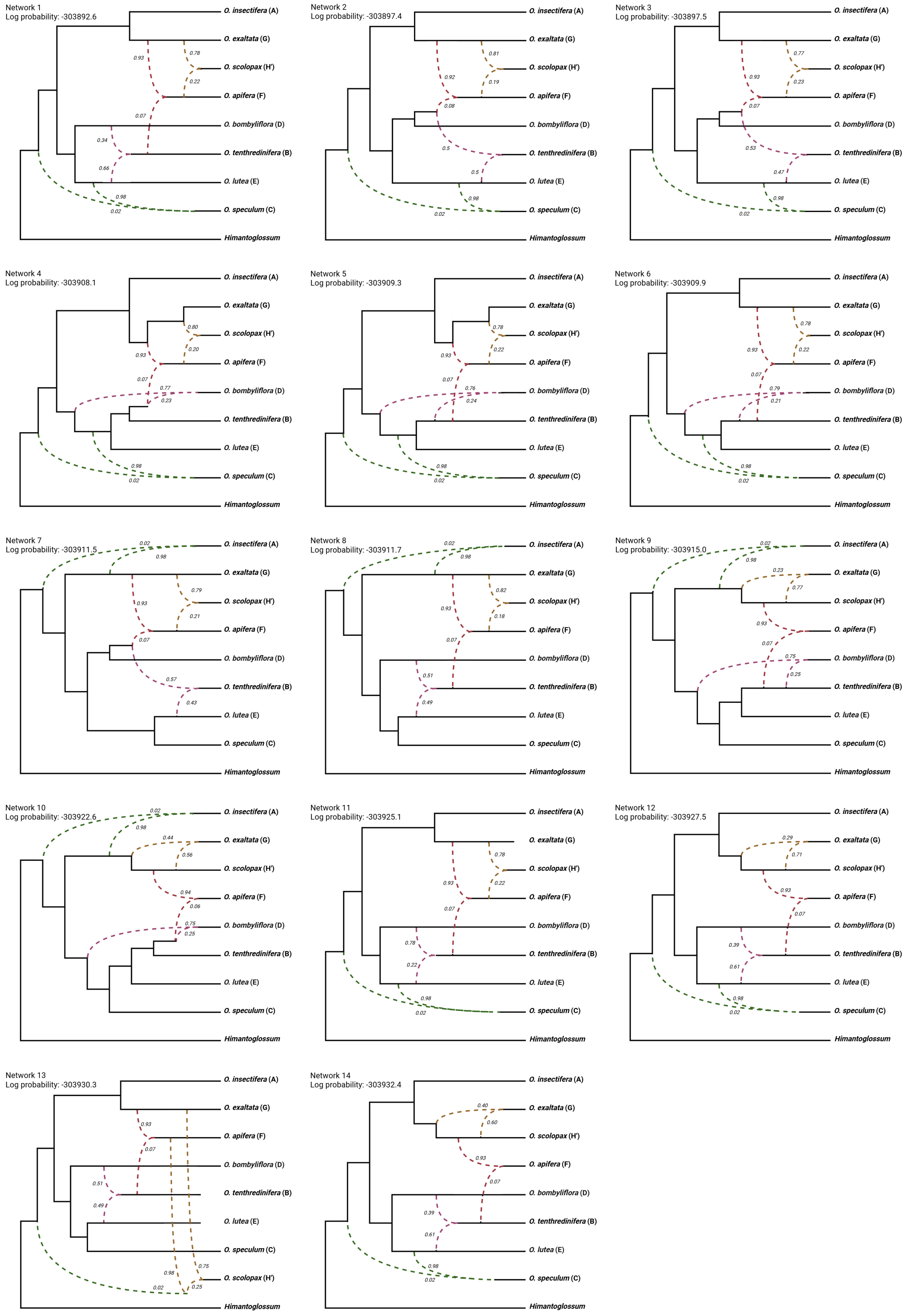


**Figure S4.** 14 PhyloNet best network with 4 reticulations according to pseudo-likelihood score without accounting for phylogenetic uncertainty. Dashed lines represent the different reticulation events, and the associated numbers the associated inheritance probabilities. Branch lengths are arbitrary.

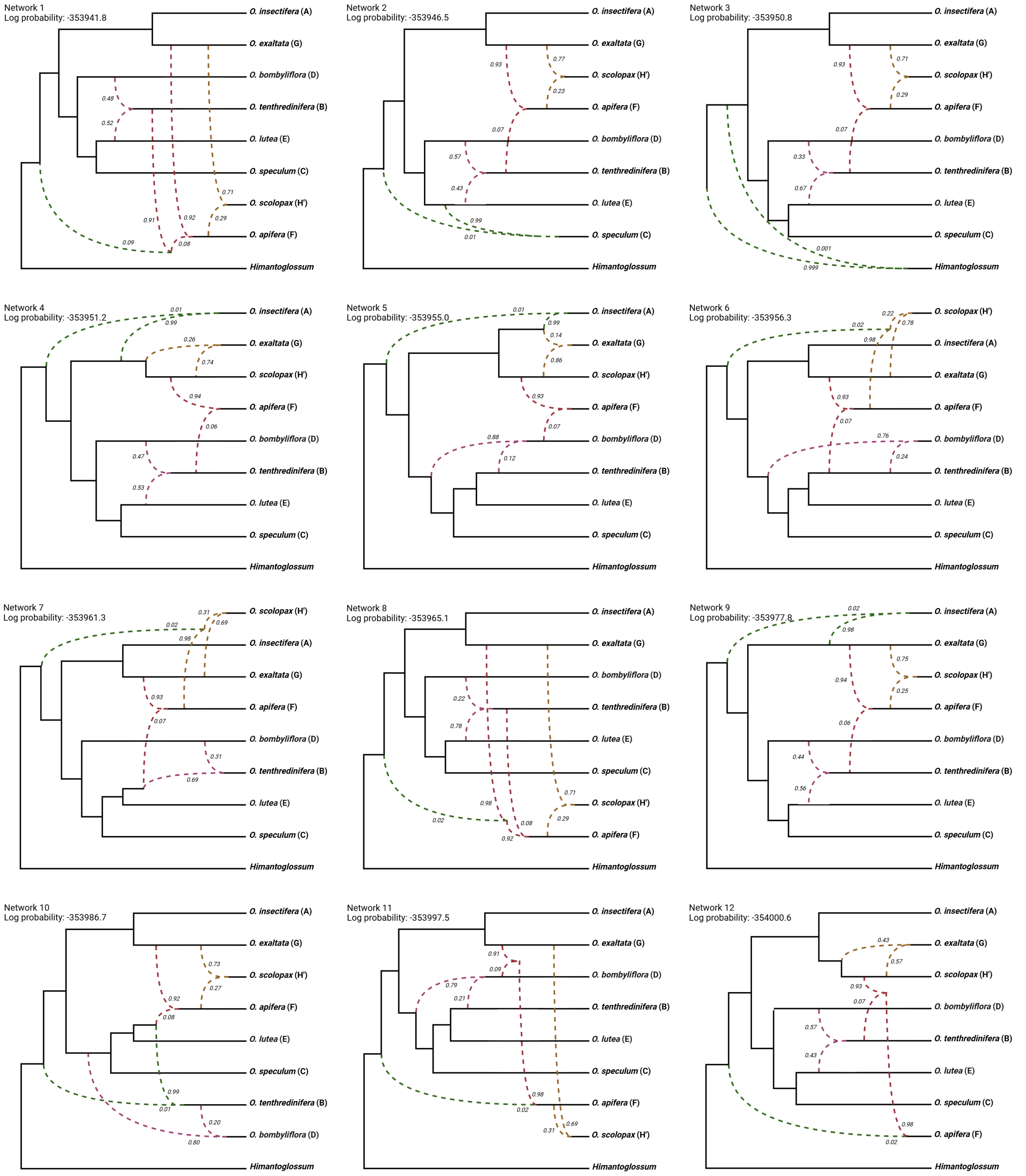


**Figure S5.** 12 PhyloNet best network with 4 reticulations according to pseudo-likelihood score when accounting for phylogenetic uncertainty. Dashed lines represent the different reticulation events, and the associated numbers the associated inheritance probabilities. Branch lengths are arbitrary.

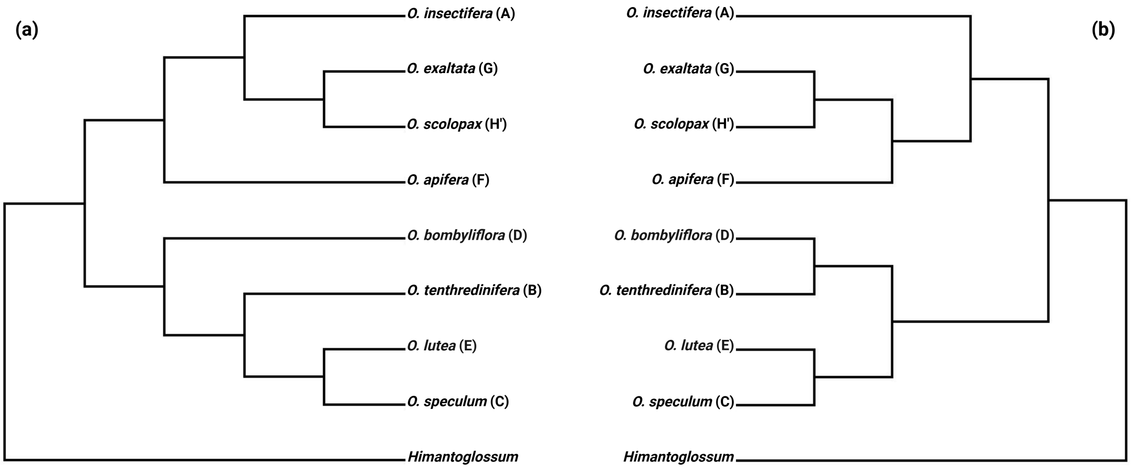


**Figure S6.** Phylogenetic network backbone trees (cladograms) visualised with Dendroscope 3. (a) Majority backbone tree: supported by 9 out of 14 best PhyloNet networks inferred without accounting for phylogenetic uncertainty, and by 9 out of 12 best networks inferred while accounting for uncertainty. (b) Minority backbone tree: supported by 5 out of 14 best PhyloNet networks inferred without accounting for phylogenetic uncertainty, and by 3 out of 12 best networks inferred while accounting for uncertainty. Branch lengths are arbitrary.

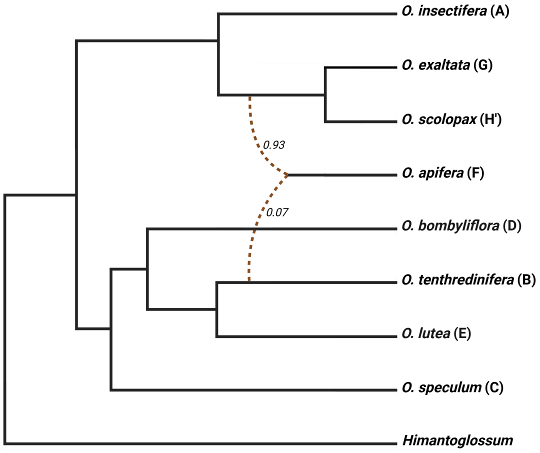


**Figure S7.** Best SNaQ network with 1 reticulation according to pseudo-likelihood score. Dashed lines represent the reticulation event, and the associated numbers the inheritance probabilities. Branch lengths are arbitrary.

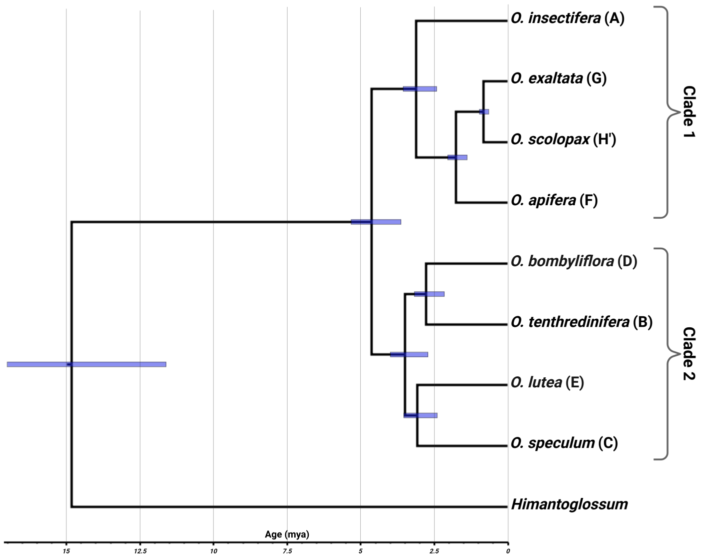


**Figure S8.** Dated network minority backbone tree inferred with mcmctree, based on 1,000 clock-like genes selected using SortaDate. The scale represents the absolute age in million years. Blue bars at nodes represent 95% highest posterior density intervals.

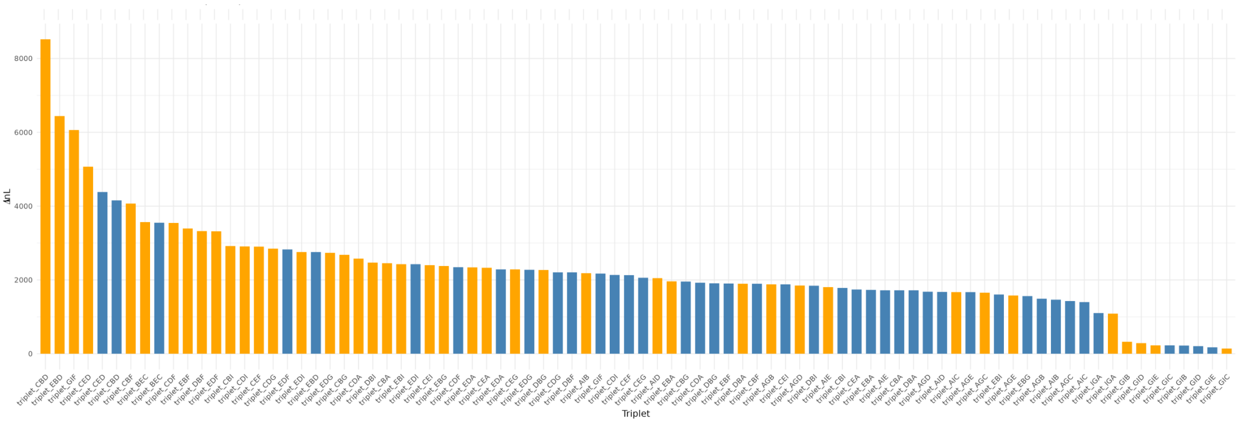


**Figure S9.** Bar plots showing, for each analysed triplet $((P_1,P_2),P_3)$ using Aphid, the difference in likelihood scores $(\Delta \ln L)$ between the full model with gene flow and the constrained model where either the $P_1$-$P_3$ gene flow probability was set to zero (blue) or the $P_2$-$P_3$ gene flow probability was set to zero (orange). Triplets with an $I_{\mathrm{ILS}}$ value significantly different from 0.5 ($p$-value $< 0.01$) are not shown.

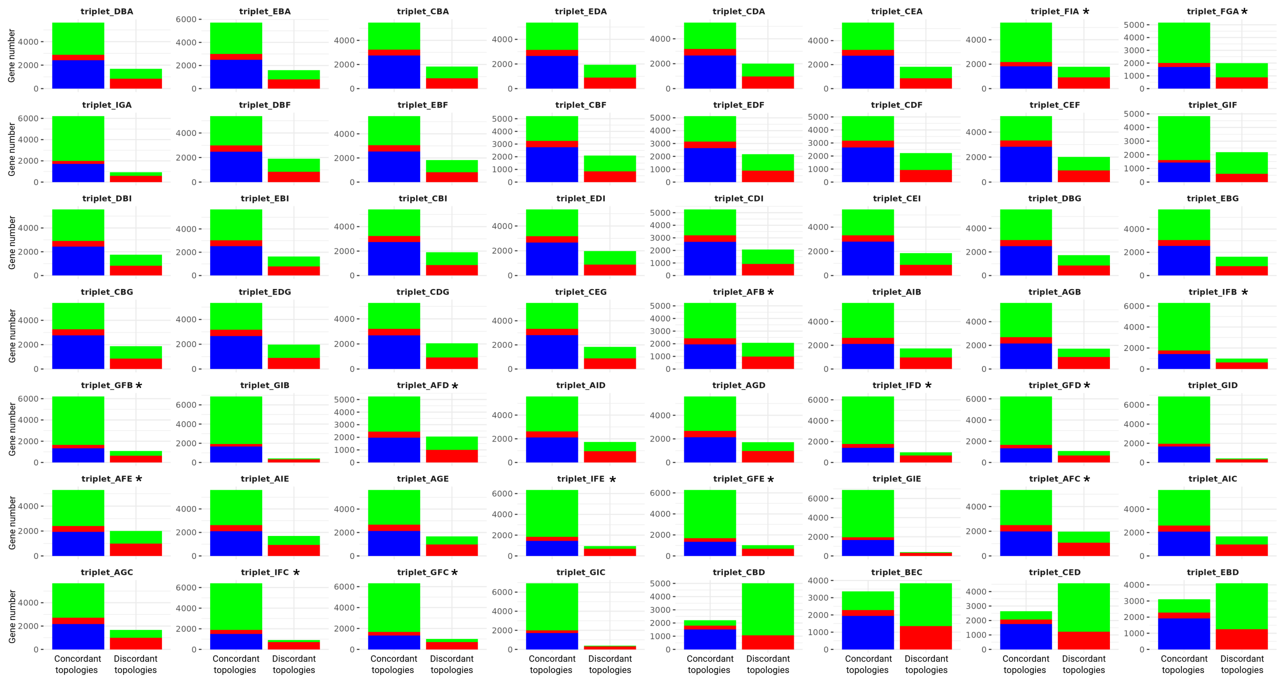


**Figure S10.** Stacked bar plots representing, for each analysed triplet $((P_1,P_2),P_3)$ using Aphid, the numbers of genes identified as mainly arising from gene flow (green), incomplete lineage sorting (red) or no-event (i.e. evolution identical to the species tree, blue). Asterisks ('*') indicate triplets filtered out by the $I_{\mathrm{ILS}}$ criterion, with an $I_{\mathrm{ILS}}$ value significantly different from 0.5 ($p$-value $< 0.01$).

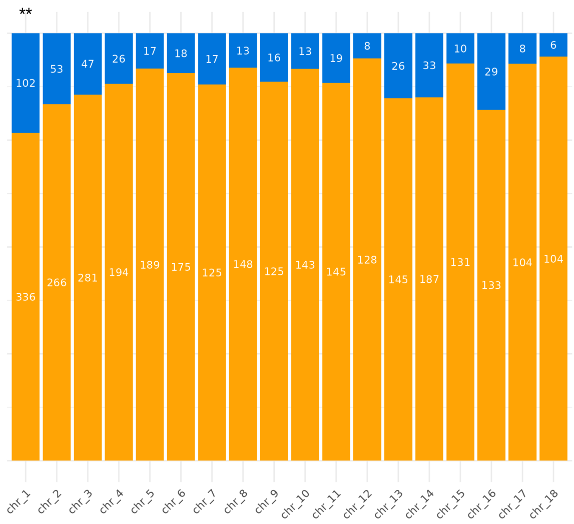


**Figure S11.** Proportion of genes per chromosome ('chr') showing potential gene flow between *O. apifera* and *O. scolopax* (in blue). Non-introgressed genes are shown in orange. Asterisks ('*') indicate chromosomes with a significant difference in the proportion of introgressed genes based on a $\chi^2$ test (: *p*-value $< 0.01$; **: *p*-value $< 0.001$).

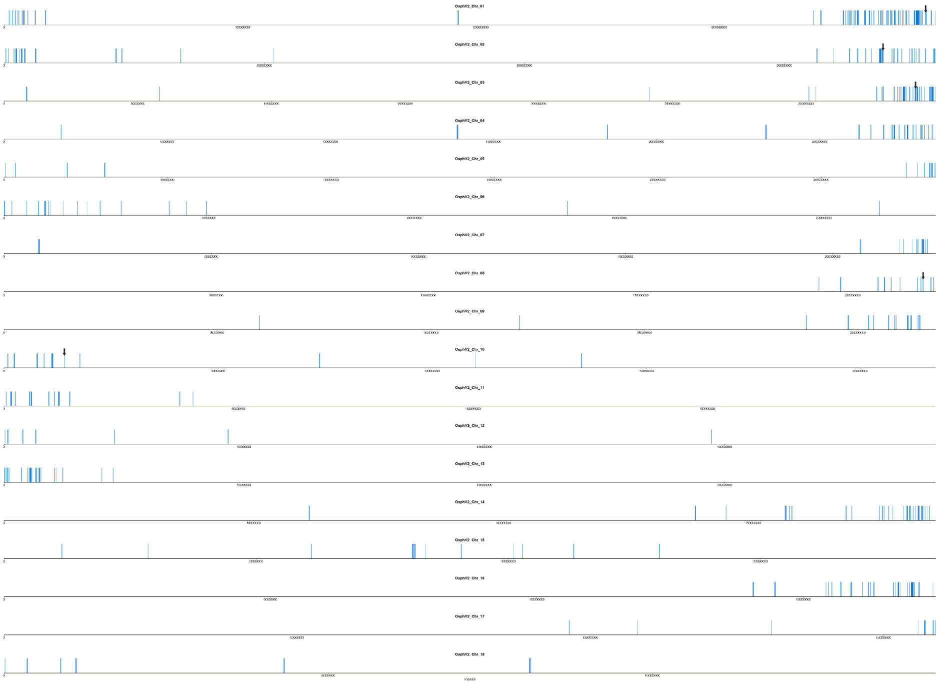

**Figure S12.** Positions along chromosomes ('OsphV2_Chr') of genes showing potential gene flow between *O. apifera* and *O. scolopax* (in blue). Black arrows indicate, among these genes, the potential key genes associated with floral phenotype.

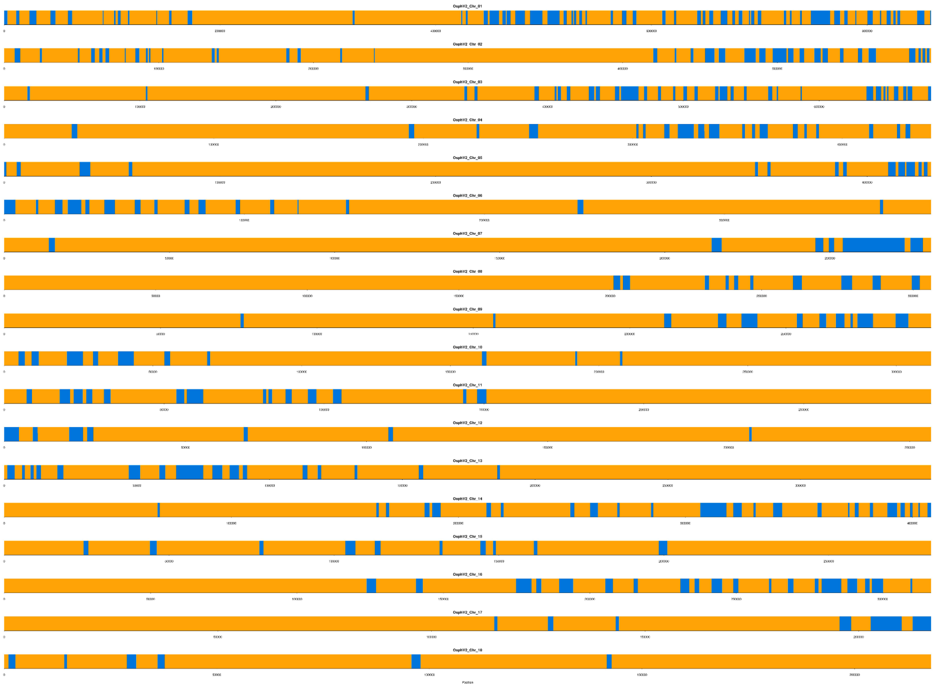

**Figure S13.** Positions along pseudo-chromosomes (‘OsphV2_Chr’) of genes showing potential gene flow between *O. apifera* and *O. scolopax* (in blue). Other genes are coloured in orange.

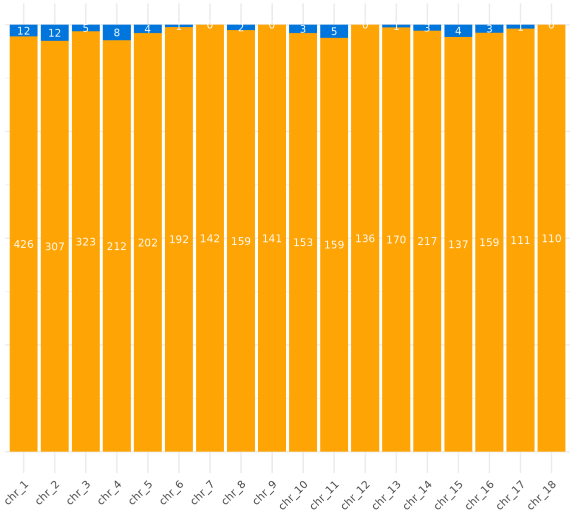


**Figure S14.** Proportion of genes per chromosome (chr’) showing potential gene flow between *O. apifera* and *O. tenthredinifera* (in blue). Non-introgressed genes are shown in orange. Asterisks (‘*’) indicate chromosomes with a significant difference in the proportion of introgressed genes based on a $\chi^2$ test (*p*-value $< 0.01$; **: *p*-value $< 0.001$).

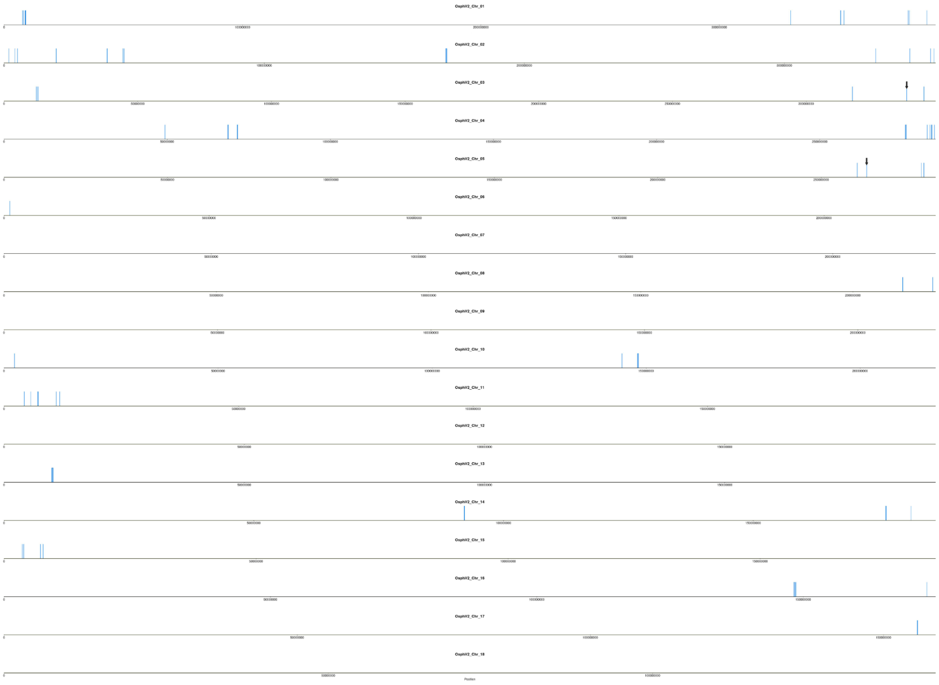

**Figure S15.** Positions along chromosomes ('OsphV2_Chr') of genes showing potential gene flow between *O. apifera* and *O. tenthredinifera* (in blue). Black arrows indicate, among these genes, the potential key genes associated with floral phenotype.

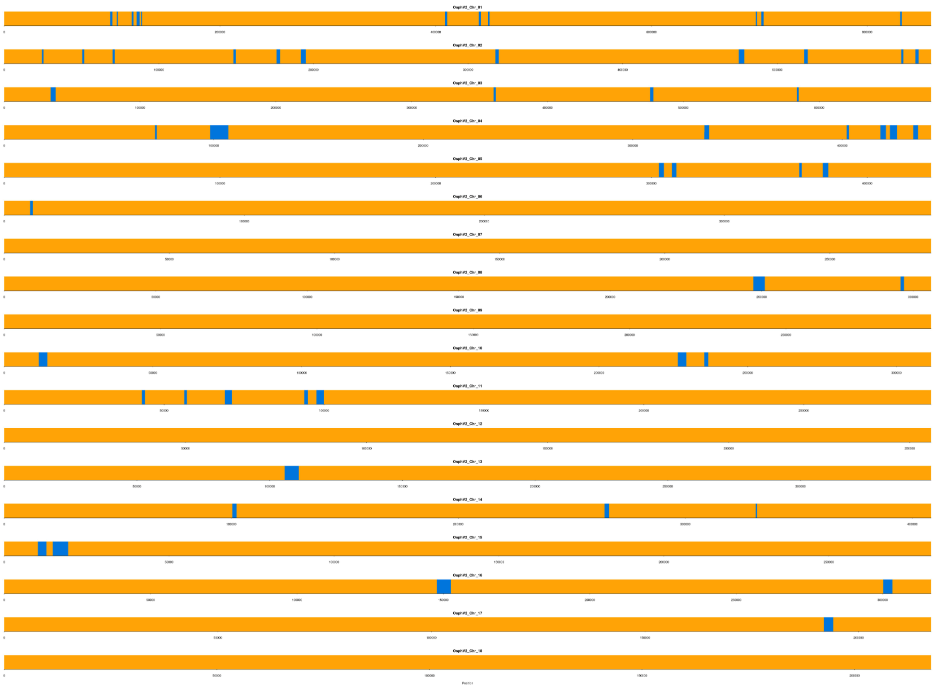

**Figure S16.** Positions along pseudo-chromosomes ('OsphV2_Chr') of genes showing potential gene flow between *O. apifera* and *O. tenthredinifera* (in blue). Other genes are coloured in orange.

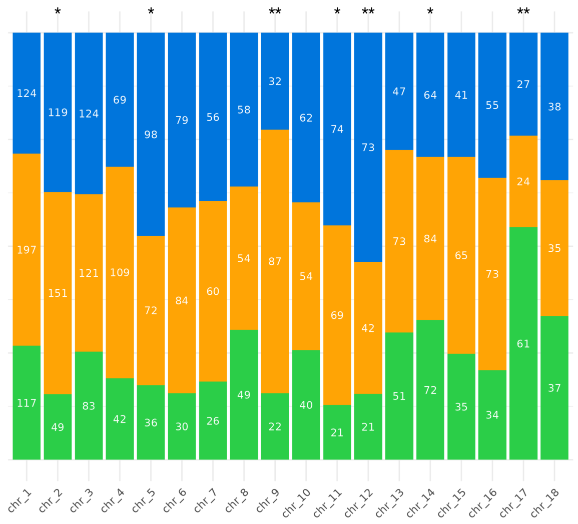


**Figure S17.** Proportion of genes per chromosome ('chr') grouping *O. tenthredinifera* and *O. lutea* (in blue) or *O. tenthredinifera* and *O. bombyliflora* (in green) as sister lineages. Other genes are shown in orange. Asterisks ('*') indicate chromosomes with a significant difference in the proportion of introgressed genes based on a $\chi^2$ test (: *p*-value $< 0.01$; **: *p*-value $< 0.001$).

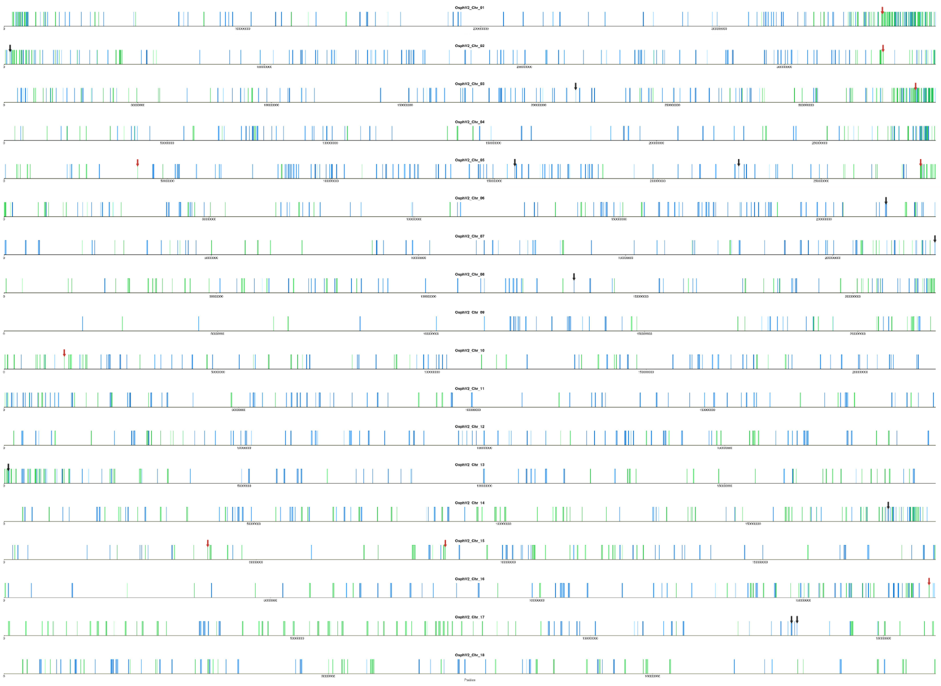

**Figure S18.** Positions along chromosomes ('OsphV2_Chr') of genes grouping *O. tenthredinifera* and *O. lutea* or *O. tenthredinifera* and *O. bombyliflora*as sister lineages.

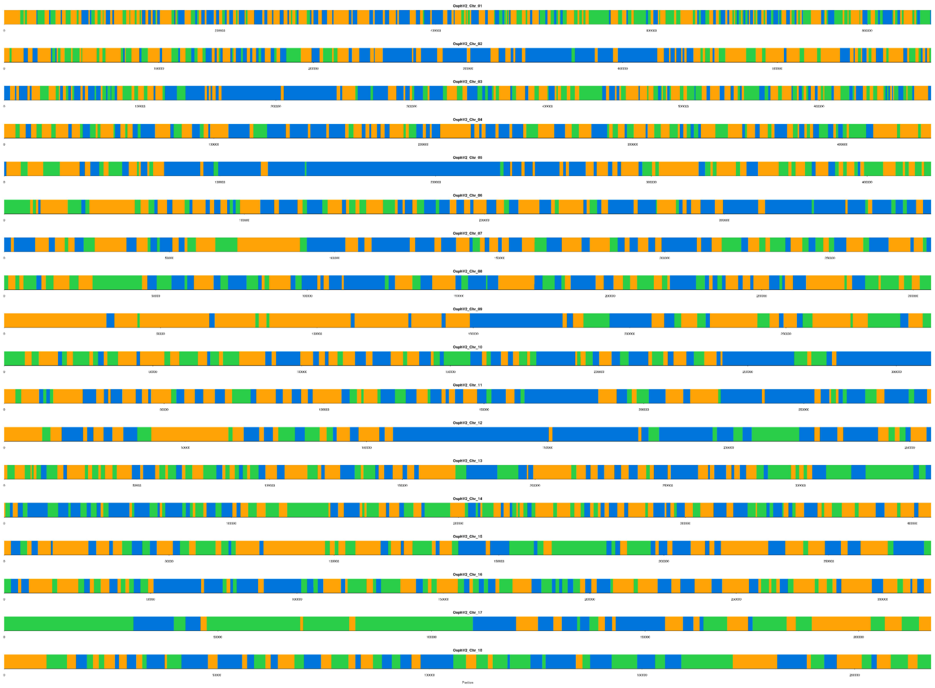

**Figure S19.** Positions along pseudo-chromosomes ('OsphV2_Chr') of genes grouping *O. tenthredinifera* and *O. lutea* (in blue) or *O. tenthredinifera* and *O. bombyliflora* (in green) as sister lineages. Other genes are coloured in orange.

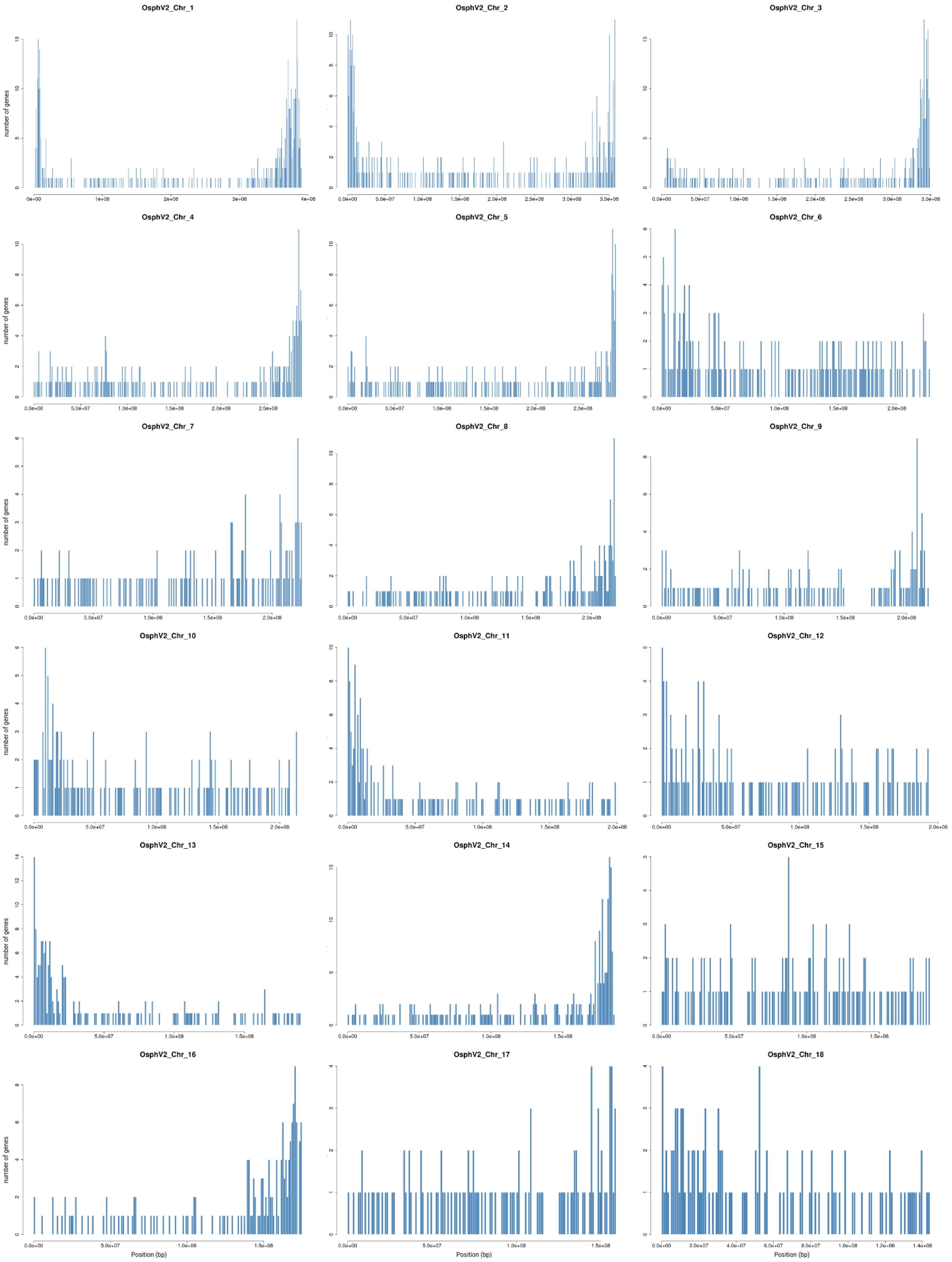


**Figure S20.** Number of genes per 1-Mb window along each chromosome (‘OsphV2_Chr’).

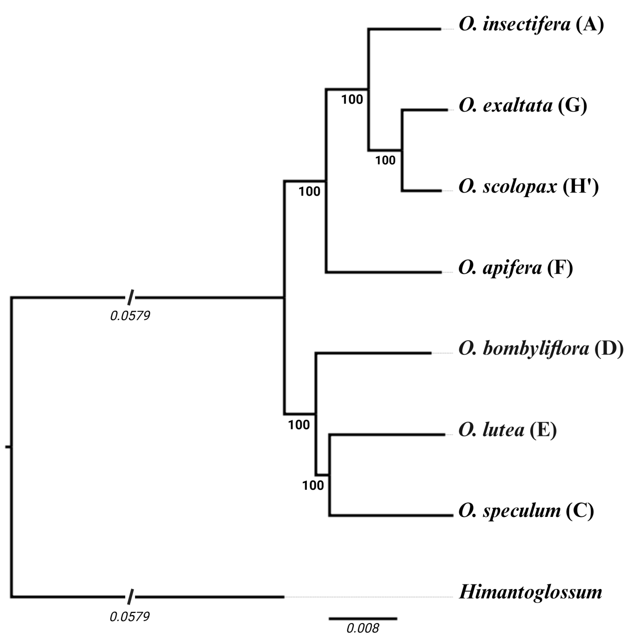


**Figure S21.** Maximum likelihood concatenation-based tree based on 7,821 orthologous gene alignments without *O. tenthredinifera*. Numbers at each node indicate bootstrap support.

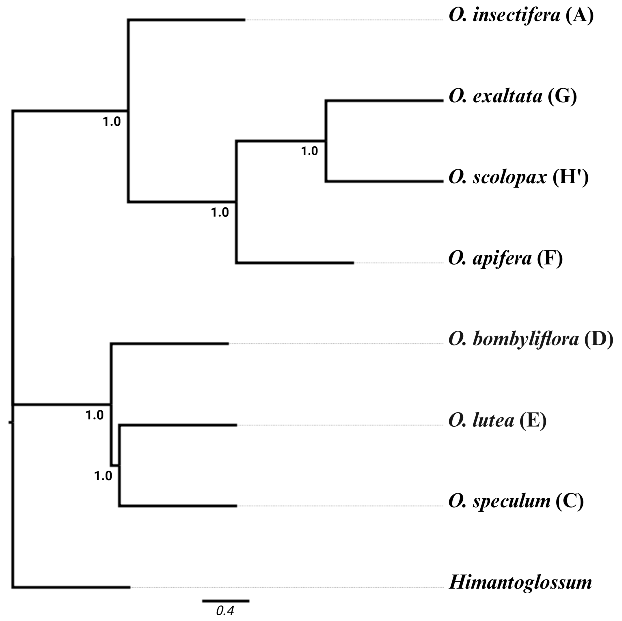


**Figure S22.** Coalescent-based tree based on 7,821 gene trees without *O. tenthredinifera*. Numbers at each node indicate local posterior probabilities.

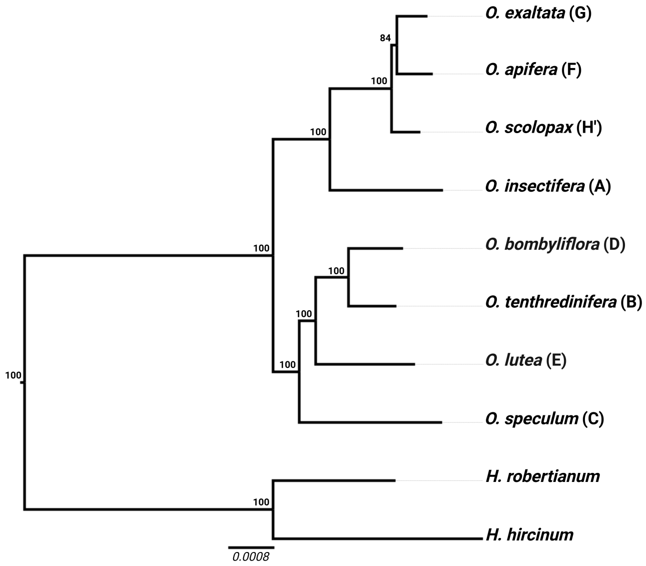


**Figure S23.** Maximum likelihood plastid tree, based on 135,806 bp and 3,013 informative positions. Numbers at each node indicate bootstrap support. Tree was drawn with FigTree (http://tree.bio.ed.ac.uk/software/figtree/).

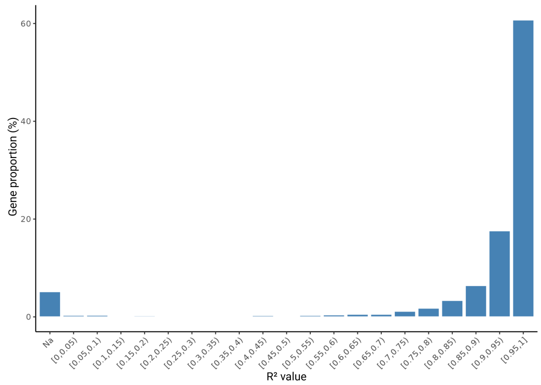


**Figure S24.** Bar plot showing the distribution of the correlation coefficient $R^2$ between branch lengths of alignment halves for each of the 7,821 genes in the final dataset. Na indicates genes for which $R^2$ could not be calculated.

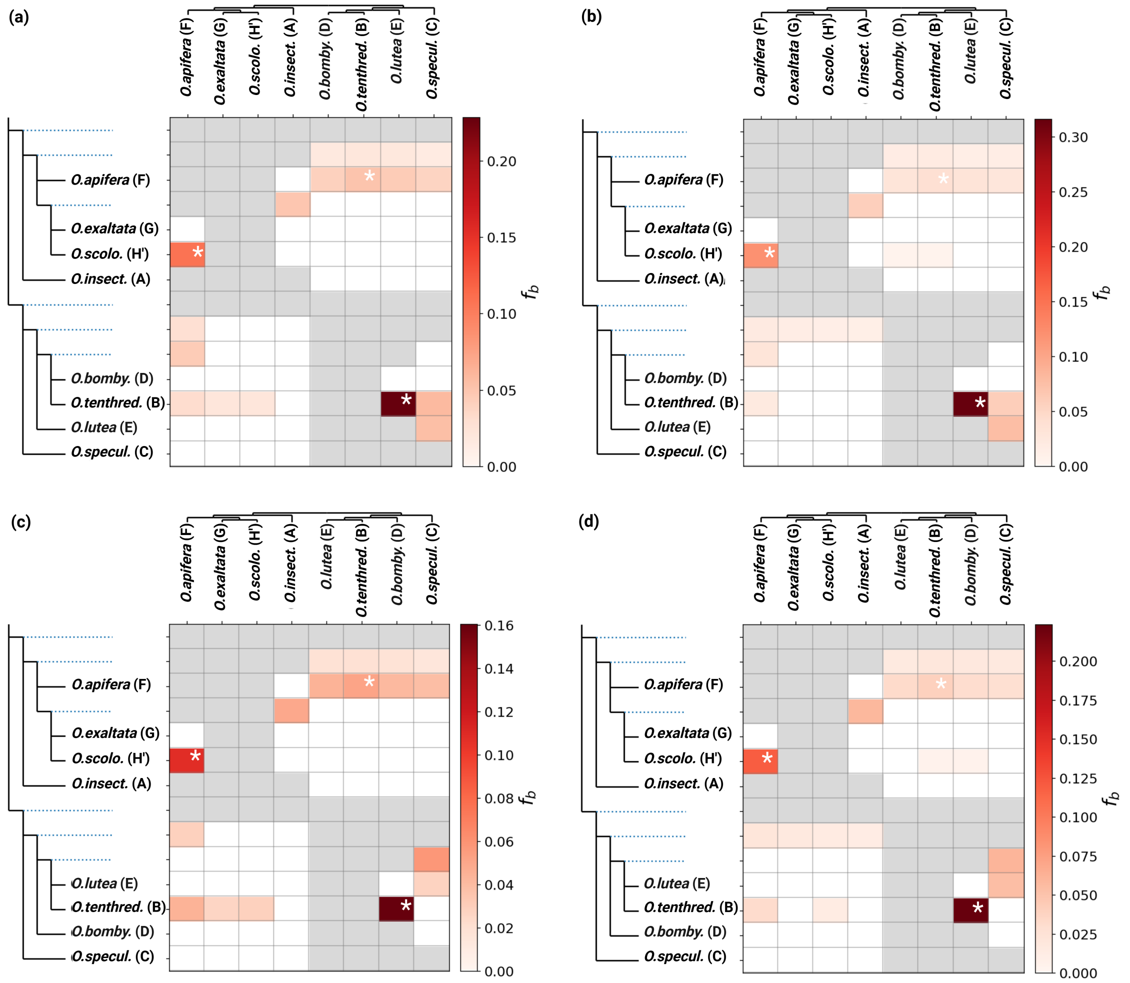


**Figure S25.** 'f branch' ('$f_b$') heatmap inferred using the concatenation-based (a & b) and coalescence-based (c & d) topology as reference, without accounting for phylogenetic uncertainty (on the left) and after keeping only nodes $\geq 70\%$ bootstrap support (on the right). Color shading represents the proportion of introgressed genes calculated for each triplet presenting a significant $\boldsymbol{\Delta}$ ($p$-value $< 0.01$). White asterisks represent the three gene flow inferred by PhyloNet.

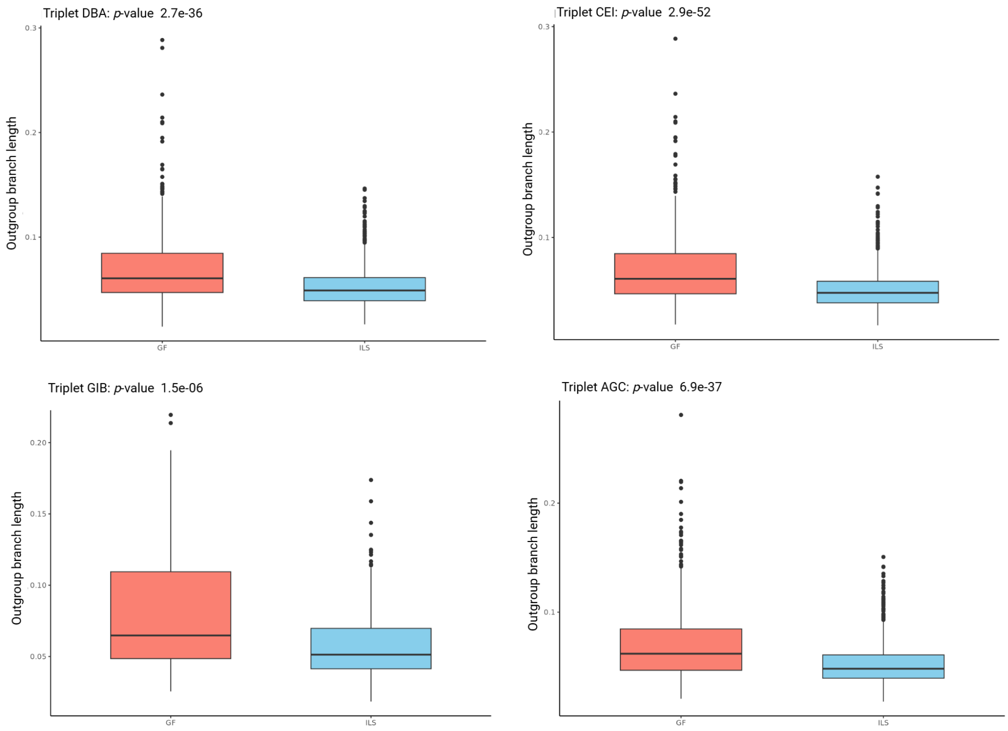


**Figure S26.** Box plots of outgroup branch lengths for discordant topologies attributed to gene flow (left) and incomplete lineage sorting (right), based on Aphid results for four different triplets. Each triplet is associated with the *p*-value from a Wilcoxon rank-sum test comparing outgroup branch lengths between gene flow and ILS.

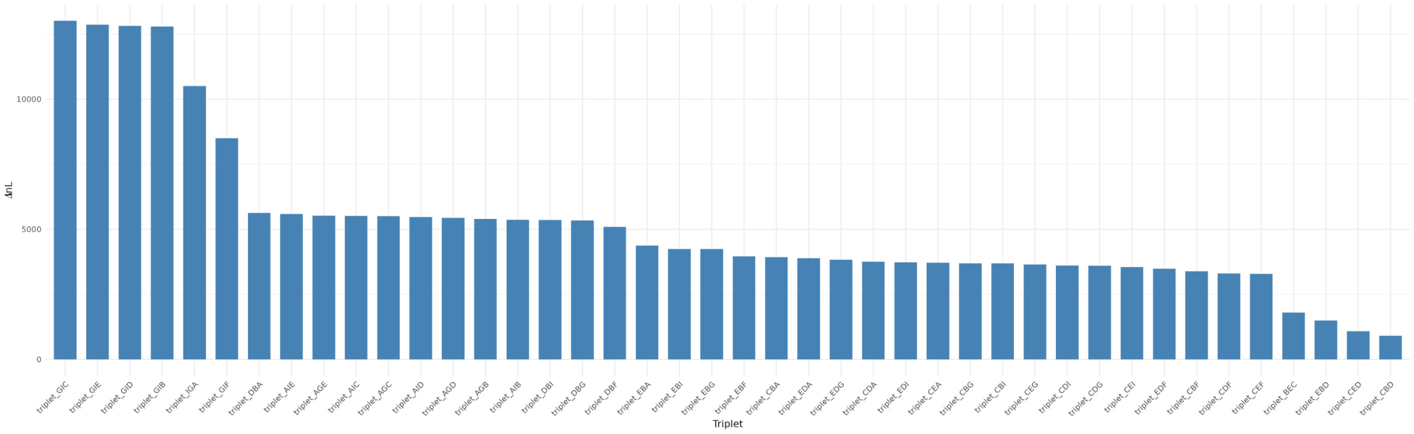


**Figure S27.** Bar plots showing, for each analysed triplet $((P_1,P_2),P_3)$ using Aphid, the difference in likelihood scores $(\Delta \ln L)$ between the full model with gene flow and the constrained model where the $P_1$-$P_2$ gene flow probability was set to zero (blue). Triplets with an $I_{\text{ILS}}$ value significantly different from 0.5 (*p*-value $< 0.01$) are not shown.

**Table S1.** Proportion of Liliopsida BUSCOs genes before and after removing redundancy for each individual.
(S): Complete and single-copy, (D): complete and duplicated, (F): fragmented, (M) missing BUSCOs genes.

| | With redundancy | | | | Without redundancy | | | |
|---|---|---|---|---|---|---|---|---|
| Species | (S) (%) | (D) (%) | (F) (%) | (M) (%) | (S) (%) | (D) (%) | (F) (%) | (M) (%) |
| *O. insectifera* | 13.1 | 74.0 | 3.3 | 9.6 | 81.9 | 2.1 | 4.6 | 11.4 |
| *O. tenthredinifera* | 12.3 | 76.0 | 3.1 | 8.6 | 82.4 | 2.7 | 4.7 | 10.2 |
| *O. bombyliflora* | 12.9 | 75.8 | 3.2 | 8.1 | 83.4 | 2.3 | 4.7 | 9.6 |
| *O. speculum* | 13.3 | 71.9 | 4.0 | 10.8 | 80.3 | 1.9 | 5.5 | 12.3 |
| *O. lutea* | 12.1 | 76.4 | 3.2 | 8.3 | 83.4 | 2.3 | 4.4 | 9.9 |
| *O. apifera* | 17.3 | 72.0 | 3.1 | 7.6 | 83.9 | 2.5 | 4.2 | 9.4 |
| *O. exaltata* | 11.2 | 78.1 | 3.2 | 7.5 | 83.8 | 2.2 | 4.7 | 9.3 |
| *O. scolopax* | 13.2 | 77.2 | 2.7 | 6.9 | 84.7 | 2.4 | 3.9 | 9.0 |
| *H. hircinum* | 14.6 | 71.9 | 4.0 | 9.5 | 81.6 | 2.2 | 5.2 | 11.0 |
| *H. robertianum* | 12.5 | 77.6 | 2.5 | 7.4 | 84.5 | 2.8 | 3.4 | 9.3 |

**Table S2a.** Δ test results summary inferred using the majority backbone tree as a reference, without accounting for phylogenetic uncertainty
For each triplet (($P_1$, $P_2$), $P_3$), the number of matching topologies (BBAA) and discordant topologies (ABBA and BABA) are reported.
The Δ statistic, with Δ = (ABBA − BABA) / (ABBA + BABA), and associated *p* -values and Benjamini–Hochberg corrected *p* -values are provided.
The proportion of introgressed genes (γ) is also reported, with γ = (ABBA − BABA) / (ABBA + BABA + BBAA).

| Triplet ID | $P_1$ | $P_2$ | $P_3$ | Δ statistic | *p* -value | corrected p-value | γ | BBAA | ABBA | BABA |
|---|---|---|---|---|---|---|---|---|---|---|
| DBA | *Obombyliflora* | *Otenthredinifera* | *Oinsectifera* | 0,06149584 | 0,009285735 | 0,01209305 | | 6016 | 958 | 847 |
| EBA | *Olutea* | *Otenthredinifera* | *Oinsectifera* | 0,06387921 | 0,00814037 | 0,011060572 | | 6099 | 916 | 806 |
| CBA | *Ospeculum* | *Otenthredinifera* | *Oinsectifera* | 0,1056338 | 3,43E-06 | 7,11E-06 | 0,026850786 | 5833 | 1099 | 889 |
| EDA | *Obombyliflora* | *Olutea* | *Oinsectifera* | 0,00714626 | 0,74701 | 0,760592 | | 5722 | 1057 | 1042 |
| CDA | *Ospeculum* | *Obombyliflora* | *Oinsectifera* | 0,06555863 | 0,00232072 | 0,003420008 | 0,018156246 | 5655 | 1154 | 1012 |
| CEA | *Ospeculum* | *Olutea* | *Oinsectifera* | 0,03922567 | 0,08628044 | 0,102802226 | | 5858 | 1020 | 943 |
| FIA | *Oapifera* | *Oscolopax* | *Oinsectifera* | 0,2130638 | 0 | 0 | 0,052550825 | 5892 | 1170 | 759 |
| FGA | *Oapifera* | *Oexaltata* | *Oinsectifera* | 0,1781951 | 0 | 0 | 0,049737885 | 5638 | 1286 | 897 |
| IGA | *Oscolopax* | *Oexaltata* | *Oinsectifera* | 0,003952569 | 0,8979873 | 0,8979873 | | 6809 | 508 | 504 |
| DBF | *Obombyliflora* | *Otenthredinifera* | *Oapifera* | 0,1200768 | 8,79E-08 | 2,14E-07 | 0,031965222 | 5739 | 1166 | 916 |
| EBF | *Olutea* | *Otenthredinifera* | *Oapifera* | 0,1780822 | 7,33E-15 | 2,16E-14 | 0,044879171 | 5850 | 1161 | 810 |
| CBF | *Ospeculum* | *Otenthredinifera* | *Oapifera* | 0,1998239 | 0 | 0 | 0,058048843 | 5549 | 1363 | 909 |
| EDF | *Olutea* | *Obombyliflora* | *Oapifera* | 0,0388391 | 0,058237 | 0,070897217 | | 5478 | 1217 | 1126 |
| CDF | *Ospeculum* | *Obombyliflora* | *Oapifera* | 0,09399586 | 2,26E-06 | 4,87E-06 | 0,029024421 | 5406 | 1321 | 1094 |
| CEF | *Ospeculum* | *Olutea* | *Oapifera* | 0,05354691 | 0,01609636 | 0,020031026 | | 5636 | 1151 | 1034 |
| GIF | *Oexaltata* | *Oscolopax* | *Oapifera* | 0,3478798 | 0 | 0 | 0,10804245 | 5392 | 1637 | 792 |
| DBI | *Obombyliflora* | *Otenthredinifera* | *Oscolopax* | 0,08762613 | 0,000101297 | 0,000189087 | 0,021097046 | 5938 | 1024 | 859 |
| EBI | *Olutea* | *Otenthredinifera* | *Oscolopax* | 0,1295455 | 1,36E-08 | 3,63E-08 | 0,029152282 | 6061 | 994 | 766 |
| CBI | *Ospeculum* | *Otenthredinifera* | *Oscolopax* | 0,1461875 | 1,22E-11 | 3,42E-11 | 0,038486127 | 5762 | 1180 | 879 |
| EDI | *Olutea* | *Obombyliflora* | *Oscolopax* | 0,03135236 | 0,1461298 | 0,163665376 | | 5684 | 1102 | 1035 |
| CDI | *Ospeculum* | *Obombyliflora* | *Oscolopax* | 0,07352278 | 0,000468972 | 0,000772424 | 0,020841325 | 5604 | 1190 | 1027 |
| CEI | *Ospeculum* | *Olutea* | *Oscolopax* | 0,03441397 | 0,1156188 | 0,132135771 | | 5816 | 1037 | 968 |
| DBG | *Obombyliflora* | *Otenthredinifera* | *Oexaltata* | 0,08866201 | 0,000208279 | 0,000376246 | 0,021097046 | 5960 | 1013 | 848 |
| EBG | *Olutea* | *Otenthredinifera* | *Oexaltata* | 0,1191294 | 6,88E-07 | 1,61E-06 | 0,026595065 | 6075 | 977 | 769 |
| CBG | *Ospeculum* | *Otenthredinifera* | *Oexaltata* | 0,121063 | 7,68E-08 | 1,95E-07 | 0,031453778 | 5789 | 1139 | 893 |
| EDG | *Olutea* | *Obombyliflora* | *Oexaltata* | 0,02247191 | 0,2938133 | 0,310444242 | | 5685 | 1092 | 1044 |
| CDG | *Ospeculum* | *Obombyliflora* | *Oexaltata* | 0,06079855 | 0,004263458 | 0,006121888 | 0,017133359 | 5617 | 1169 | 1035 |
| CEG | *Ospeculum* | *Olutea* | *Oexaltata* | 0,02802803 | 0,2212189 | 0,242907028 | | 5823 | 1027 | 971 |
| AFB | *Oinsectifera* | *Oapifera* | *Otenthredinifera* | 0,2703554 | 0 | 0 | 0,076844393 | 5598 | 1412 | 811 |
| AIB | *Oinsectifera* | *Oscolopax* | *Otenthredinifera* | 0,1067024 | 4,24E-06 | 8,48E-06 | 0,025444317 | 5956 | 1032 | 833 |
| AGB | *Oinsectifera* | *Oexaltata* | *Otenthredinifera* | 0,08297414 | 0,000571683 | 0,000914692 | 0,019690577 | 5965 | 1005 | 851 |
| IFB | *Oscolopax* | *Oapifera* | *Otenthredinifera* | 0,3527205 | 0 | 0 | 0,048075694 | 6755 | 721 | 345 |
| GFB | *Oexaltata* | *Oapifera* | *Otenthredinifera* | 0,3663535 | 0 | 0 | 0,054852321 | 6650 | 800 | 371 |
| GIB | *Oexaltata* | *Oscolopax* | *Otenthredinifera* | 0,07317073 | 0,1078016 | 0,125768533 | | 7370 | 242 | 209 |
| AFD | *Oinsectifera* | *Oapifera* | *Obombyliflora* | 0,2409091 | 0 | 0 | 0,06776627 | 5621 | 1365 | 835 |
| AID | *Oinsectifera* | *Oscolopax* | *Obombyliflora* | 0,1034483 | 1,15E-05 | 2,22E-05 | 0,02454929 | 5965 | 1024 | 832 |
| AGD | *Oinsectifera* | *Oexaltata* | *Obombyliflora* | 0,07483731 | 0,001534602 | 0,002322641 | 0,017644802 | 5977 | 991 | 853 |
| IFD | *Oscolopax* | *Oapifera* | *Obombyliflora* | 0,2836676 | 0 | 0 | 0,037974684 | 6774 | 672 | 375 |
| GFD | *Oexaltata* | *Oapifera* | *Obombyliflora* | 0,3067591 | 0 | 0 | 0,045262754 | 6667 | 754 | 400 |
| GID | *Oexaltata* | *Oscolopax* | *Obombyliflora* | 0,1145374 | 0,0128596 | 0,016366764 | | 7367 | 253 | 201 |
| AFE | *Oinsectifera* | *Oapifera* | *Olutea* | 0,232634 | 0 | 0 | 0,063802583 | 5676 | 1322 | 823 |
| AIE | *Oinsectifera* | *Oscolopax* | *Olutea* | 0,08140814 | 0,000352723 | 0,00059856 | 0,018923411 | 6003 | 983 | 835 |
| AGE | *Oinsectifera* | *Oexaltata* | *Olutea* | 0,06364139 | 0,008295429 | 0,011060572 | | 6014 | 961 | 846 |
| IFE | *Oscolopax* | *Oapifera* | *Olutea* | 0,3346457 | 0 | 0 | 0,043472702 | 6805 | 678 | 338 |
| GFE | *Oexaltata* | *Oapifera* | *Olutea* | 0,3279133 | 0 | 0 | 0,046413502 | 6714 | 735 | 372 |
| GIE | *Oscolopax* | *Oexaltata* | *Olutea* | 0,05479452 | 0,246076 | 0,265004923 | | 7383 | 232 | 207 |
| AFC | *Oinsectifera* | *Oapifera* | *Ospeculum* | 0,2145558 | 0 | 0 | 0,058048843 | 5705 | 1285 | 831 |
| AIC | *Oinsectifera* | *Oscolopax* | *Ospeculum* | 0,08467072 | 0,000341763 | 0,000598085 | 0,019562716 | 6014 | 980 | 827 |
| AGC | *Oinsectifera* | *Oexaltata* | *Ospeculum* | 0,06489185 | 0,005166241 | 0,007232737 | 0,014959724 | 6018 | 960 | 843 |
| IFC | *Oscolopax* | *Oapifera* | *Ospeculum* | 0,299075 | 0 | 0 | 0,037207518 | 6848 | 632 | 341 |
| GFC | *Oexaltata* | *Oapifera* | *Ospeculum* | 0,2943361 | 0 | 0 | 0,040531901 | 6744 | 697 | 380 |
| GIC | *Oexaltata* | *Oscolopax* | *Ospeculum* | 0,04895105 | 0,3128859 | 0,324474267 | | 7392 | 225 | 204 |
| CBD | *Ospeculum* | *Otenthredinifera* | *Obombyliflora* | 0,265912 | 0 | 0 | 0,183224652 | 2432 | 3411 | 1978 |
| BEC | *Otenthredinifera* | *Olutea* | *Ospeculum* | 0,05215311 | 0,00103879 | 0,001615896 | 0,027873673 | 3641 | 2199 | 1981 |
| CED | *Ospeculum* | *Olutea* | *Obombyliflora* | 0,06998572 | 7,84E-07 | 1,76E-06 | 0,043856284 | 2920 | 2622 | 2279 |
| EBD | *Olutea* | *Otenthredinifera* | *Obombyliflora* | 0,2823053 | 0 | 0 | 0,160337553 | 3379 | 2848 | 1594 |

**Table S2b.** Δ test results summary inferred using the majority backbone tree as a reference, after retaining only nodes with ≥70% bootstrap support.
For each triplet (($P_1$, $P_2$), $P_3$), the number of matching topologies (BBAA) and discordant topologies (ABBA and BABA) are reported.
The Δ statistic, with Δ = (ABBA − BABA) / (ABBA + BABA), and associated *p* -values and Benjamini–Hochberg corrected *p* -values are provided.
The proportion of introgressed genes (γ) is also reported, with γ = (ABBA − BABA) / (ABBA + BABA + BBAA).

| Triplet ID | $P_1$ | $P_2$ | $P_3$ | Δ statistic | *p* -value | corrected p-value | γ | BBAA | ABBA | BABA |
|---|---|---|---|---|---|---|---|---|---|---|
| DBA | *Obombyliflora* | *Otenthredinifera* | *Oinsectifera* | 0,02233251 | 0,6636916 | 0,6636916 | | 4068 | 206 | 197 |
| EBA | *Olutea* | *Otenthredinifera* | *Oinsectifera* | 0,05882353 | 0,2493488 | 0,273794761 | | 4161 | 207 | 184 |
| CBA | *Ospeculum* | *Otenthredinifera* | *Oinsectifera* | 0,1990741 | 2,56E-05 | 4,94E-05 | 0,020481067 | 3767 | 259 | 173 |
| EDA | *Obombyliflora* | *Olutea* | *Oinsectifera* | 0,03643725 | 0,4203668 | 0,435935941 | | 3683 | 256 | 238 |
| CDA | *Ospeculum* | *Obombyliflora* | *Oinsectifera* | 0,1779661 | 7,13E-05 | 0,000124775 | 0,020563036 | 3613 | 278 | 194 |
| CEA | *Ospeculum* | *Olutea* | *Oinsectifera* | 0,1306413 | 0,006937245 | 0,009034552 | 0,013036265 | 3798 | 238 | 183 |
| FIA | *Oapifera* | *Oscolopax* | *Oinsectifera* | 0,3460439 | 0 | 0 | 0,060139165 | 4987 | 706 | 343 |
| FGA | *Oapifera* | *Oexaltata* | *Oinsectifera* | 0,2953191 | 0 | 0 | 0,058963466 | 4710 | 761 | 414 |
| IGA | *Oscolopax* | *Oexaltata* | *Oinsectifera* | 0,03265306 | 0,4705291 | 0,479084175 | | 6161 | 253 | 237 |
| DBF | *Obombyliflora* | *Otenthredinifera* | *Oapifera* | 0,1754685 | 1,79E-05 | 3,58E-05 | 0,023224352 | 3848 | 345 | 242 |
| EBF | *Olutea* | *Otenthredinifera* | *Oapifera* | 0,2608696 | 3,11E-10 | 9,17E-10 | 0,031879566 | 3965 | 348 | 204 |
| CBF | *Ospeculum* | *Otenthredinifera* | *Oapifera* | 0,3892617 | 0 | 0 | 0,055729042 | 3567 | 414 | 182 |
| EDF | *Olutea* | *Obombyliflora* | *Oapifera* | 0,06200318 | 0,1095305 | 0,127785583 | | 3483 | 334 | 295 |
| CDF | *Ospeculum* | *Obombyliflora* | *Oapifera* | 0,2125206 | 1,13E-07 | 2,88E-07 | 0,032049689 | 3418 | 368 | 239 |
| CEF | *Ospeculum* | *Olutea* | *Oapifera* | 0,1714836 | 3,19E-05 | 5,95E-05 | 0,021533995 | 3614 | 304 | 215 |
| GIF | *Oexaltata* | *Oscolopax* | *Oapifera* | 0,4778068 | 0 | 0 | 0,119764398 | 4580 | 1132 | 400 |
| DBI | *Obombyliflora* | *Otenthredinifera* | *Oscolopax* | 0,09450549 | 0,04279654 | 0,050991622 | | 4016 | 249 | 206 |
| EBI | *Olutea* | *Otenthredinifera* | *Oscolopax* | 0,144186 | 0,002004916 | 0,003034467 | 0,013614405 | 4124 | 246 | 184 |
| CBI | *Ospeculum* | *Otenthredinifera* | *Oscolopax* | 0,2700422 | 6,50E-10 | 1,82E-09 | 0,03047619 | 3726 | 301 | 173 |
| EDI | *Olutea* | *Obombyliflora* | *Oscolopax* | 0,04743833 | 0,2719458 | 0,292864708 | | 3639 | 276 | 251 |
| CDI | *Ospeculum* | *Obombyliflora* | *Oscolopax* | 0,180198 | 3,72E-05 | 6,72E-05 | 0,02231486 | 3573 | 298 | 207 |
| CEI | *Ospeculum* | *Olutea* | *Oscolopax* | 0,1390135 | 0,003374727 | 0,00462669 | 0,014751368 | 3757 | 254 | 192 |
| DBG | *Obombyliflora* | *Otenthredinifera* | *Oexaltata* | 0,06912442 | 0,1406727 | 0,1607688 | | 4039 | 232 | 202 |
| EBG | *Olutea* | *Otenthredinifera* | *Oexaltata* | 0,1259259 | 0,01323021 | 0,016838449 | | 4144 | 228 | 177 |
| CBG | *Ospeculum* | *Otenthredinifera* | *Oexaltata* | 0,2044444 | 1,85E-06 | 4,32E-06 | 0,02193087 | 3745 | 271 | 179 |
| EDG | *Olutea* | *Obombyliflora* | *Oexaltata* | 0,04230769 | 0,3397586 | 0,358990219 | | 3657 | 271 | 249 |
| CDG | *Ospeculum* | *Obombyliflora* | *Oexaltata* | 0,1371769 | 0,003123046 | 0,004484374 | 0,016866292 | 3588 | 286 | 217 |
| CEG | *Ospeculum* | *Olutea* | *Oexaltata* | 0,09954751 | 0,03259651 | 0,039682708 | | 3768 | 243 | 199 |
| AFB | *Oinsectifera* | *Oapifera* | *Otenthredinifera* | 0,4349355 | 0 | 0 | 0,071815718 | 4313 | 612 | 241 |
| AIB | *Oinsectifera* | *Oscolopax* | *Otenthredinifera* | 0,2195122 | 3,73E-08 | 9,95E-08 | 0,025714286 | 4635 | 375 | 240 |
| AGB | *Oinsectifera* | *Oexaltata* | *Otenthredinifera* | 0,1710526 | 1,67E-05 | 3,46E-05 | 0,019790676 | 4647 | 356 | 252 |
| IFB | *Oscolopax* | *Oapifera* | *Otenthredinifera* | 0,4887526 | 0 | 0 | 0,036617129 | 6038 | 364 | 125 |
| GFB | *Oexaltata* | *Oapifera* | *Otenthredinifera* | 0,5685484 | 0 | 0 | 0,044276967 | 5873 | 389 | 107 |
| GIB | *Oexaltata* | *Oscolopax* | *Otenthredinifera* | 0,2229299 | 0,003984526 | 0,005312701 | 0,004985755 | 6863 | 96 | 61 |
| AFD | *Oinsectifera* | *Oapifera* | *Obombyliflora* | 0,3844252 | 0 | 0 | 0,060706617 | 4314 | 560 | 249 |
| AID | *Oinsectifera* | *Oscolopax* | *Obombyliflora* | 0,2045827 | 4,05E-07 | 9,86E-07 | 0,02380499 | 4640 | 368 | 243 |
| AGD | *Oinsectifera* | *Oexaltata* | *Obombyliflora* | 0,1527094 | 8,94E-05 | 0,000147247 | 0,017667173 | 4655 | 351 | 258 |
| IFD | *Oscolopax* | *Oapifera* | *Obombyliflora* | 0,4083885 | 0 | 0 | 0,028487835 | 6041 | 319 | 134 |
| GFD | *Oexaltata* | *Oapifera* | *Obombyliflora* | 0,4587738 | 0 | 0 | 0,034167848 | 5878 | 345 | 128 |
| GID | *Oexaltata* | *Oscolopax* | *Obombyliflora* | 0,2317881 | 0,003387398 | 0,00462669 | 0,004988597 | 6865 | 93 | 58 |
| AFE | *Oinsectifera* | *Oapifera* | *Olutea* | 0,3857868 | 0 | 0 | 0,059270813 | 4341 | 546 | 242 |
| AIE | *Oinsectifera* | *Oscolopax* | *Olutea* | 0,1844332 | 9,68E-06 | 2,08E-05 | 0,02076586 | 4658 | 350 | 241 |
| AGE | *Oinsectifera* | *Oexaltata* | *Olutea* | 0,1413969 | 0,000467823 | 0,000727725 | 0,01578547 | 4671 | 335 | 252 |
| IFE | *Oscolopax* | *Oapifera* | *Olutea* | 0,4592075 | 0 | 0 | 0,03042471 | 6046 | 313 | 116 |
| GFE | *Oexaltata* | *Oapifera* | *Olutea* | 0,5045045 | 0 | 0 | 0,03533123 | 5896 | 334 | 110 |
| GIE | *Oscolopax* | *Oexaltata* | *Olutea* | 0,1895425 | 0,01763845 | 0,021950071 | | 6886 | 91 | 62 |
| AFC | *Oinsectifera* | *Oapifera* | *Ospeculum* | 0,388203 | 0 | 0 | 0,055653884 | 4356 | 506 | 223 |
| AIC | *Oinsectifera* | *Oscolopax* | *Ospeculum* | 0,1956522 | 2,70E-06 | 6,05E-06 | 0,020646148 | 4679 | 330 | 222 |
| AGC | *Oinsectifera* | *Oexaltata* | *Ospeculum* | 0,1565836 | 0,000149215 | 0,000238744 | 0,016758713 | 4689 | 325 | 237 |
| IFC | *Oscolopax* | *Oapifera* | *Ospeculum* | 0,4635417 | 0 | 0 | 0,027575523 | 6071 | 281 | 103 |
| GFC | *Oexaltata* | *Oapifera* | *Ospeculum* | 0,4669927 | 0 | 0 | 0,030211958 | 5913 | 300 | 109 |
| GIC | *Oexaltata* | *Oscolopax* | *Ospeculum* | 0,129771 | 0,1442076 | 0,161512512 | | 6892 | 74 | 57 |
| CBD | *Ospeculum* | *Otenthredinifera* | *Obombyliflora* | 0,3738505 | 0 | 0 | 0,267525036 | 994 | 1718 | 783 |
| BEC | *Otenthredinifera* | *Olutea* | *Ospeculum* | 0,07692308 | 0,00289195 | 0,004261821 | 0,054388489 | 1882 | 891 | 702 |
| CED | *Ospeculum* | *Olutea* | *Obombyliflora* | 0,09287926 | 8,00E-05 | 0,000135758 | 0,057526366 | 1191 | 1059 | 879 |
| EBD | *Olutea* | *Otenthredinifera* | *Obombyliflora* | 0,4210526 | 0 | 0 | 0,223355705 | 1749 | 1404 | 572 |

**Table S3a.** Aphid results summary inferred using the majority backbone tree as a reference and *Himantoglossum* as the outgroup in every triplets.
For each triplet (($P_1$, $P_2$), $P_3$), the following are reported:
Number of trees kept by the preprocessing step (**n_trees**);
Estimated contributions of evolutionary scenarios: no-event (**$p_{none}$**), ILS (**$p_{ILS}$**) or GF (**$p_{GF}$**) with a (($P_1$, $P_2$), $P_3$) topology;
Estimated contributions of evolutionary scenarios: ILS (**$p_{ILSc}$**) or GF (**$p_{GFc}$**) with a conflicting topology;
Probability of ancient gene flow (**$p_a$**);
ILS imbalance (**$I_{ILS}$**) and associated *p* -values (***p* -value$_{ILS}$**) and corrected *p* -values;
Maximum log-likelihood for: model with gene flow (**lnL**), model without $P_1$-$P_2$ gene flow (**$lnL_{P_1P_2}$**), without $P_1$-$P_3$ gene flow (**$lnL_{P_1P_3}$**) and without $P_2$-$P_3$ gene flow (**$lnL_{P_2P_3}$**);
P-values of Likelihood ratio test between models with and without gene flow (***p* -value$_{P_1P_2}$**, ***p* -value$_{P_1P_3}$**, ***p* -value$_{P_2P_3}$**);
Differences in log-likelihood between models with and without gene flow (**$\Delta lnL_{P_1P_2}$**, **$\Delta lnL_{P_1P_3}$**, **$\Delta lnL_{P_2P_3}$**).

| Triplet ID | $P_1$ | $P_2$ | $P_3$ | n_trees | $p_{none}$ | $p_{ILS}$ | $p_{GF}$ | $p_{ILSc}$ | $p_{GFc}$ | $p_a$ | $I_{ILS}$ | *p* -value$_{ILS}$ | corrected *p* -value$_{ILS}$ | lnL | $lnL_{P_1P_2}$ | $LnL_{P_1P_3}$ | $lnL_{P_2P_3}$ | *P* -value$_{P_1P_2}$ | *P* -value$_{P_1P_3}$ | *P* -value$_{P_2P_3}$ | $\Delta lnL_{P_1P_2}$ | $\Delta lnL_{P_1P_3}$ | $\Delta lnL_{P_2P_3}$ |
|---|---|---|---|---|---|---|---|---|---|---|---|---|---|---|---|---|---|---|---|---|---|---|---|
| DBA | *Obombyliflora* | *Otenthredinifera* | *Oinsectifera* | 7305 | 0,331 | 0,064 | 0,375 | 0,116 | 0,115 | 0,834503 | 0,504 | 0,306657915 | 0,336722416 | -133931,847 | -139557,85 | -135649,284 | -135831,019 | 0 | 0 | 0 | 5626,003 | 1717,437 | 1899,172 |
| EBA | *Olutea* | *Otenthredinifera* | *Oinsectifera* | 7300 | 0,343 | 0,067 | 0,373 | 0,11 | 0,108 | 0,88938 | 0,522 | 0,090677999 | 0,123852877 | -133006,057 | -137385,15 | -134735,686 | -134969,202 | 0 | 0 | 0 | 4379,093 | 1729,629 | 1963,145 |
| CBA | *Ospeculum* | *Otenthredinifera* | *Oinsectifera* | 7310 | 0,375 | 0,067 | 0,307 | 0,117 | 0,134 | 0,90339 | 0,538 | 0,008719606 | 0,025699892 | -133511,619 | -137443,678 | -135231,134 | -135964,482 | 0 | 0 | 0 | 3932,059 | 1719,515 | 2452,863 |
| EDA | *Olutea* | *Obombyliflora* | *Oinsectifera* | 7306 | 0,363 | 0,068 | 0,303 | 0,123 | 0,143 | 0,893905 | 0,52 | 0,061484142 | 0,093057079 | -133963,164 | -137854,299 | -136251,876 | -136302,071 | 0 | 0 | 0 | 3891,135 | 2288,712 | 2338,907 |
| CDA | *Ospeculum* | *Obombyliflora* | *Oinsectifera* | 7313 | 0,363 | 0,072 | 0,29 | 0,133 | 0,142 | 0,903691 | 0,52 | 0,084451028 | 0,118231439 | -134927,923 | -138681,926 | -136854,49 | -137503,848 | 0 | 0 | 0 | 3754,003 | 1926,567 | 2575,925 |
| CEA | *Ospeculum* | *Olutea* | *Oinsectifera* | 7311 | 0,375 | 0,068 | 0,31 | 0,119 | 0,129 | 0,910112 | 0,502 | 0,445253744 | 0,461744623 | -133891,051 | -137608,412 | -135631,116 | -136223,013 | 0 | 0 | 0 | 3717,361 | 1740,065 | 2331,962 |
| FIA | *Oapifera* | *Oscolopax* | *Oinsectifera* | 7176 | 0,254 | 0,049 | 0,45 | 0,128 | 0,12 | 0,73907 | 0,609 | 1,51E-12 | 1,06E-11 | -132652,492 | -140155,656 | -134546,909 | -135475,358 | 0 | 0 | 0 | 7503,164 | 1894,417 | 2822,866 |
| FGA | *Oapifera* | *Oexaltata* | *Oinsectifera* | 7173 | 0,235 | 0,045 | 0,441 | 0,124 | 0,155 | 0,819593 | 0,592 | 2,52E-08 | 1,28E-07 | -132019,188 | -137335,322 | -134663,767 | -135516,071 | 0 | 0 | 0 | 5316,134 | 2644,579 | 3496,883 |
| IGA | *Oscolopax* | *Oexaltata* | *Oinsectifera* | 7150 | 0,241 | 0,037 | 0,592 | 0,081 | 0,049 | 0,523474 | 0,515 | 0,171250114 | 0,20404269 | -124944,912 | -135450,08 | -126046,058 | -126033,897 | 0 | 0 | 0 | 10505,168 | 1101,146 | 1088,985 |
| DBF | *Obombyliflora* | *Otenthredinifera* | *Oapifera* | 7293 | 0,339 | 0,07 | 0,329 | 0,114 | 0,148 | 0,832946 | 0,507 | 0,233344598 | 0,272235364 | -134247,339 | -139342,066 | -136449,89 | -137571,83 | 0 | 0 | 0 | 5094,727 | 2202,551 | 3324,491 |
| EBF | *Olutea* | *Otenthredinifera* | *Oapifera* | 7288 | 0,349 | 0,07 | 0,332 | 0,112 | 0,138 | 0,874283 | 0,532 | 0,023839072 | 0,047678143 | -133225,496 | -137186,882 | -135125,901 | -136620,426 | 0 | 0 | 0 | 3961,386 | 1900,405 | 3394,93 |
| CBF | *Ospeculum* | *Otenthredinifera* | *Oapifera* | 7299 | 0,376 | 0,069 | 0,269 | 0,117 | 0,169 | 0,884264 | 0,536 | 0,014263351 | 0,031949907 | -133455,547 | -136837,823 | -135353,464 | -137525,296 | 0 | 0 | 0 | 3382,276 | 1897,917 | 4069,749 |
| EDF | *Olutea* | *Obombyliflora* | *Oapifera* | 7297 | 0,362 | 0,07 | 0,272 | 0,122 | 0,174 | 0,889109 | 0,513 | 0,150479818 | 0,184307595 | -133917,852 | -137403,889 | -136745,187 | -137235,119 | 0 | 0 | 0 | 3486,037 | 2827,335 | 3317,267 |
| CDF | *Ospeculum* | *Obombyliflora* | *Oapifera* | 7301 | 0,365 | 0,071 | 0,258 | 0,127 | 0,178 | 0,896106 | 0,517 | 0,103931654 | 0,135352851 | -134372,596 | -137672,424 | -136718,27 | -137918,053 | 0 | 0 | 0 | 3299,828 | 2345,674 | 3545,457 |
| CEF | *Ospeculum* | *Olutea* | *Oapifera* | 7301 | 0,386 | 0,071 | 0,267 | 0,127 | 0,148 | 0,911095 | 0,5 | 0,490848922 | 0,490848922 | -133354,567 | -136641,16 | -135482,73 | -136258,887 | 0 | 0 | 0 | 3286,593 | 2128,163 | 2904,32 |
| GIF | *Oexaltata* | *Oscolopax* | *Oapifera* | 7034 | 0,203 | 0,026 | 0,459 | 0,087 | 0,225 | 0,579882 | 0,539 | 0,010784647 | 0,029036025 | -128785,912 | -137290,958 | -130957,725 | -134848,655 | 0 | 0 | 0 | 8505,046 | 2171,813 | 6062,743 |
| DBI | *Obombyliflora* | *Otenthredinifera* | *Oscolopax* | 7306 | 0,332 | 0,067 | 0,361 | 0,114 | 0,125 | 0,832557 | 0,504 | 0,265633277 | 0,300935698 | -134362,786 | -139720,399 | -136207,125 | -136835,122 | 0 | 0 | 0 | 5357,613 | 1844,339 | 2472,336 |
| EBI | *Olutea* | *Otenthredinifera* | *Oscolopax* | 7300 | 0,345 | 0,068 | 0,363 | 0,108 | 0,116 | 0,881337 | 0,522 | 0,077298839 | 0,110993205 | -133936,094 | -138181,159 | -135543,009 | -136364,536 | 0 | 0 | 0 | 4245,065 | 1606,915 | 2428,442 |
| CBI | *Ospeculum* | *Otenthredinifera* | *Oscolopax* | 7309 | 0,375 | 0,067 | 0,297 | 0,117 | 0,144 | 0,886614 | 0,532 | 0,019989216 | 0,041459114 | -134037,814 | -137729,249 | -135822,828 | -136956,184 | 0 | 0 | 0 | 3691,435 | 1785,014 | 2918,37 |
| EDI | *Olutea* | *Obombyliflora* | *Oscolopax* | 7311 | 0,365 | 0,07 | 0,294 | 0,12 | 0,151 | 0,886473 | 0,513 | 0,151395524 | 0,184307595 | -134966,655 | -138698,344 | -137391,569 | -137724,607 | 0 | 0 | 0 | 3731,689 | 2424,914 | 2757,952 |
| CDI | *Ospeculum* | *Obombyliflora* | *Oscolopax* | 7317 | 0,366 | 0,07 | 0,282 | 0,127 | 0,155 | 0,89412 | 0,5 | 0,397562052 | 0,420065564 | -135591,167 | -139196,844 | -137724,825 | -138499,561 | 0 | 0 | 0 | 3605,677 | 2133,658 | 2908,394 |
| CEI | *Ospeculum* | *Olutea* | *Oscolopax* | 7309 | 0,383 | 0,07 | 0,293 | 0,121 | 0,132 | 0,909699 | 0,514 | 0,132346301 | 0,168440747 | -134094,173 | -137645,164 | -135973,589 | -136495,886 | 0 | 0 | 0 | 3550,991 | 1879,416 | 2401,713 |
| DBG | *Obombyliflora* | *Otenthredinifera* | *Oexaltata* | 7308 | 0,34 | 0,071 | 0,354 | 0,117 | 0,119 | 0,839164 | 0,521 | 0,049270524 | 0,081151452 | -134470,069 | -139814,245 | -136377,468 | -136741,552 | 0 | 0 | 0 | 5344,176 | 1907,399 | 2271,483 |
| EBG | *Olutea* | *Otenthredinifera* | *Oexaltata* | 7304 | 0,349 | 0,07 | 0,359 | 0,11 | 0,112 | 0,883884 | 0,526 | 0,048008641 | 0,081151452 | -133915,088 | -138158,652 | -135478,098 | -136294,199 | 0 | 0 | 0 | 4243,564 | 1563,01 | 2379,111 |
| CBG | *Ospeculum* | *Otenthredinifera* | *Oexaltata* | 7311 | 0,379 | 0,068 | 0,295 | 0,116 | 0,141 | 0,886575 | 0,536 | 0,016561495 | 0,035670911 | -133981,416 | -137674,117 | -135937,062 | -136659,984 | 0 | 0 | 0 | 3692,701 | 1955,646 | 2678,568 |
| EDG | *Olutea* | *Obombyliflora* | *Oexaltata* | 7313 | 0,364 | 0,069 | 0,296 | 0,122 | 0,148 | 0,887031 | 0,506 | 0,313317087 | 0,337418401 | -134778,555 | -138612,532 | -137053,273 | -137510,642 | 0 | 0 | 0 | 3833,977 | 2274,718 | 2732,087 |
| CDG | *Ospeculum* | *Obombyliflora* | *Oexaltata* | 7317 | 0,366 | 0,071 | 0,283 | 0,125 | 0,155 | 0,89085 | 0,508 | 0,268692588 | 0,300935698 | -135428,119 | -139030,713 | -137633,447 | -138275,62 | 0 | 0 | 0 | 3602,594 | 2205,328 | 2847,501 |
| CEG | *Ospeculum* | *Olutea* | *Oexaltata* | 7311 | 0,385 | 0,071 | 0,293 | 0,12 | 0,132 | 0,903099 | 0,501 | 0,480857117 | 0,489599974 | -134260,599 | -137909,361 | -136319,82 | -136547,739 | 0 | 0 | 0 | 3648,762 | 2059,221 | 2287,14 |
| AFB | *Oinsectifera* | *Oapifera* | *Otenthredinifera* | 7303 | 0,268 | 0,065 | 0,383 | 0,136 | 0,148 | 0,828228 | 0,563 | 1,12E-05 | 4,84E-05 | -135888,783 | -141236,208 | -137269,338 | -139823,964 | 0 | 0 | 0 | 5347,425 | 1380,555 | 3935,181 |
| AIB | *Oinsectifera* | *Oscolopax* | *Otenthredinifera* | 7306 | 0,29 | 0,07 | 0,402 | 0,134 | 0,103 | 0,828626 | 0,532 | 0,013154263 | 0,03069328 | -136133,061 | -141497,805 | -137598,886 | -138317,764 | 0 | 0 | 0 | 5364,744 | 1465,825 | 2184,703 |
| AGB | *Oinsectifera* | *Oexaltata* | *Otenthredinifera* | 7306 | 0,296 | 0,073 | 0,395 | 0,138 | 0,097 | 0,83365 | 0,527 | 0,029406487 | 0,053121395 | -136011,8 | -141407,704 | -137503,255 | -137893,502 | 0 | 0 | 0 | 5395,904 | 1491,455 | 1881,702 |
| IFB | *Oscolopax* | *Oapifera* | *Otenthredinifera* | 7291 | 0,194 | 0,049 | 0,62 | 0,085 | 0,051 | 0,703314 | 0,636 | 1,14E-12 | 9,11E-12 | -132902,737 | -141577,278 | -133576,878 | -134671,898 | 0 | 3,64E-295 | 0 | 8674,541 | 674,141 | 1769,161 |
| GFB | *Oexaltata* | *Oapifera* | *Otenthredinifera* | 7293 | 0,182 | 0,045 | 0,625 | 0,088 | 0,06 | 0,723396 | 0,642 | 3,33E-16 | 6,22E-15 | -132257,981 | -139224,705 | -132911,401 | -134383,678 | 0 | 3,69E-286 | 0 | 6966,724 | 653,42 | 2125,697 |
| GIB | *Oexaltata* | *Oscolopax* | *Otenthredinifera* | 7294 | 0,227 | 0,038 | 0,678 | 0,045 | 0,012 | 0,494699 | 0,548 | 0,028365129 | 0,053121395 | -126192,46 | -138988,068 | -126414,317 | -126516,03 | 0 | 1,68E-98 | 9,36E-143 | 12795,608 | 221,857 | 323,57 |
| AFD | *Oinsectifera* | *Oapifera* | *Obombyliflora* | 7306 | 0,269 | 0,067 | 0,382 | 0,136 | 0,145 | 0,838079 | 0,546 | 0,000402587 | 0,001610347 | -137249,31 | -142766,609 | -138863,641 | -140557,601 | 0 | 0 | 0 | 5517,299 | 1614,331 | 3308,291 |
| AID | *Oinsectifera* | *Oscolopax* | *Obombyliflora* | 7308 | 0,288 | 0,071 | 0,404 | 0,13 | 0,107 | 0,82695 | 0,522 | 0,052271893 | 0,083635029 | -137455,422 | -142931,631 | -139129,078 | -139502,753 | 0 | 0 | 0 | 5476,209 | 1673,656 | 2047,331 |
| AGD | *Oinsectifera* | *Oexaltata* | *Obombyliflora* | 7308 | 0,292 | 0,074 | 0,4 | 0,132 | 0,103 | 0,827781 | 0,517 | 0,094293813 | 0,125725084 | -137194,261 | -142636,711 | -138874,419 | -139042,851 | 0 | 0 | 0 | 5442,45 | 1680,158 | 1848,59 |
| IFD | *Oscolopax* | *Oapifera* | *Obombyliflora* | 7293 | 0,193 | 0,05 | 0,624 | 0,093 | 0,041 | 0,704048 | 0,616 | 5,91E-11 | 3,67E-10 | -133779,407 | -141721,162 | -134391,59 | -134996,945 | 0 | 3,09E-268 | 0 | 7941,755 | 612,183 | 1217,538 |
| GFD | *Oexaltata* | *Oapifera* | *Obombyliflora* | 7298 | 0,182 | 0,044 | 0,627 | 0,091 | 0,056 | 0,720178 | 0,634 | 7,99E-15 | 8,95E-14 | -133179,997 | -139839,991 | -133952,743 | -134834,682 | 0 | 0 | 0 | 6659,994 | 772,746 | 1654,685 |
| GID | *Oexaltata* | *Oscolopax* | *Obombyliflora* | 7296 | 0,228 | 0,038 | 0,677 | 0,046 | 0,011 | 0,493937 | 0,574 | 0,004189812 | 0,014928361 | -127335,018 | -140150,634 | -127540,726 | -127622,526 | 0 | 1,80E-91 | 4,55E-127 | 12815,616 | 205,708 | 287,508 |
| AFE | *Oinsectifera* | *Oapifera* | *Olutea* | 7305 | 0,265 | 0,065 | 0,396 | 0,135 | 0,139 | 0,827722 | 0,576 | 1,77E-08 | 9,94E-08 | -136726,002 | -142259,531 | -138315,876 | -139691,197 | 0 | 0 | 0 | 5533,529 | 1589,874 | 2965,195 |
| AIE | *Oinsectifera* | *Oscolopax* | *Olutea* | 7308 | 0,287 | 0,072 | 0,41 | 0,128 | 0,104 | 0,825711 | 0,54 | 0,004265246 | 0,014928361 | -136974,701 | -142562,054 | -138695,293 | -138777,916 | 0 | 0 | 0 | 5587,353 | 1720,592 | 1803,215 |
| AGE | *Oinsectifera* | *Oexaltata* | *Olutea* | 7311 | 0,291 | 0,073 | 0,407 | 0,133 | 0,096 | 0,825674 | 0,534 | 0,01140701 | 0,029036025 | -137128,367 | -142654,497 | -138797,504 | -138706,861 | 0 | 0 | 0 | 5526,13 | 1669,137 | 1578,494 |
| IFE | *Oscolopax* | *Oapifera* | *Olutea* | 7295 | 0,198 | 0,053 | 0,62 | 0,093 | 0,036 | 0,701467 | 0,659 | 0 | 0 | -133370,347 | -141121,254 | -133812,145 | -134438,38 | 0 | 3,61E-194 | 0 | 7750,907 | 441,798 | 1068,033 |
| GFE | *Oexaltata* | *Oapifera* | *Olutea* | 7299 | 0,183 | 0,049 | 0,629 | 0,093 | 0,046 | 0,721305 | 0,648 | 1,67E-15 | 2,33E-14 | -132848,113 | -139095,564 | -133360,177 | -134299,673 | 0 | 1,02E-224 | 0 | 6247,451 | 512,064 | 1451,56 |
| GIE | *Oexaltata* | *Oscolopax* | *Olutea* | 7297 | 0,229 | 0,038 | 0,678 | 0,044 | 0,01 | 0,494303 | 0,537 | 0,057311193 | 0,089150744 | -126795,461 | -139662,582 | -126966,998 | -127024,616 | 0 | 1,37E-76 | 1,12E-101 | 12867,121 | 171,537 | 229,155 |
| AFC | *Oinsectifera* | *Oapifera* | *Ospeculum* | 7318 | 0,271 | 0,07 | 0,39 | 0,147 | 0,122 | 0,83901 | 0,569 | 1,80E-07 | 8,39E-07 | -137236,924 | -142880,03 | -138610,567 | -139825,225 | 0 | 0 | 0 | 5643,106 | 1373,643 | 2588,301 |
| AIC | *Oinsectifera* | *Oscolopax* | *Ospeculum* | 7320 | 0,283 | 0,071 | 0,418 | 0,134 | 0,095 | 0,824324 | 0,533 | 0,012587569 | 0,030647993 | -137345,355 | -142857,301 | -138746,928 | -139016,629 | 0 | 0 | 0 | 5511,946 | 1401,573 | 1671,274 |
| AGC | *Oinsectifera* | *Oexaltata* | *Ospeculum* | 7321 | 0,296 | 0,075 | 0,402 | 0,137 | 0,09 | 0,827306 | 0,527 | 0,029009157 | 0,053121395 | -137372,406 | -142880,929 | -138797,118 | -139027,854 | 0 | 0 | 0 | 5508,523 | 1424,712 | 1655,448 |
| IFC | *Oscolopax* | *Oapifera* | *Ospeculum* | 7306 | 0,204 | 0,056 | 0,617 | 0,096 | 0,028 | 0,699372 | 0,635 | 6,11E-14 | 5,70E-13 | -133605,598 | -140985,814 | -133991,515 | -134399,805 | 0 | 7,18E-170 | 0 | 7380,216 | 385,917 | 794,207 |
| GFC | *Oexaltata* | *Oapifera* | *Ospeculum* | 7308 | 0,18 | 0,048 | 0,636 | 0,095 | 0,041 | 0,696491 | 0,648 | 0 | 0 | -133182,346 | -139407,575 | -133784,572 | -134265,979 | 0 | 6,57E-264 | 0 | 6225,229 | 602,226 | 1083,633 |
| GIC | *Oexaltata* | *Oscolopax* | *Ospeculum* | 7307 | 0,233 | 0,04 | 0,674 | 0,044 | 0,009 | 0,491125 | 0,567 | 0,00706788 | 0,021988961 | -126927,074 | -139938,619 | -127155,242 | -127065,624 | 0 | 3,01E-101 | 3,22E-62 | 13011,545 | 228,168 | 138,55 |
| CBD | *Ospeculum* | *Otenthredinifera* | *Obombyliflora* | 7220 | 0,209 | 0,041 | 0,057 | 0,147 | 0,546 | 0,877038 | 0,518 | 0,074307822 | 0,109506263 | -122919,688 | -123828,002 | -127074,006 | -131437,717 | 0 | 0 | 0 | 908,314 | 4154,318 | 8518,029 |
| BEC | *Otenthredinifera* | *Olutea* | *Ospeculum* | 7203 | 0,27 | 0,046 | 0,152 | 0,187 | 0,345 | 0,917909 | 0,526 | 0,01135474 | 0,029036025 | -121772,345 | -123570,869 | -125321,807 | -125338,437 | 0 | 0 | 0 | 1798,492 | 3549,43 | 3566,06 |
| CED | *Ospeculum* | *Olutea* | *Obombyliflora* | 7218 | 0,243 | 0,043 | 0,081 | 0,17 | 0,463 | 0,922887 | 0,53 | 0,006609483 | 0,021772415 | -122637,411 | -123715,566 | -127020,644 | -127707,126 | 0 | 0 | 0 | 1078,155 | 4383,233 | 5069,715 |
| EBD | *Olutea* | *Otenthredinifera* | *Obombyliflora* | 7205 | 0,267 | 0,049 | 0,116 | 0,174 | 0,394 | 0,894993 | 0,519 | 0,048884768 | 0,081151452 | -122371,018 | -123868,702 | -125124,565 | -128809,092 | 0 | 0 | 0 | 1497,684 | 2753,547 | 6438,074 |

**Table S3b.** Alternate Aphid results for the 8 triplets testing for gene flow within clades 1 and 2.
*O. speculum* was used as outgroup in triplets FIG, FGA, IGA, GIF.
*O. insectifera* was used as outgroup for triplets CBD, BEC, CED, EBD.
For each triplet (($P_1$, $P_2$), $P_3$), the following are reported:
Number of trees kept by the preprocessing step (**n_trees**);
Estimated contributions of evolutionary scenarios: no-event (**$p_{none}$**), ILS (**$p_{ILS}$**) or GF (**$p_{GF}$**) with a (($P_1$, $P_2$), $P_3$) topology;
Estimated contributions of evolutionary scenarios: ILS (**$p_{ILSc}$**) or GF (**$p_{GFc}$**) with a conflicting topology;
Probability of ancient gene flow (**$p_a$**);
ILS imbalance (**$I_{ILS}$**) and associated *p* -values (***p* -value$_{ILS}$**) and corrected *p* -values;
Maximum log-likelihood for: model with gene flow (**lnL**), model without $P_1$-$P_2$ gene flow (**$lnL_{P_1P_2}$**), without $P_1$-$P_3$ gene flow (**$lnL_{P_1P_3}$**) and without $P_2$-$P_3$ gene flow (**$lnL_{P_2P_3}$**);
P-values of Likelihood ratio test between models with and without gene flow (***p* -value$_{P_1P_2}$**, ***p* -value$_{P_1P_3}$**, ***p* -value$_{P_2P_3}$**);
Differences in log-likelihood between models with and without gene flow (**$\Delta lnL_{P_1P_2}$**, **$\Delta lnL_{P_1P_3}$**, **$\Delta lnL_{P_2P_3}$**).

| Triplet ID | $P_1$ | $P_2$ | $P_3$ | n_trees | $p_{none}$ | $p_{ILS}$ | $p_{GF}$ | $p_{ILSc}$ | $p_{GFc}$ | $p_a$ | $I_{ILS}$ | *p* -value$_{ILS}$ | corrected *p* -value$_{ILS}$ | lnL | $lnL_{P_1P_2}$ | $LnL_{P_1P_3}$ | $lnL_{P_2P_3}$ | *P* -value$_{P_1P_2}$ | *P* -value$_{P_1P_3}$ | *P* -value$_{P_2P_3}$ | $\Delta lnL_{P_1P_2}$ | $\Delta lnL_{P_1P_3}$ | $\Delta lnL_{P_2P_3}$ |
|---|---|---|---|---|---|---|---|---|---|---|---|---|---|---|---|---|---|---|---|---|---|---|---|
| FIA | *Oapifera* | *Oscolopax* | *Oinsectifera* | 4814 | 0,348 | 0,052 | 0,418 | 0,108 | 0,074 | 0,754986 | 0,548 | 0,007890625 | 0,022503235 | -72848,738 | -76942,505 | -73647,436 | -73895,523 | 0 | 0 | 0 | 4093,767 | 798,698 | 1046,785 |
| FGA | *Oapifera* | *Oexaltata* | *Oinsectifera* | 4777 | 0,314 | 0,046 | 0,429 | 0,108 | 0,103 | 0,850204 | 0,55 | 0,006961796 | 0,022503235 | -72171,666 | -74684,752 | -73251,991 | -73433,146 | 0 | 0 | 0 | 2513,086 | 1080,325 | 1261,48 |
| IGA | *Oscolopax* | *Oexaltata* | *Oinsectifera* | 5080 | 0,276 | 0,035 | 0,586 | 0,069 | 0,034 | 0,522236 | 0,503 | 0,378080355 | 0,407163459 | -74154,932 | -80465,502 | -74605,374 | -74650,716 | 0 | 6,30E-198 | 1,22E-217 | 6310,57 | 450,442 | 495,784 |
| GIF | *Oexaltata* | *Oscolopax* | *Oapifera* | 5689 | 0,199 | 0,034 | 0,467 | 0,114 | 0,186 | 0,592837 | 0,543 | 0,009279682 | 0,024745818 | -85548,82 | -90665,934 | -86386,405 | -89138,344 | 0 | 0 | 0 | 5117,114 | 837,585 | 3589,524 |
| CBD | *Ospeculum* | *Otenthredinifera* | *Obombyliflora* | 4619 | 0,183 | 0,039 | 0,074 | 0,193 | 0,51 | 0,873902 | 0,513 | 0,101575649 | 0,13674058 | -69225,041 | -70009,612 | -70543,137 | -72715,762 | 0 | 0 | 0 | 784,571 | 1318,096 | 3490,721 |
| BEC | *Otenthredinifera* | *Olutea* | *Ospeculum* | 4716 | 0,268 | 0,052 | 0,189 | 0,209 | 0,282 | 0,92552 | 0,543 | 0,000980051 | 0,004221757 | -69576,672 | -70801,143 | -70930,791 | -71051,141 | 0 | 0 | 0 | 1224,471 | 1354,119 | 1474,469 |
| CED | *Ospeculum* | *Olutea* | *Obombyliflora* | 4524 | 0,214 | 0,046 | 0,123 | 0,208 | 0,409 | 0,929249 | 0,511 | 0,201229117 | 0,244974577 | -66989,821 | -67778,11 | -68274,645 | -68565,88 | 0 | 0 | 0 | 788,289 | 1284,824 | 1576,059 |
| EBD | *Olutea* | *Otenthredinifera* | *Obombyliflora* | 4724 | 0,264 | 0,054 | 0,135 | 0,203 | 0,344 | 0,894188 | 0,534 | 0,007510521 | 0,022503235 | -69668,234 | -70620,502 | -70699,343 | -72568,354 | 0 | 0 | 0 | 952,268 | 1031,109 | 2900,12 |

**Table S4.** Detailed Aphid results for triplets $((P_1,P_2),P_3)$ supporting the three major gene flow events:
from *O.tenthredinifera* to *O.apifera* (triplets DBF, EBF, CBF);
from *O.apifera* to *O.scolopax* (triplet GIF);
from *O.bombyliflora* to *O.tenthredinifera* (triplets CBD, EBD).
In all triplets, major gene flow corresponds to gene flow between the $P_2$ and $P_3$ taxa, with $p_{GFc}$ the proportion of introgressed genes associated.
Gene flow timing is indicated using 2 or 10 timing categories, defined relative to $P_1$-$P_2$ speciation time, with for example:
**1:** probability that gene flow occurred at the speciation time;
**0.5:** probability that gene flow occurred twice as recently;
**0.1:** probability that gene flow occurred ten time more recently.

| | | 2 timing categories | | 10 timing categories | | | | | | | | | |
|---|---|---|---|---|---|---|---|---|---|---|---|---|---|
| Triplet ID | $p_{GFc}$ | 1 | 0,5 | 1 | 0,9 | 0,8 | 0,7 | 0,6 | 0,5 | 0,4 | 0,3 | 0,2 | 0,1 |
| DBF | 0,086 | 0,867 | 0,133 | 0,776 | 0,031 | 0,038 | 0,046 | 0,043 | 0,028 | 0,017 | 0,011 | 0,007 | 0,004 |
| EBF | 0,085 | 0,871 | 0,129 | 0,808 | 0,021 | 0,028 | 0,037 | 0,037 | 0,029 | 0,02 | 0,011 | 0,006 | 0,003 |
| CBF | 0,108 | 0,863 | 0,137 | 0,79 | 0,023 | 0,034 | 0,043 | 0,042 | 0,03 | 0,018 | 0,009 | 0,007 | 0,003 |
| GIF | 0,161 | 0,678 | 0,322 | 0,56 | 0,056 | 0,048 | 0,046 | 0,045 | 0,044 | 0,044 | 0,047 | 0,05 | 0,06 |
| CBD | 0,365 | 0,861 | 0,139 | 0,784 | 0,025 | 0,035 | 0,043 | 0,04 | 0,031 | 0,021 | 0,013 | 0,006 | 0,002 |
| EBD | 0,276 | 0,885 | 0,115 | 0,831 | 0,019 | 0,025 | 0,031 | 0,031 | 0,025 | 0,018 | 0,013 | 0,007 | 0,003 |

**Table S5.** Function and genomic location (through Gene_ID) on the *Ophrys* reference genome from Russo et al. (2024) of the 58 Ophrys flower phenotype key genes identified in our dataset.
For each key gene showing evidence of one of the three major gene flow events (based on gene tree topology) Aphid results are reported:
Genes are classified by Aphid as originating from gene flow (GF), incomplete lineage sorting (ILS), speciation (no-event),
or filtered during Aphid pre-processing (Na).
apifera-scolopax: *O.apifera* to *O.scolopax* gene flow
tenthred-apifera: *O.tenthredinifera* to *O.apifera* gene flow
Bomby-tenthred: *O.tenthredinifera* putative hybrid origin, genes grouping *O.bombyliflora* and *O.tenthredinifera*
Lutea-tenthred: *O.tenthredinifera* putative hybrid origin, genes grouping *O.lutea* and *O.tenthredinifera*

| Gene_ID | Function | apifera-scolopax | tenthred-apifera | bomby-tenthred | lutea-tenthred |
|---|---|---|---|---|---|
| | **Anthocyanin biosynthesis** | | | | |
| Osph3G27250.1 | ANS1 | - | - | - | - |
| Osph15G18650.1 | CHS3 | - | - | GF | - |
| Osph5G19490.1 | F3H1 | - | - | - | - |
| Osph4G56570.1 | - | - | - | - | - |
| | **Carotenoid biosynthesis** | | | | |
| Osph2G11980.1 | ABA2H1 | - | - | - | No-event |
| Osph20G10110.1 | CHY1 | - | - | GF | - |
| Osph18G13090.1 | NCED3 | - | - | - | - |
| Osph219G10330.1 | PDS1 | - | - | - | No-event |
| Osph91G10050.2 | PSY1 | - | - | - | - |
| Osph10G16730.1 | VDE1 | GF | - | GF | - |
| Osph68G11190.1 | ZEP1 | - | - | GF | - |
| Osph5G39630.1 | - | - | - | - | No-event |
| Osph5G48260.1 | CYP707A1 | - | Na | - | - |
| Osph14G43970 | PSY1 | - | - | - | No-event |
| | **Fatty acid, wax and hydrocarbon biosynthesis** | | | | |
| Osph16G32630.1 | DGAT3 | - | - | - | - |
| Osph56G12460.2 | DGAT4 | - | - | - | - |
| Osph116G10900.1 | DGAT5 | - | - | GF | - |
| Osph9G12320.4 | ECR3 | - | - | - | - |
| Osph5G55210.3 | FAT5 | - | - | GF | - |
| Osph3G76480.2 | FCT1 | ILS | - | GF | - |
| Osph81G10950.1 | KASi2 | GF | - | - | - |
| Osph97G10840.1 | KASi3 | - | - | GF | - |
| Osph50G12130.1 | LGPAT6 | - | - | - | No-event |
| Osph103G10680.1 | LGPAT7 | - | - | - | - |
| Osph282G10060.1 | LGPAT8 | GF | - | GF | - |
| Osph13G15790.1 | - | - | - | - | - |
| Osph16G37930.1 | - | - | - | ILS | - |
| Osph3G74440.1 | - | - | Na | - | - |
| Osph7G47300 | - | - | - | - | No-event |
| Osph8G26560 | - | - | - | - | No-event |
| Osph8G43810 | - | GF | - | - | - |
| Osph13G10580 | ECI1 | - | - | - | No-event |
| | **VLCFA and hydrocarbon biosynthesis** | | | | |
| Osph5G20630.1 | ACP3 | - | - | GF | - |
| Osph152G10450.1 | CER3H1 | - | - | GF | - |
| Osph21G10840.2 | CYTB5H2 | - | - | ILS | - |
| Osph1G68690.1 | EAR1 | - | - | GF | - |
| Osph4G30620.1 | ECR2 | - | - | - | - |
| Osph1G80960.1 | EMR2Ψ | GF | - | - | - |
| Osph5G32390.1 | FAD2Ψ | - | - | - | No-event |
| Osph33G11900.1 | FAD4 | - | - | - | No-event |
| Osph219G10030.1 | FAT2 | - | - | - | No-event |
| Osph17G26180.1 | HAD1 | - | - | - | GF |
| Osph6G35180.1 | HCD1 | - | - | - | - |
| Osph15G26270.1 | KAR1Ψ | - | - | GF | - |
| Osph9G29660.1 | KCS5 | - | - | - | - |
| Osph3G50760.1 | KCS9 | - | - | - | No-event |
| Osph30G10690.1 | KCS10 | - | - | Na | - |
| Osph7G31430.1 | LACS1 | - | - | - | - |
| Osph17G26020.1 | LACS2 | - | - | - | No-event |
| | **Transcription factors of interest** | | | | |
| Osph1456G10010.2 | DEF2H1 | - | - | - | - |
| Osph3G26320.1 | DEF3H1 | - | - | - | ILS |
| Osph18G15220.1 | DEF4H1Ψ | - | - | - | - |
| Osph17G28910.1 | GL3H1Ψ | - | - | - | - |
| Osph1G68550.1 | SPL9H2 | - | - | GF | - |
| Osph6G51030.1 | SPY1 | - | - | - | GF |
| Osph10G10780.1 | MAS | - | - | - | - |
| Osph36G10370.1 | MADS | - | - | - | No-event |
| | **Other** | | | | |
| Osph2G62780 | VPS45 | GF | - | GF | - |

**Table S6.** Summary of Aphid results for 10 triplets.
**% removed:** percentage of genes removed during Aphid pre-processing.
**% GF:** percentage of genes identified as originating from GF among the remaining genes.
Statistics were calculated for three datasets defined by the correlation coefficient ($R^2$) between branch lengths and those of the concatenation-based tree.
**$R^2 > 0.8$:** final dataset of 7,821 genes with $R^2$ greater than 0.8.
**$0.8 > R^2 > 0.5$:** 141 genes removed by the Branch Length Comparison method, with $R^2$ between 0.8 and 0.5.
**$R^2 < 0.5$:** 65 genes removed by the Branch Length Comparison method, with $R^2$ below 0.5.

| | $R^2 > 0.8$ | | $0.8 > R^2 > 0.5$ | | $R^2 < 0.5$ | |
|---|---|---|---|---|---|---|
| Triplets | % removed | % GF | % removed | % GF | % removed | % GF |
| DBA | 6,6 | 48,7 | 20,6 | 44,6 | 55,4 | 44,8 |
| IGA | 35 | 62,3 | 72,3 | 66,7 | 78,5 | 85,7 |
| DBF | 6,8 | 47,3 | 21,3 | 50,5 | 53,8 | 26,7 |
| CDF | 6,6 | 42,8 | 21,3 | 45,9 | 50,8 | 34,4 |
| GIF | 27,2 | 65,6 | 65,2 | 55,1 | 69,2 | 80 |
| CEI | 6,5 | 41,7 | 17 | 46,2 | 40 | 30,8 |
| GIB | 6,7 | 69,7 | 17,7 | 62,9 | 50,8 | 62,5 |
| AGC | 6,4 | 49,7 | 17,7 | 46,6 | 47,7 | 35,3 |
| CBD | 41 | 57,3 | 76,6 | 57,6 | 78,5 | 71,4 |
| EBD | 39,7 | 45,9 | 75,2 | 57,1 | 76,9 | 60 |

**Table S7.** Estimation of heterozygosity rate for each sequenced individual.

| Species | Heterozygosity rate (%) |
|---|---|
| *O. insectifera* | 10,9 |
| *O. tenthredinifera* | 11,1 |
| *O. bombyliflora* | 10,5 |
| *O. speculum* | 9,1 |
| *O. lutea* | 11,4 |
| *O. apifera* | 10,3 |
| *O. exaltata* | 11,7 |
| *O. scolopax* | 10,3 |
| *H. hircinum* | 4,2 |
| *H. robertianum* | 5,6 |
